# Cryo-EM Structure of the Fork Protection Complex Bound to CMG at a Replication Fork

**DOI:** 10.1101/2019.12.18.880690

**Authors:** Domagoj Baretić, Michael Jenkyn-Bedford, Valentina Aria, Giuseppe Cannone, Mark Skehel, Joseph T.P. Yeeles

## Abstract

The eukaryotic replisome, organized around the Cdc45-MCM-GINS (CMG) helicase, orchestrates chromosome replication. Multiple factors associate directly with CMG including Ctf4 and the heterotrimeric fork protection complex (Csm3/Tof1 and Mrc1), that have important roles including aiding normal replication rates and stabilizing stalled forks. How these proteins interface with CMG to execute these functions is poorly understood. Here we present 3-3.5 Å resolution cryo-EM structures comprising CMG, Ctf4, Csm3/Tof1 and Mrc1 at a replication fork. The structures provide high-resolution views of CMG:DNA interactions, revealing the mechanism of strand separation. Furthermore, they illustrate the topology of Mrc1 in the replisome and show Csm3/Tof1 ‘grips’ duplex DNA ahead of CMG via a network of interactions that are important for efficient replication fork pausing. Our work reveals how four highly conserved replisome components collaborate with CMG to facilitate replisome progression and maintain genome stability.

---

Replication of eukaryotic genomes is initiated when double-hexameric minichromosome maintenance (MCM) complexes are activated to form two CMG helicases (Douglas et al., 2018; Yeeles et al., 2015). MCM is a two-tiered ring composed of six related subunits (Mcm2-7), with one tier of amino-terminal domains (N-tier), and a second tier of carboxy-terminal AAA+ domains that power ATP-dependent DNA unwinding (C-tier) (Bell and Botchan, 2013). Cdc45 and GINS are loaded onto MCM by the combined action of multiple ‘firing factors’ where they stabilize the N-tier through interactions with Mcm2, 3 and 5 (Costa et al., 2011; Yuan et al., 2016). CMG travels in an N-tier first orientation (Douglas et al., 2018; Georgescu et al., 2017), unwinding DNA by translocating 3′ – 5′ along the leading-strand template which is threaded through the central channel of MCM while the lagging-strand template is excluded (Fu et al., 2011). Translocation is proposed to occur via a non-symmetric rotary mechanism with ATP binding promoting single-stranded DNA (ssDNA) engagement by loops in the MCM C-tier (Eickhoff et al., 2019). Despite the importance of these ssDNA contacts, they are yet to be observed in CMG structures at high resolution. Moreover, the mechanism by which ssDNA translocation is coupled to strand separation is unresolved.

We previously showed that robust DNA synthesis in a reconstituted system requires only the proteins necessary for CMG activation, together with primase and DNA polymerases (Yeeles et al., 2015; Yeeles et al., 2017). However, these minimal replisomes exhibited partial functionality due to an absence of critical accessory proteins (Yeeles et al., 2017), many of which are essential for proper replication fork progression and maintenance of genome stability. These include Ctf4 (And-1), Mrc1 (Claspin) and Csm3/Tof1 (Tipin/Timeless) that were identified as components of replisome progression complexes - stable CMG-containing complexes isolated from S-phase yeast cells (Gambus et al., 2006). Ctf4 is a trimeric hub that recruits proteins involved in sister chromatid cohesion, ribosomal DNA maintenance, parental histone transfer and gene silencing (Evrin et al., 2018; Gan et al., 2018; Samora et al., 2016; Simon et al., 2014; Villa et al., 2016). Csm3/Tof1 and Mrc1 are key replisome modulators, collectively termed the fork protection complex, that intimately associate with one another, both physically and functionally (Bando et al., 2009; Katou et al., 2003). They are essential for normal replication rates (Petermann et al., 2008; Somyajit et al., 2017; Szyjka et al., 2005; Tourriere et al., 2005; Yeeles et al., 2017), maintaining coupling of DNA synthesis to CMG in response to hydroxyurea (HU) (Katou et al., 2003), limiting trinucleotide repeat instability (Gellon et al., 2019), and fully activating the S-phase checkpoint (Alcasabas et al., 2001; Foss, 2001).

Currently, very little is known about how these functions are accomplished. Understanding their mechanisms has been hindered by a lack of structures for these proteins in the context of a replisome. In fact, there are no structures of Mrc1 nor Csm3, and only an incomplete structure of the human Tof1 orthologue – Timeless – in isolation (Holzer et al., 2017). To address these issues, we have determined the cryo-EM structure of a 1.4 MDa complex comprising CMG, Ctf4 and the fork protection complex at a replication fork.

## Cryo-EM structure of Csm3/Tof1, Mrc1 and Ctf4 in complex with CMG at a replication fork

To assemble complexes for cryo-EM we incubated *S. cerevisiae* CMG with fork DNA, Ctf4, Csm3/Tof1, Mrc1 and the non-hydrolysable ATP analogue adenylyl-imidodiphosphate (AMP-PNP) (Figure S1A). Analysis of complex formation over glycerol gradients revealed Csm3/Tof1, Mrc1 and Ctf4 co-sedimenting with CMG (Figure 1A and Figure S1B). Samples for cryo-EM were prepared following glycerol gradient fixation (Kastner et al., 2008) yielding 3D reconstructions that enabled model building of CMG, the homotrimeric Ctf4 C-terminus, ∼900 residues of the Csm3/Tof1 heterodimer and fork DNA (Figure 1B,C, Figure S1 and S2, and Tables S1 and S2). In addition to assembling complexes by reconstitution, we also determined cryo-EM reconstructions of the same complex prepared following co-overexpression of all 15 proteins in *S. cerevisiae* (Figure S3). Figure S3H shows there are no major differences in the architecture of Csm3/Tof1 and Ctf4 bound to CMG between the different assembly methods.

**Figure 1.**
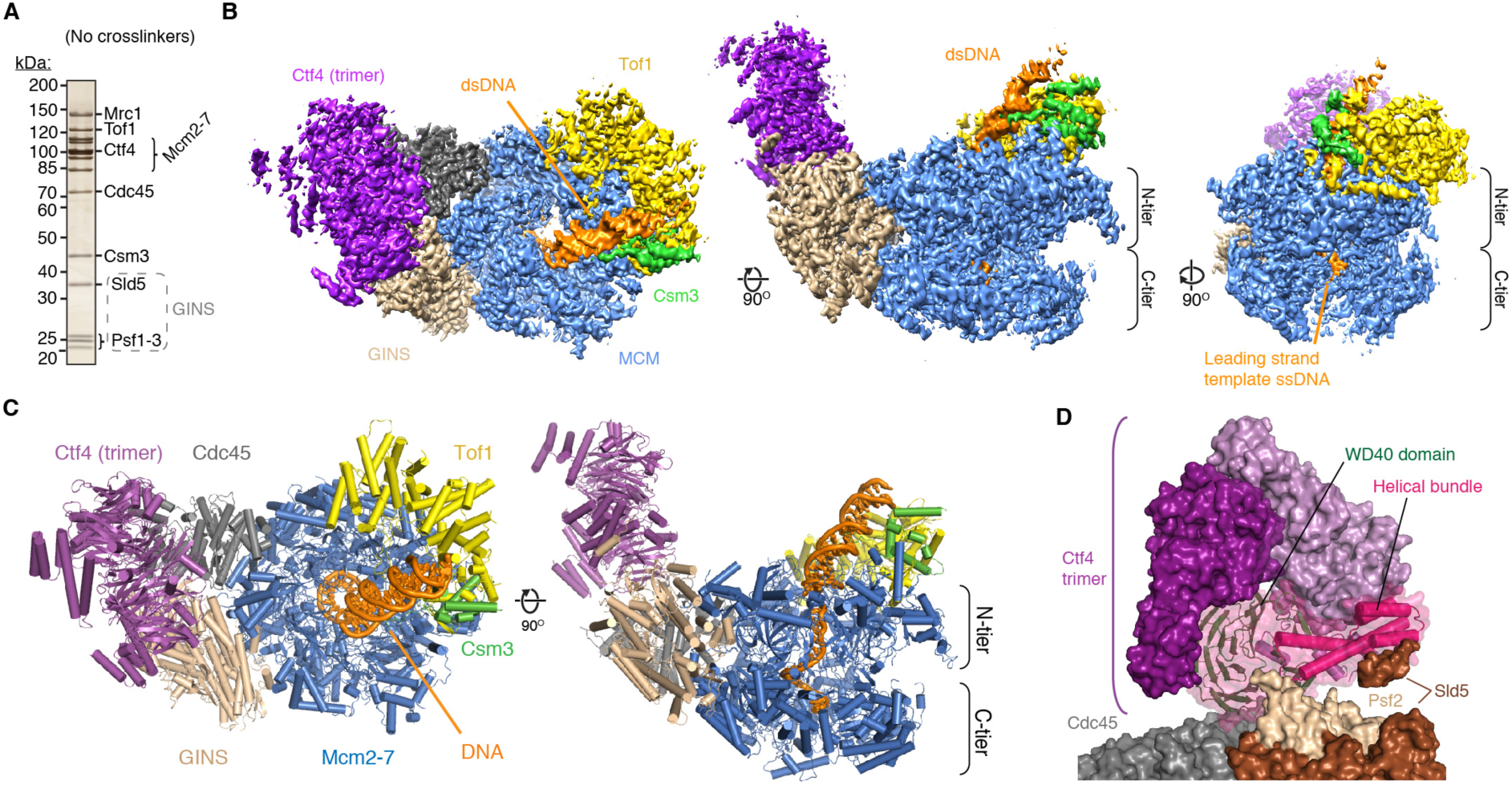
Structure of CMG bound to Csm3/Tof1, Mrc1, Ctf4 and a DNA fork. (A) Silver-stained SDS-PAGE of a representative glycerol gradient fraction of non-crosslinked sample (Fraction 11, Figure S1B) equivalent to fractions used for cryo-EM. (B, C) Cryo-EM density map (B) and corresponding atomic model (C) of a complex assembled as in Figure S1A observed in conformation 1. (D) Model of the Ctf4 trimer bound across the Cdc45-GINS interface of CMG, rendered as a surface. The Ctf4 monomer mediating the interaction with CMG is rendered also as a cartoon.

Overall the complex displays a ‘horseshoe-like’ configuration with the discoidal Ctf4 trimer sitting across the GINS:Cdc45 interface, whereas Csm3/Tof1 is located on the N-tier face of MCM (Figure 1B-D). Two turns of double-stranded DNA (dsDNA) protrude from the N-tier of the MCM central channel, tilted from the vertical axis of the channel towards Csm3/Tof1, where it contacts both proteins. We observe two major conformations for the MCM C-tier, termed conformation 1 and 2, although there are no major differences to the MCM N-tier or other subunits between the two conformations (Figure S1H-J, L and Movie S1).

## Structure of Ctf4 in the replisome

Within our structure the region of Ctf4 we resolve is very similar to crystal structures of isolated Ctf4 and And-1 C-terminal domains (Figure S4A) and a recent cryo-EM structure of CMG-Ctf4 lacking DNA (Yuan et al., 2019), indicating that neither the fork protection complex nor the replication fork alter its position in the replisome. The trimer is rigidly bound, sitting side-on across the interface between Cdc45 and the GINS subunit Psf2, with the helical domains facing away from the replication fork (Figure 1C,D and Figure S1L [left]). One Ctf4 monomer mediates the interaction, burying 520 Å^2^ of Cdc45 and 790 Å^2^ of Psf2, primarily involving two blades of its WD40 domain (Figure S4B-F). We do not observe clear density for the ∼400-residue (1-383) Ctf4 N-terminus predicted to contain a WD40-like domain (Guan et al., 2017), consistent with it being loosely connected to the C-terminus (Simon et al., 2014) and not stabilized by DNA or the fork protection complex in our structure. Because dsDNA is bent away from Ctf4 by Csm3/Tof1 at the front of the replisome, partner proteins that dock on to the replisome via Ctf4 may be located a considerable distance from the parental DNA duplex approaching the fork junction. This may be of particular significance for Pol α, which is involved in parental H3-H4 transfer to the lagging strand in conjunction with the N-terminal extension (NTE) of Mcm2 (Gan et al., 2018) (Figure S4G).

## Mrc1 spans one side of the replisome

The location of Csm3/Tof1 ahead of the replication fork has implications for Mrc1 topology in the replisome: in addition to binding Csm3/Tof1 (Bando et al., 2009; Lewis et al., 2017), Mrc1 associates with the Mcm6 winged-helix and Pol ε (Komata et al., 2009; Lou et al., 2008), both of which travel behind CMG (Goswami et al., 2018; Sun et al., 2015). Mrc1 might therefore stretch from the front to the rear of the replisome. Accordingly, although cryo-EM density accounting for a significant portion of Mrc1 (125 kDa) is not present in our initial reconstructions (Figure 1B and C), we observe disconnected helical densities across Tof1, Mcm6, Mcm2 and Cdc45 in samples prepared both by reconstitution and co-expression, whose sequence identity cannot not be assigned, but that we hypothesized could be contributed by Mrc1 (Figure 2A and Figure S5A). Cross-linking mass spectrometry supported this hypothesis: the N-terminus of Mrc1 crosslinked to Tof1, a cluster of cross-links involving the Mrc1 mid-region was observed at the C-tier by Mcm6/Mcm2, and several cross-links involving more C-terminal regions of Mrc1 were detected on Cdc45 and Ctf4 (Figure 2A and B and Table S3). Moreover, further processing involving signal subtraction and focused classification revealed larger regions of lower-resolution density in the vicinity of sites on CMG that crosslinked to Mrc1 (Figure S5B-D). Collectively, these data indicate Mrc1 contacts CMG in multiple locations across one side of the complex (Figure 2C), extending from its N-terminal association with Tof1 to its C-terminus located in the vicinity of Cdc45, which is also close to the binding site for the C-terminal non-catalytic domain of Pol ε with which it is proposed to interact (Goswami et al., 2018; Lou et al., 2008).

**Figure 2.**
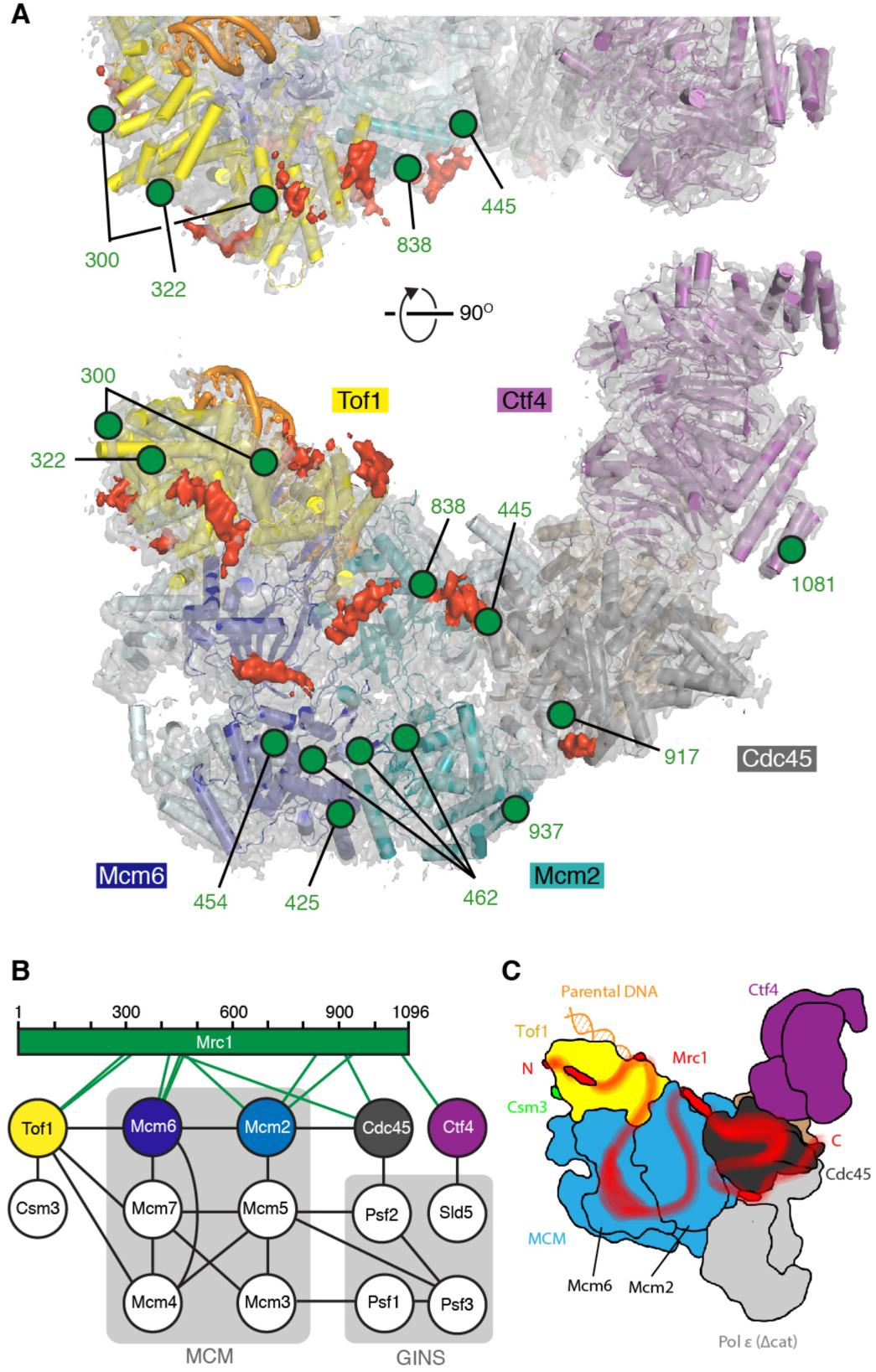
Position of Mrc1 in the eukaryotic replisome. (A) Cryo-EM density map with corresponding model for a reconstituted sample, with regions of unassigned density colored red. The positions of residues observed to crosslink with Mrc1 in cross-linking mass spectrometry (XL-MS) experiments performed on a co-expressed sample are indicated by green circles; green labels indicate which Mrc1 residues crosslink to these sites. (B) Summary of XL-MS for a co-expressed replisome sub-complex (see Table S3 for details of all inter- and intra-subunit cross-links). Lines indicate inter-subunit crosslinks. (C) Schematic of how Mrc1 might bind across one side of the eukaryotic replisome. The position of non-catalytic domains of Pol *ε* as previously determined by cryo-EM (Goswami et al., 2018) are shown.

We also note the positioning of Mrc1 in our structures, together with Tof1 and Ctf4, display considerable overlap with putative Mcm10 binding sites on CMG (Mayle et al., 2019), and therefore it is possible Mcm10 does not remain associated with CMG after its activation once Ctf4 and Mrc1 have been incorporated into the replisome.

## High-resolution details of CMG-ssDNA interactions in the MCM AAA+ C-tier

Despite the presence of the fork protection complex in our structure, we observe two C-tier configurations similar to two conformational states previously determined at lower resolution for isolated CMG in the presence of ATP (Eickhoff et al., 2019) (conformations 1 and 2, see Figure S1H-J,L and Movie S1). These two states were reported to represent CMG translocation intermediates, enabling CMG:ssDNA contacts made during translocation to be inferred from our structures. Conformations 1 and 2 differ in ATPase site occupancy and single-stranded DNA (ssDNA) engagement by the presensor-1 (PS1) hairpin and helix-2 (H2)/helix-2 insertion loop (H2I) that protrude into the MCM central channel (Figure 3A,B). In both conformations MCM subunits contact backbone phosphates using four highly conserved residues; every other phosphate is coordinated by both a PS1 lysine and the backbone amide of an H2I valine/isoleucine from a single MCM, while each remaining phosphate is bound by an H2 serine and PS1 alanine from two neighboring subunits (Figure 3C,D and Figure S6 and S7A-C). This establishes a highly repetitive arrangement whereby every phosphate pair is coordinated in an equivalent manner, as is observed in homohexameric archaeal MCM (Meagher et al., 2019) (Figure S7D [far-left]). Although the precise ssDNA contacts and number of phosphates bound per subunit varies amongst diverse hexameric helicases, the formation of repetitive phosphate backbone contacts is universal (Figure S7D).

**Figure 3.**
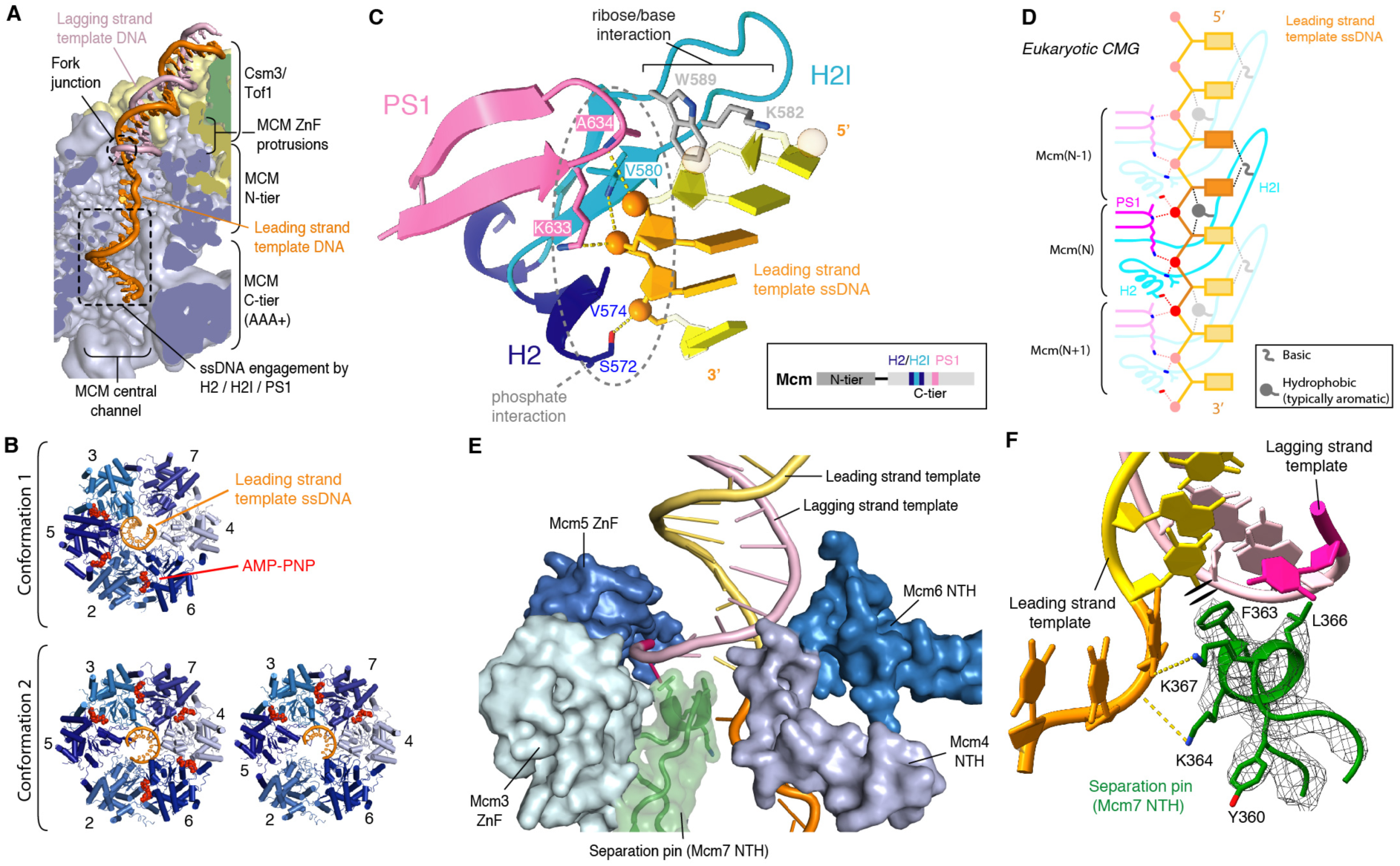
Interaction of eukaryotic CMG helicase with fork DNA. (A) Cut-away showing path of DNA approaching and traversing the MCM central channel. (B) Comparison of MCM C-tier between conformations 1 and 2 (subclasses bound to five or three AMP-PNP molecules shown). (C) Individual MCM ssDNA-binding motif (Mcm2 shown). Three phosphates contacted by the single MCM subunit are colored orange. Ribose/base contacts observed in most but not all subunits (see Figure S6 and S7A,E). Insert: locations of the ssDNA-binding loops in the MCM primary sequence. (D) Schematic demonstrating repeating nature of MCM-ssDNA contacts. For variations in sugar/base contacts, see Figure S7E. Bolder colors highlight ssDNA-binding motif of a single MCM subunit. Phosphates colored red. (E) MCM N-tier loops contacting DNA around the fork junction. Loops rendered as surfaces, with Mcm7 NTH separation pin also represented as cartoon. For panels (E) and (F), unpaired ssDNA is colored darker pink/orange for the lagging- and leading-strand template, respectively. (F) Detailed view of the strand separation pin displayed in cryo-EM density (mesh), inserting between the two strands of DNA at the point of unwinding. F363 makes π-π interactions with DNA. ZnF, zinc-finger; H2, helix-2; H2I, helix-2 insertion loop; PS1, presensor-1; NTH, N-terminal hairpin.

In addition to the phosphate backbone contacts, the H2I loops of several MCM subunits interact with ribose and bases primarily through residues in two conserved positions; in Mcm2, 5, 6 and 7, a basic residue contacts bases whilst an aromatic/methionine side chain contacts the sugar (Figure 3C,D and Figure S6 and S7A,E). These contacts are similar between the two conformations, except Mcm7 is disengaged in conformation 1, and the position of Mcm5 near the 5′-end and Mcm2 and 6 nearer the 3′-end in conformation 1 is reversed in conformation 2 (Figure S7E). The Mcm4 H2I basic-aromatic pair is disengaged from DNA in both conformations, while Mcm3 contains an arginine in place of the usual aromatic residue, contacting nucleotides where the path of ssDNA begins to diverge toward the fork junction in conformation 1 and disengaging in conformation 2 (Figure S7A,E). Additional basic residues at a distinct position in the H2I loops of Mcm5 (R460) and Mcm2 (K587) also project toward the fork junction in conformation 1, likely contacting bases as the ssDNA is translocated from the N-tier toward the C-tier (Figure S7A,E [top]). These contacts might be important to guide ssDNA along the correct path from the fork junction toward engagement by the MCM motor domains.

## Mechanism of strand separation by the replisome

Single-stranded DNA translocation by the C-tier motor domains must be coupled to dsDNA unwinding during replication and the high resolution of our structure at the fork junction reveals how this is accomplished. Consistent with lower resolution structures (Georgescu et al., 2017; Goswami et al., 2018), dsDNA enters the circle of zinc finger (ZnF) domains on top of the N-terminal face of the MCM ring and this is where strand separation occurs (Figure 3A). Multiple structural features contact DNA in this region, several of which require extensive remodeling from their positions in the MCM double hexamer (Figure 3E, Figure S8A,B and Movie S2). This remodeling enables the N-terminal hairpins (NTHs) of Mcm6 and Mcm4 to engage the lagging-strand template ahead of strand separation, where they help guide the incoming dsDNA onto the NTH of Mcm7 which acts as a separation pin, splitting the two strands of the incoming duplex (Figure 3F). The separation pin also undergoes extensive rearrangement from the double hexamer, where it is engaged with the Mcm5 ZnF of the second hexamer (Figure S8B); the lack of this second Mcm5 ZnF in the active helicase enables both the DNA and Mcm7 NTH to reposition, such that the Mcm7 NTH can insert between the two strands of DNA at the fork junction. Furthermore, it remodels to form a helical turn at its apex, positioning an invariant phenylalanine to base-stack with the last base pair in the DNA (Figure 3E,F and Figure S8A-C and Movie S2). Positioning hydrophobic residues to appose the last base pair is a hallmark of separation pins in diverse helicases, including the T7 replisome (Figure S8D) (Gao et al., 2019). Two separation pin lysines previously interacting with the Mcm5 ZnF in the double hexamer, one of which is invariant, contact the unwound leading-strand template at which point ssDNA is kinked almost 90° before continuing towards the C-tier (Figure 3F and Figure S8A-C and Movie S2). We propose this arrangement of intricate DNA contacts surrounding the fork junction function destabilizes the DNA duplex and blocks entry of the lagging-strand template to the MCM central channel while the leading-strand template is translocated through.

After strand separation, a single residue is clearly resolved for the unwound lagging-strand template which runs above the Mcm7 NTH approaching the ZnFs of Mcm3 and Mcm5 (Figure 3E and Movie S2). However, it is not further resolved beyond this point despite the fork substrate containing 14 additional nucleotides of ssDNA (Figure S1A), consistent with prior lower-resolution CMG structures (Georgescu et al., 2017; Goswami et al., 2018). Based on the position of the final nucleotide resolved in our structures, we consider it most likely the lagging-strand template will exit between the ZnF domains of Mcm3/5 or Mcm5/2 (Figure S8E).

## Structure of Csm3/Tof1

Before the template is unwound, dsDNA projects towards Csm3/Tof1 at the front of the replisome (Figure 1B,C). Tof1 (13-781) forms a right-handed α-solenoid with a crescent-like architecture that mimics the curvature of the MCM ring, traversing the N-tier face above Mcm2, 6, 4 and 7 (Figure 1B,C and 4A and Movie S3). The α-solenoid is composed of nine tandem 2- or 3-helix units resembling either armadillo or HEAT repeats which can be subdivided into a Head (repeats 1-5) and Body (repeats 6-9) (Figure 4A,B and Figure S9A). The first six repeats superpose well with the crystal structure of the N-terminus of human Timeless (Holzer et al., 2017), while repeats 5-9 are reminiscent of p120 catenin (Chi et al., 2002) (Figure S9B,C). Clear density for the C-terminal third of Tof1 is not observed, indicating it is flexibly tethered. Two large insertions absent from the Timeless crystal structure (Holzer et al., 2017) embellish the α-solenoid; a 35-residue Ω-loop and an ∼100 amino acid region, hereafter referred to as the *MCM-plugin,* that extends in a large loop over the surface of the MCM N-tier encircling the base of the Tof1 Body (Figure 4A,B and Figure S9D-F and S10A)

**Figure 4.**
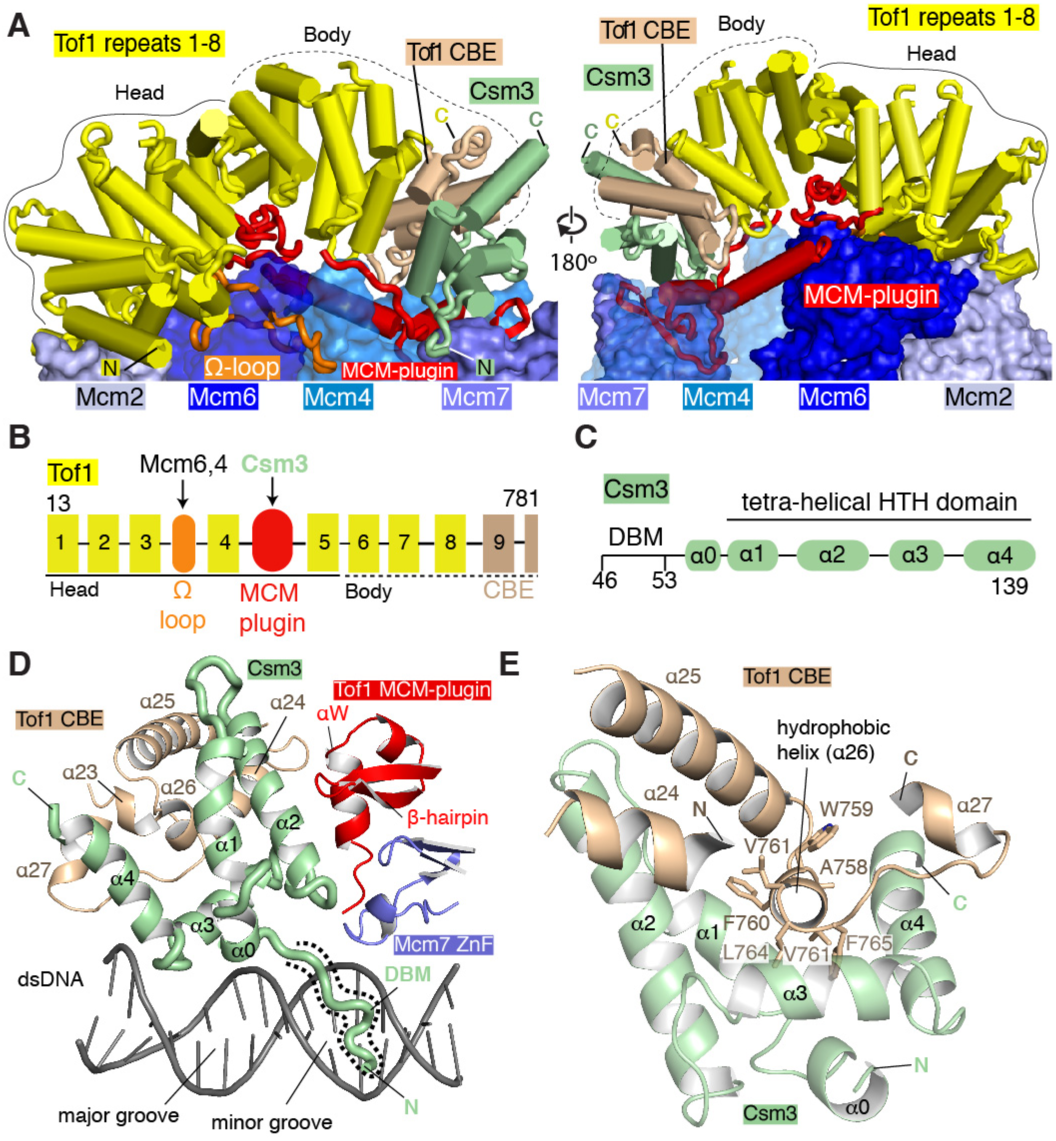
Csm3/Tof1 structure. (A) Structures of Tof1 and Csm3 shown as cylinders above the MCM N-tier (surface representation). Tof1 insertions (cartoon representation): the Ω-loop (orange) and the MCM-plugin (red) are highlighted. The Csm3-binding element (CBE) of Tof1 is colored brown. The positions of the Tof1 Head and Body are outlined with solid and dashed black lines respectively. For clarity dsDNA is not shown. (B) Schematic illustrating the positions of Tof1 helical repeats (numbered 1-9, see Figure S8A for repeat assignment), CBE, Ω-loop and MCM-plugin. The Head and Body are marked with solid and dashed black lines respectively and primary interaction sites in Tof1 insertions for MCM subunits and Csm3 are denoted by arrows. (C) Schematic illustration of Csm3 domain architecture. (D) Overview of Csm3 structure (46-139) and its interface with Tof1 and the Mcm7 ZnF. The Csm3 DBM is highlighted by a dashed outline. (E) Overview of interactions between Csm3 and the Tof1 CBE. Hydrophobic residues from Tof1 helix α26 are shown. CBE, Csm3-binding element; DBM, DNA binding motif.

We resolve 94 residues of Csm3 comprising five α-helices, which fold around a hydrophobic core to form a tetra-helical helix-turn-helix (HTH) domain (α1-4), with a short DNA binding motif (DBM) preceding α0 (Figure 4A,C-D and Figure S10B). The Csm3 HTH encases a hydrophobic helix (*α*26) at the C-terminal end of the Tof1 Body, part of a larger interface called the Csm3 binding element (CBE) (Figure 4, Figure S9F and Movie S3). This extensive interface and additional contacts with the MCM-plugin ensure Csm3 is correctly positioned in the replisome.

## Csm3/Tof1 interactions with MCM and dsDNA

In our structure Csm3/Tof1 binds to MCM exclusively via Tof1. Although the N-terminus of the α-solenoid forms an interface with the NTE of Mcm2 (Figure S11A), binding is mediated primarily by the MCM-plugin and Ω-loop (Figure 5A, Figure S9F and Movie S4). The MCM-plugin contains four structural features (Figure 5A; Bridge, Anchor, Wedge, L-loop). The Bridge comprises an α-helix spanning the helical domains of Mcm6 and Mcm4 making contacts with Mcm6 through a small conserved hydrophobic patch flanked by two glutamates (Figure 5A, Figure S10A and S11B). It is anchored on Mcm4 by a short loop (Anchor) positioned in a cleft between the Mcm4 helical domain, ZnF and OB-fold (Figure 5A, Figure S10A and S11C). The MCM-plugin then stretches upwards into a β-hairpin and short helix αW (collectively the Wedge) sandwiched between regions of Mcm4 and Mcm7 and beneath Csm3 (Figure 4A,D, 5A and Figure S11D). Finally, it returns towards the Head, contacting dsDNA via a short DNA binding motif (DBM), before looping back in an L-shape (L-loop) over the surface of Mcm6, where it binds both the helical domain and N-terminal extension (NTE) (Figure 5A and Figure S11E).

**Figure 5.**
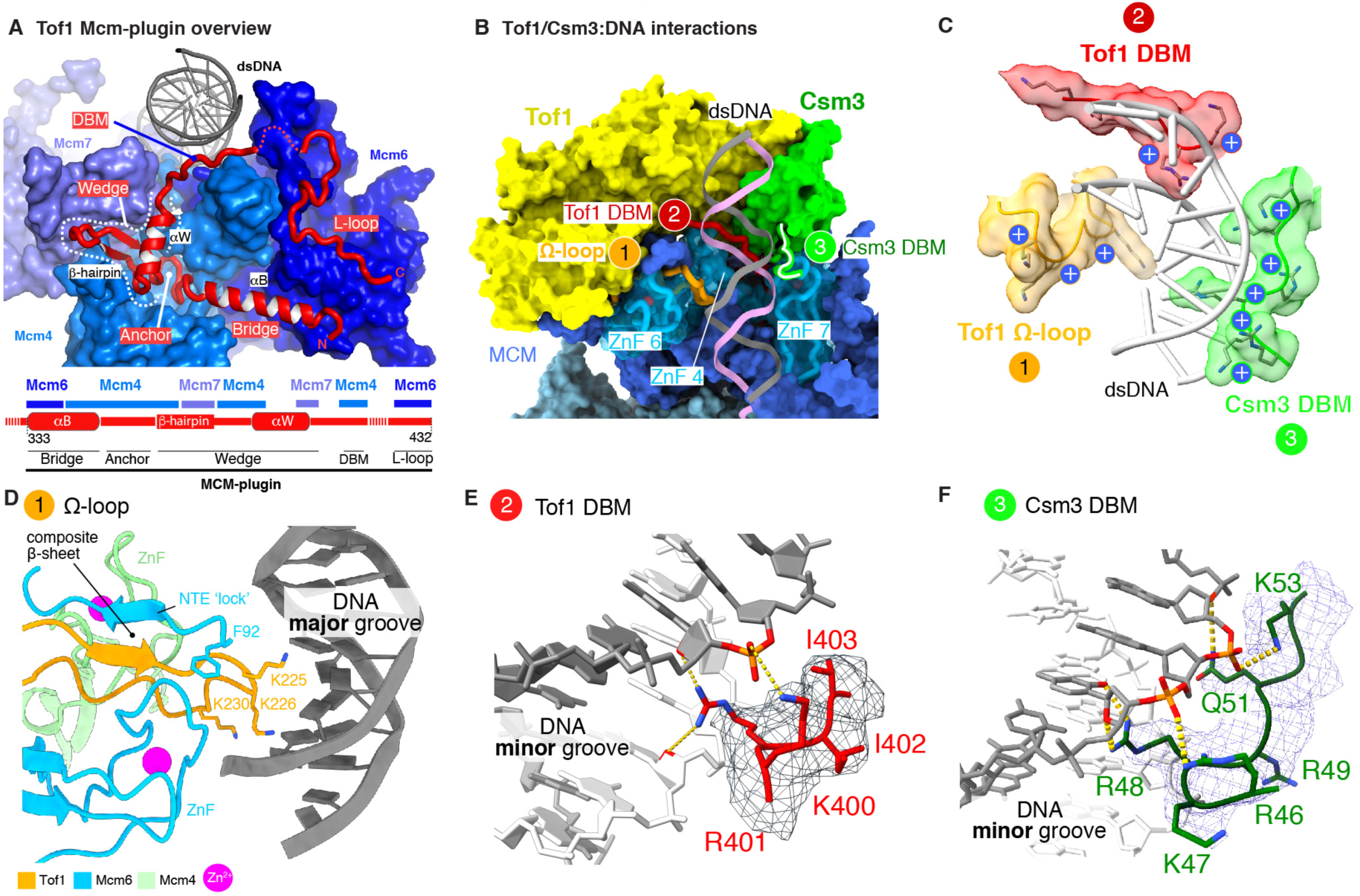
Tof1 interactions with MCM and DNA. (A) Overview of the Tof1 MCM-plugin and its position on the MCM N-tier. Top: The MCM-plugin is shown in cartoon representation above the MCM N-tier (surface representation) and structural elements involved in MCM binding are illustrated, as is the location of a DNA binding motif (DBM). Bottom: Schematic illustrating the organization of the MCM-plugin. Structural elements involved in MCM binding are illustrated, together with the specific Mcm subunits that they bind. (B) Overview of Csm3/Tof1 dsDNA contacts at the front of the replisome. The Mcm4, 6 and 7 ZnF domains important for Csm3/Tof1 binding are displayed as cartoons in transparent surfaces. (C) Closeup view of the Csm3/Tof1 dsDNA grip. For simplicity only the Ω-loop and DBMs are shown. (D) Detailed view of Ω-loop interactions with Mcm6, Mcm4 and dsDNA. (E, F) Detailed views of the Tof1 (E) and Csm3 (F) DBMs with the cryo-EM density for the DBMs shown as mesh. DBM; DNA binding motif.

Being located at the front of the replisome enables Csm3/Tof1 to ‘grip’ the parental DNA duplex via a network of interactions with the phosphate backbone and both the major and minor grooves (Figure 5B,C and Figure S9F and Movie S5). The ‘grip’ embraces three-quarters of a turn of dsDNA and comprises contacts mediated by the Ω-loop and conserved DBMs from both the MCM-plugin and Csm3. The Ω-loop protrudes from the α-solenoid towards dsDNA, slotting between the ZnF domains of Mcm6 and Mcm4 (Figure 5D and Figure S11F). The entire hydrophobic core of the loop packs against the Mcm6 ZnF positioned to one side, whereas considerably fewer contacts are made with the Mcm4 ZnF on the opposite side. The Mcm6 NTE ‘locks’ the Ω-loop in place by forming a composite β-sheet and folding back over it with the aromatic moiety of F96 contacting its own ZnF (Figure 5D). This configuration places the tip of Ω-loop facing the major groove where several lysine residues interact with the phosphate backbone (Figure 5D and Figure S9F and S10A). The equivalent loop in Timeless is considerably shorter than the Tof1 Ω-loop and therefore this mode of DNA interaction may not be universal across eukaryotes (Figure S10A).

The DBMs from the Tof1 MCM-plugin and Csm3 both bind the minor groove (Figure 5B,C and Movie S5). In the Tof1 DBM (400-404), R401 is placed into the minor groove contacting sugars approximately one turn above the fork junction and is flanked by two conserved lysine residues (K400 contacts phosphate, K404 faces the Tof1 Body) (Figure 5D,E). The Csm3 DBM (46-53) inserts R48 into the minor groove where it is positioned to contact both bases and ribose (Figure 5F). R46 and K47 are close to the backbone, perhaps to stabilize R48, while Q51 projects toward the minor groove contacting ribose and K53 coordinates the backbone phosphate at the end of the DBM. This configuration is reminiscent of minor groove binding by the N-terminal arm of homeodomain transcription factors that also precedes a HTH (Figure S12). Finally, a fourth region of Csm3/Tof1 could also bind dsDNA because several basic residues in the loop between Csm3 α3 and α4, together with the linker between Tof1 α26 and α27 are in close proximity to the backbone (Figure S12A).

## The dsDNA ‘grip’ stabilizes Csm3/Tof1 in the replisome

We showed previously Csm3/Tof1 is required for maximal replication rates in a reconstituted system (Yeeles et al., 2017) and considered the DBMs might be important for this functionality. Therefore we substituted DBM residues for alanine and purified stable heterodimers of Csm3^R49A/K53A^/Tof1 (Csm3-2A/Tof1), Csm3^K47A, R48A, R49A, Q51A, K53A^/Tof1 (Csm3-5A/Tof1), Csm3/Tof1^K400A, R401A, K404A^ (Csm3/Tof1-3A) and double mutants where both subunits were mutated (Csm3-2A/Tof1-3A and Csm3-5A/Tof1-3A) (Figure S13A). Surprisingly, Figure S13B shows that all DBM mutants bound to fork DNA with comparable affinity to wild type, indicating the complex possesses additional DNA binding regions, perhaps in the ∼400 amino acid C-terminal region of Tof1 and/or the C-terminus of Csm3 not resolved in our structures. We then performed origin-dependent replication reactions (Aria and Yeeles, 2019; Taylor and Yeeles, 2018) on a linear template with the origin located to generate leading strands of 1.9 kb (Right) and 8.2 kb (Left) using conditions that require Csm3/Tof1 for rapid and efficient DNA replication (Yeeles et al., 2017) (Figure 6A,B). Figure 6C shows that mutation of either the Csm3 (2A and 5A) or Tof1 DBMs had little effect on left leading-strand products (compare lanes 3, 4 and 5 with 2). In contrast, when both subunits were mutated we observed a minor replication defect with Csm3-2A/Tof1-3A (compare lanes 2 and 6) and a marked replication defect with Csm3-5A/Tof1-3A (compare lanes 2 and 7), although not as severe as Csm3/Tof1 omission and some 8.2 kb Left lead products were still synthesized. At longer time points both double mutants supported the synthesis of fully replicated left leading strands to levels above those generated in the absence of Csm3/Tof1 (Figure S13C,D). Whereas 8.2 kb left lead products were only slightly reduced with Csm3-2A/Tof1-3A (Figure S13C, compare lanes 2 and 5), they were notably less intense for Csm3-5A/Tof1-3A (Figure S13D, lanes 2 and 5), despite comparable levels of the shorter right leading strand being synthesized. Collectively, these data indicate that individual DBMs are dispensable for rapid and efficient DNA replication, but that disrupting both motifs compromises the net rate of leading-strand synthesis.

**Figure 6.**
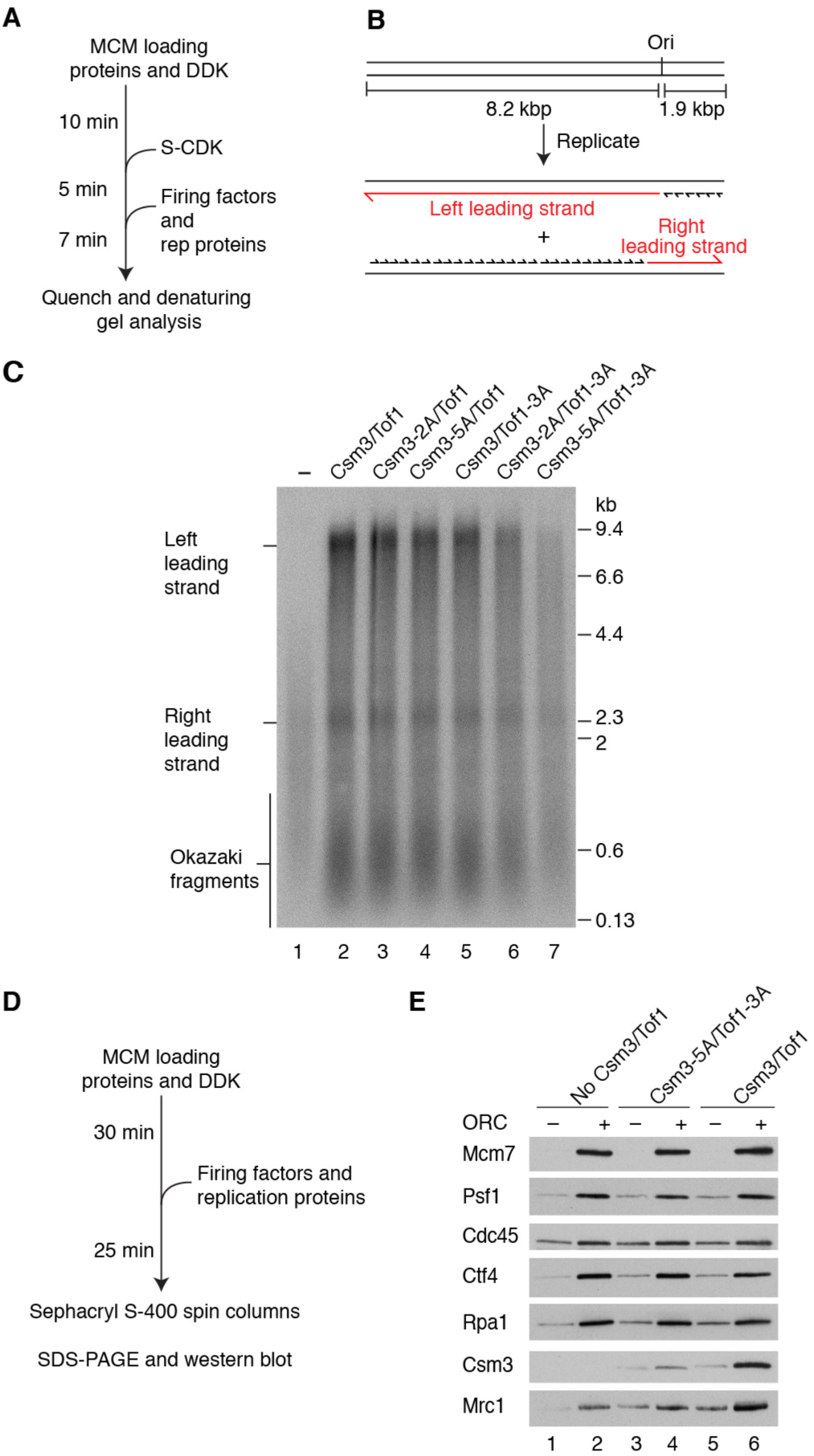
Csm3/Tof1 DNA binding is important for replisome stability. (A) Reaction scheme for origin-dependent replication assay. (B) Schematic of DNA template and anticipated replication products. (C) Origin-dependent replication reaction (7 min) with the indicated Csm3/Tof1 proteins performed as illustrated in (A). Products were separated through a 0.6% alkaline agarose gel. (D) Reaction scheme for protein association experiments. (E) Western-blot analysis of a reaction performed as in (D) with the indicated Csm3/Tof1 proteins.

Rate enhancement by Csm3/Tof1 requires Mrc1 (Yeeles et al., 2017) and Csm3/Tof1 stabilize Mrc1 in the replisome (Bando et al., 2009). We therefore considered the defects observed with the DBM double mutants might be due to compromised Mrc1 function, for example if they failed to correctly stabilize or position Mrc1 at the replication fork. To test this, we developed an assay to monitor replisome stability (Figure 6D). Elution of Mcm7, Cdc45, Psf1 (GINS), Ctf4, RPA, Csm3 and Mrc1 was dependent on DNA, ORC (required for MCM loading) and Dpb11 (required for CMG assembly) confirming the detection of replisome associated proteins (Figure S13E). In the absence of Csm3/Tof1, Mrc1 replisome association was reduced but not eliminated (Figure 6E), consistent with its ability to modestly stimulate fork rate in reactions lacking Csm3/Tof1 (Yeeles et al., 2017). Moreover this indicates that Mrc1 is stabilized in the replisome by Csm3/Tof1 as is observed *in vivo* (Bando et al., 2009). In contrast, in the reaction containing Csm3-5A/Tof1-3A, Csm3 association was appreciably weaker and levels of Mrc1 were comparable to the experiment lacking Csm3/Tof1 (Figure 6E, compare lanes 4 and 6). Figure S13F shows the Csm3/Tof1 single DBM mutants associated with the replisome like wild type, as did Mrc1, whereas there was a minor reduction in both proteins with Csm3-2A/Tof1-3A. We also observed a defect in the binding of Csm3, Tof1 and Mrc1 to CMG in glycerol gradients with Csm3-5A/Tof1-3A (Figure S13G). The weaker replisome association displayed by Csm3-5A/Tof1-3A indicated the complex may function more distributively during replication and Figure S13H supported this idea: whereas 10 nM Csm3/Tof1 was saturating for replication, DNA synthesis was enhanced across the entire titration range (2.5-80 mM) for Csm3-5A/Tof1-3A. Taken together, these results reveal the Csm3/Tof1 dsDNA ‘grip’ stabilizes the entire fork protection complex in the replisome, strongly suggesting the replication defect observed with Csm3-5A/Tof1-3A (Figure 6C) stems from its inability to stabilize Mrc1 (Figure 6E).

## The Csm3/Tof1 dsDNA ‘grip’ is important for replication fork pausing

Csm3/Tof1 is essential for directional fork pausing at replication fork barriers (RFBs) to limit head-on collisions between the replisome and RNA polymerase (Hizume et al., 2018; Mohanty et al., 2006; Takeuchi et al., 2003). The RFB contains multiple binding sites for Fob1 (Kobayashi, 2003), which is required for barrier activity (Kobayashi and Horiuchi, 1996). To assess if DNA binding by Csm3/Tof1 facilitates fork pausing at the RFB, we performed replication reactions in the presence of Fob1 on a linear template containing the RFB in the non-permissive orientation ∼3 kb left of the origin (Figure 7A). Fork pausing will generate a slowly migrating stalled fork and an ∼3 kb stalled left leading strand that can be visualized in native and denaturing gels respectively. Figure 7B shows that Csm3/Tof1-dependent fork stalling was recapitulated in this system (compare lanes 1 and 2). Similar results were observed in a lower salt buffer (Yeeles et al., 2017) that supports fully replicated left lead products in the absence of Csm3/Tof1 (Figure S14A, compare lanes 1 and 2). Whereas the responses of Csm3-2A/Tof1 and Csm3/Tof1-3A to the RFB were comparable to wild type, we observed a 30-40% reduction in pausing for Csm3-5A/Tof1 and an ∼60% reduction with Csm3-2A/Tof1-3A (Figure 7B,C and Figure S14A,B). RFB activity with Csm3-5A/Tof1-3A was reduced almost to background levels. Importantly, although the replisome association of this mutant was compromised (Figure 6E and Figure S13G), it retained the ability to significantly stimulate replication such that the 8.2 kb left lead products in Figure 7B were highly dependent on the mutant protein (compare lanes 1 and 7). The fork pausing defect displayed by Csm3-5A/Tof1-3A therefore likely represents a specific defect in responding to the RFB and is not simply a consequence of the protein being absent from the replisome. These findings demonstrate the Csm3/Tof1 dsDNA ‘grip’ is crucial for efficient replication fork pausing at the RFB.

**Figure 7.**
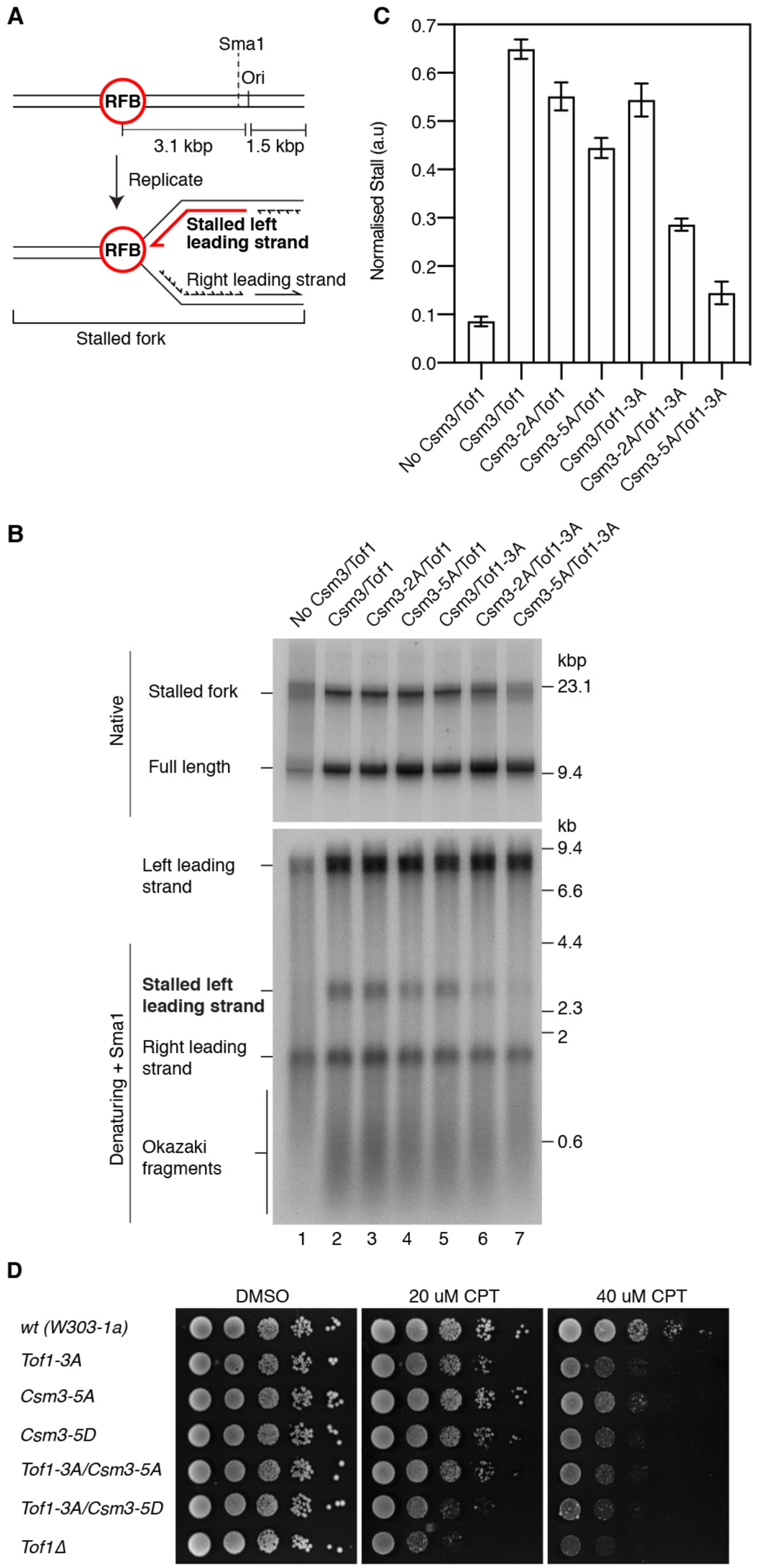
The Csm3/Tof1 dsDNA ‘grip’ is required for efficient fork pausing. (A) Schematic of the template used for RFB experiments and the anticipated products of fork stalling at the RFB. (B) Origin-dependent replication reaction performed for 20 min in the presence of Fob1. The Csm3/Tof1 concentration was increased to 80 nM to increase replication efficiency (Figure S10H). Reaction products were treated with Sma1 prior to denaturing gel electrophoresis to remove heterogeneity in the position of leading-strand initiation (Taylor and Yeeles, 2018). (C) Quantitation of experiments performed as in (B). Error bars represent the standard error of the mean (SEM) from 3 experiments. (D) Spot dilution assay with Tof1 and Csm3 DBM mutants. 10-fold serial dilutions were plated and grown at 25°C for 3 days. RFB, replication fork barrier.

Fork pausing at proteinaceous barriers can also have deleterious consequences, for example when forks stall at covalently trapped topoisomerase I (Topo I) complexes following treatment of cells with camptothecin (CPT). Csm3 and Tof1 are important for cellular tolerance of CPT (Redon et al., 2006), although their mechanism of action is unclear. Given the DBMs are important for pausing at the RFB, we considered they may also be involved in the response to fork stalling during CPT treatment. To test this hypothesis, we generated haploid yeast expressing either Csm3-5A or Tof1-3A from the endogenous promoter and a Csm3 charge-reversal mutant (*Csm3-5D*). *tof1-3A* and *csm3-5D* exhibited mild growth defects at 20 µM CPT, whereas *csm3-5A* grew almost as wild type (Figure 7D). At 40 μM CPT both *tof1-3A* and *csm3-5D* displayed significant growth defects and *csm3-5A* showed mild sensitivity. Combining *csm3-5D* and *tof1-3A* had an additive effect on CPT sensitivity such that cell growth was only marginally better than *tof1Δ*. Importantly, none of the mutants displayed sensitivity to hydroxyurea (HU) and only *tof1-3A/csm3-5D* showed a very mild sensitivity to methyl methanesulphonate (MMS) (Figure S14C), indicating the observed defects with CPT were not a general response to replication stress. Collectively, these data reveal the Csm3/Tof1 dsDNA ‘grip’ participates in the maintenance of genome stability following Topo I inhibition by CPT, likely by stabilizing stalled replication forks.

## Discussion

Our structures provide the first near-atomic resolution views of the CMG helicase bound to accessory proteins at a replication fork, providing the most complete picture of the eukaryotic replisome to date (Figure S15). They reveal the structure of Csm3/Tof1 and its extensive network of interactions with MCM, which redefines the architecture of the front of the replisome by placing the complex ahead of CMG (Figure 1). This advanced positioning of Csm3/Tof1 should facilitate Tof1 recruitment of Topo I ahead of the fork (Park and Sternglanz, 1999) - potentially to limit excessive fork rotation (Schalbetter et al., 2015). The conformation of Csm3/Tof1 and the dsDNA ‘grip’ we observe likely reflect an active mode of replisome progression because mutation of both DBMs compromises replication in the absence of exogenous DNA damage (Figure 6C). Moreover, our work demonstrates the ‘grip’ plays a key role in stabilizing the entire fork protection complex in the replisome and is especially important for efficient fork pausing at the RFB (Figure 7B). Although the mechanism of RFB recognition is currently unknown, we speculate the dsDNA ‘grip’ might function to stabilize the replisome on the template once Fob1 is encountered by the fork. Alternatively, it might be required to correctly position Csm3/Tof1 at the front of the replisome to recognize Fob1, perhaps via a direct protein:protein interaction. Gripping dsDNA could also enable Csm3/Tof1 to detect structural perturbations in the DNA template or protein roadblocks in advance of CMG, which is likely to be important for its fork stabilization functions. This hypothesis is consistent with the involvement of the dsDNA ‘grip’ in the cellular tolerance of CPT (Figure 7D).

The normal replication rates displayed by the single DBM mutants (Figure 6C), the dependence on Mrc1 for Csm3/Tof1-dependent rate enhancement (Yeeles et al., 2017), and the reduced association of Mrc1 in experiments using Csm3-5A/Tof1-3A (Figure 6E) indicate that Csm3/Tof1 promotes rate enhancement indirectly likely by stabilizing and/or positioning Mrc1 in the replisome. It was recently reported that the human Tof1 ortholog Timeless is displaced from replisomes to slow replication in response to redox changes (Somyajit et al., 2017) and our findings now indicate this displacement could be mediated by disrupting the dsDNA ‘grip’.

Our discovery that Mrc1 stretches from the front of the replisome to the rear (Figure 2) affords a mechanism for coordination of events ahead of the fork - perhaps monitored by Csm3/Tof1 - with leading-strand polymerization. Moreover, the positioning of Mrc1 across one side of CMG could enable it to directly modulate helicase activity to control fork rate and maintain the coupling of DNA synthesis to CMG template unwinding when leading-strand polymerization is compromised (Katou et al., 2003). The N-terminal half of Mrc1 interacts with the flexibly linked catalytic domain of Pol ε (Lou et al., 2008) and several amino acids from this region crosslinked to Mcm6 and Mcm2 in close proximity to where the unwound leading-strand template will emerge from CMG (Figure 2A). We hypothesize Mrc1 might stimulate fork rate by tethering the catalytic domain of Pol ε to this region of CMG to facilitate optimal helicase-polymerase coupling.

Our structures also offer the highest resolution views of eukaryotic CMG:DNA interactions to date (Figure 3). This has revealed a set of phosphate contacts displaying repetition across all six MCM subunits in the heterohexamer, presumably to ensure ssDNA can be engaged by all subunits at different points during a rotary translocation cycle (Eickhoff et al., 2019). In contrast, the sugar/base contacts formed by the H2I loops display greater diversity between MCM subunits; the contribution of these contacts to translocation in eukaryotic CMG remains unknown and roles in DNA melting during helicase activation have been hypothesized (Meagher et al., 2019). The identification of a strand separation pin defines the point of template unwinding in the replisome and reveals the mechanism by which it occurs. The stable positioning of dsDNA ahead of CMG (Eickhoff et al., 2019) and the utilization of a defined separation pin, will ensure the correct positioning of unwound template strands within the replisome for downstream processes. Moreover, if the C-tier motor domains were to disengage from ssDNA, the extensive contacts around the fork junction, together with the Csm3/Tof1 dsDNA ‘grip’, might function to prevent helicase backtracking.

The cryo-EM structure of the fork protection complex bound to CMG at a replication fork represents a significant advance in our understanding of eukaryotic replisome structure and mechanism. Yet, given its complexity, subunit dynamics and sophisticated regulation, much work remains to be done. The work presented here provides an ideal platform to build ever-more complex replisome assemblies for analysis by cryo-EM. This will be crucial to facilitate a complete mechanistic description of this remarkable molecular machine.

## Acknowledgements

We are grateful to S. Villa-Hernandez for preliminary experiments with the RFB and Tom Deegan for advice on replisome stability assays. We thank J. Diffley and K. Labib for yeast strains and antibodies and L. Passmore, D. Barford, J. Sale, J. Baxter and H. Williams for comments on the manuscript. We are grateful to A. Casañal, P. Emsley, G. Murshudov, R. Nicholls and O. Kovalevskiy for advice on model building and refinement, M. Anandapadamanaban for assistance with GraFix and S. Villa-Hernández for assistance with yeast strain construction. We thank S. Chen, C. Sava, J. Brown, J. Grimmett and T. Darling for smooth running of the EM and computing facilities. We acknowledge Diamond for access and support of the Cryo-EM facilities at the UK national electron bio-imaging centre (eBIC), proposals EM20976-1 & EM17434-38, funded by the Wellcome Trust, MRC and BBSRC. We are grateful to A. Klyszejko, D. Clare, T. Hoffman and C. Wigge for providing support in the use of the facilities at the eBIC. Cryo-EM data collection (experiment session MX/2091) was performed on cryo-electron microscope CM01 at the European Synchrotron Radiation Facility (ESRF), Grenoble, France. We are grateful to G. Effantin at the ESRF for assistance in using CM01. This work was supported by the Medical Research Council, as part of United Kingdom Research and Innovation (MRC grant No. MC_UP_1201/12 to J.T.P.Y).

## Author contributions

D.B. and M.J-B. purified proteins and performed all cryo-EM sample preparation, data collection, analysis and model building. V.A. generated yeast strains and performed spot assays. G.C. performed preliminary cryo-EM sample preparation, data collection and analysis. M.S. performed and analysed XL-MS. J.T.P.Y. conceived the study, generated protein expression strains, purified proteins and performed DNA replication and spot assays. J.T.P.Y., M.J-B. and D.B. wrote the manuscript.

## Materials and Methods

### Yeast strains

Vectors and strains were constructed using standard molecular biology techniques (see Table S4 for details). All genes for protein expression were codon optimized as described (Yeeles et al., 2015). All mutant haploid yeast strains were isolated by tetrad dissection of heterozygous diploid strains. Coding sequences for all genes were verified by sequencing, as were the coding regions of mutant alleles of Csm3 and Tof1 following PCR amplification from genomic DNA.

### Protein purification

Cdt1•Mcm2-7, ORC, Cdc6, DDK, Sld3/7, Sld2, Cdc45, S-CDK, Dpb11, GINS, Pol ε, Mcm10, RPA, RFC, PCNA, Pol α, Pol δ, Csm3/Tof1 and Ctf4 were purified as previously described (Taylor and Yeeles, 2018; Yeeles et al., 2015; Yeeles et al., 2017).

#### Mrc1 purification

Mrc1 was purified as previously described (Yeeles et al., 2017) but with the following modifications. The growth temperature during protein expression was reduced from 30 °C to 20 °C. All subsequent steps were performed at 4 °C. Lysed cell powder from a 10-15 L culture was resuspended in 50 mM Tris-HCl pH 8, 10% glycerol, 0.005% TWEEN 20, 0.5 mM Tris(2-carboxyethyl)phosphine hydrochloride (TCEP) and 400 mM NaCl (buffer M + 400 mM NaCl) + protease inhibitors (1 Roche complete tablet per 50 ml buffer). Insoluble material was cleared by centrifugation (235,000g, 4 °C, 1 hour) and 2-4 ml FLAG M2 Affinity gel (Sigma) was added to the supernatant. The sample was incubated for 100 min before the resin was collected in 20 ml columns (<2 ml bed volume per column) and was washed with 75 ml buffer M + 400 mM NaCl. Columns were washed with 12.5 ml buffer M + 400 mM NaCl + 5 mM Mg(OAc)_2_ + 0.5 mM ATP, followed by 25 ml buffer M + 400 mM NaCl. Mrc1 was eluted in 1 column volume (CV) buffer M + 400 mM NaCl + 0.2 mg/ml 3x FLAG peptide and 2 CV buffer M + 400 mM NaCl + 0.1 mg/ml 3x FLAG peptide. The eluate was concentrated to ∼800 μl in an Amicon Ultra-15 30,000 NMWL concentrator and applied to a Superose 6 10/300 column (GE healthcare) equilibrated in 25 mM Tris-HCl pH 7.2, 10% glycerol, 0.005% TWEEN 20, 1 mM EDTA, 0.5 mM TCEP, 150 mM NaCl. Peak fractions were pooled, frozen in liquid nitrogen and stored at −80 °C.

#### CMG purification

Diploid yeast (yJY37) (15-30 L) were grown at 30 °C to 5 x10^7^ cells per ml in YEP (1.1% yeast extract, 2.2% bactopeptone, 55 mg/L adenine hemisulphate) + 2% w/v raffinose before induction by addition of galactose to 2% w/v final concentration from a 20% w/v stock. Cell growth was continued for 3 hours at 30 °C before cells were harvested by centrifugation, washed in 150 ml 40 mM HEPES-NaOH pH 7.5, 10% glycerol, 0.005% TWEEN 20, 0.5 mM TCEP, 150 mM NaOAc (buffer C + 150 mM NaOAc) and resuspended in a minimal volume of buffer C + 150 mM NaOAc + protease inhibitors (1 Roche complete tablet per 50 ml buffer). Cell paste was frozen in liquid nitrogen and cells were lysed using a pestle and mortar filled with liquid nitrogen. Lysed cell powder (typically from a 15 L culture) was resuspended in buffer C + 150 mM NaOAc + protease inhibitors and insoluble material removed by centrifugation (235,000g, 4 °C, 1 hour). FLAG M2 Affinity gel (8 ml) was added to the lysate and incubated for 90 min at 4 °C. Resin was collected in 20 ml columns (<2 ml bed volume per column) and washed with 80 ml per column buffer C + 150 mM NaOAc. Columns were then washed with 10 ml buffer C + 150 mM NaOAc + 5 mM Mg(OAc)_2_ + 0.5 mM ATP followed by 25 ml buffer C + 150 mM NaOAc. Proteins were eluted with 1 CV buffer C + 150 mM NaOAc + 2mM CaCl_2_ + 0.2 mg/ml 3x FLAG peptide and 2 CV buffer C + 2mM CaCl_2_ + 150 mM NaOAc + 0.1 mg/ml 3x FLAG peptide. Calmodulin sepharose 4B (GE healthcare) (1 ml) was immediately added to the eluate, which was incubated for 30 min before the resin was collected in a 20 ml column. The flow-through was reapplied to the column twice before washing the column with 25 CV buffer C + 150 mM NaOAc + 2mM CaCl_2_. CMG was eluted in 8 CV buffer C + 150 mM NaOAc + 2 mM EDTA + 2 mM EGTA. Eluate was applied to a MonoQ PC 1.6/5 (GE healthcare) equilibrated in 25 mM Tris-HCl pH 7.2, 10% glycerol, 0.005% TWEEN 20, 0.5 mM TCEP, 150 mM KCl. CMG was eluted with a 20 CV gradient from 150-1000 mM KCl and peak fractions were dialysed overnight against 500 ml 25 mM HEPES-KOH pH 7.6, 40 mM KOAc, 40 mM K-glutamate, 2 mM Mg(OAc)_2_, 0.25 mM EDTA, 0.5 mM TCEP, 20% glycerol. Protein was frozen in liquid nitrogen and stored at −80 °C.

#### Fob1 purification

10 L yJY39 were grown at 30 °C to 4.5 x10^7^ cells per ml in YEP + 2% w/v raffinose before induction by addition of galactose to 2% w/v final concentration from a 20% w/v stock. Cell growth was continued for 3 hours at 30 °C before cells were harvested by centrifugation, washed in 150 ml 25 mM Tris-HCl pH 7.2, 1 mM EDTA, 10% glycerol, 0.02% NP-40-S, 0.5 mM DTT, 400 mM NaCl (buffer F + 400 mM NaCl) and resuspended in a minimal volume of buffer F + 400 mM NaCl + protease inhibitors (1 Roche complete tablet per 25 ml buffer). Cell paste was frozen in liquid nitrogen and cells were lysed using a pestle and mortar filled with liquid nitrogen. Lysed cell powder was resuspended in buffer F + 400 mM NaCl + protease inhibitors and insoluble material removed by centrifugation (235,000g, 4 °C, 1 hour). FLAG M2 Affinity gel (2.5 ml) was added to the lysate and incubated for 3 hours at 4 °C. Resin was collected in a 20 ml column and washed with 80 ml buffer F + 400 mM NaCl followed by 20 ml buffer F + 200 mM NaCl. Fob1 was eluted in 8 ml buffer F + 200 mM NaCl + 0. 2 mg/ml 3x FLAG peptide. The eluate was diluted in buffer F to the equivalent of 150 mM NaCl and was applied to a 1 ml MonoQ column equilibrated in buffer F + 150 mM NaCl. Protein was eluted with a 25 CV gradient from 150-1000 mM NaCl in buffer F. Peak fractions were pooled and dialysed against buffer F + 150 mM NaCl for 3 hours prior to freezing in liquid nitrogen and storage at −80 °C.

### Preparation of fork DNA for cryo-EM sample preparation

Fork DNA was annealed by mixing equal volumes of Fork-Lead and Fork-Lag oligos (Integrated DNA Technologies) allowing to cool gradually from 75 °C to room temperature. The Fork-Lead and Fork-Lag stock solutions were made at 53 μM each in 25 mM HEPES-NaOH, pH 7.5, 150 mM NaOAc, 0.5 mM TCEP, 2 mM Mg(OAc)_2_. The sequence of each oligo was a modified version of the fork used in Georgescu *et al* (Georgescu et al., 2017); Fork-Lead was 5’-(Cy3)TAGAGTAGGAAGTGA(Biotinylated-dT)GGTAAGTGATTAGAGAATTGGAGAGTGTG(T)_34_T*T*T*T*T*T, where * denotes phosphorothioate backbone linkages. Fork-Lag was 5’-GGCAGGCAGGCAGGCACACACTCTCCAATTCTCTAATCACTTACCA(Biotinylated-dT)CACTTCC TACTCTA.

### Glycerol 10-30% gradient preparation

For co-expression experiments, Buffer A (40 mM HEPES-NaOH, pH 7.5, 150 mM NaOAc, 0.5 mM TCEP, 10% v/v glycerol) was layered on top of an equal volume of freshly prepared Buffer B (Buffer A, except 30% v/v glycerol and 0.16% glutaraldehyde [Sigma]) in a SW40 Ti rotor 14 mL tube (Beckman) and gradients made using a gradient-making station (Biocomp Instruments, Ltd.) before cooling on ice.

For *in vitro* reconstitution experiments, 500 μM AMP-PNP and 3 mM Mg(OAc)_2_ were added to Buffers A and B. Buffer B was supplemented with a second cross-linking agent bis[sulfosuccinimidyl]suberate (BS^3^, Thermo Fisher Scientific) at 2 mM. These were layered in equal volumes in a TLS-55 rotor 2.2 mL tube (Beranek Laborgerate) and gradients prepared using a gradient-making station (Biocomp Instruments, Ltd.) before cooling on ice.

### Reconstitution of CMG-Csm3/Tof1-Mrc1-Ctf4-DNA complexes for cryo-EM

The following components were sequentially mixed with CMG stock solution while on ice. The final reaction volume of 65 μL contained 0.5 μM CMG with a 1.5 molar excess of all other components. 500 μM AMP-PNP / 3 mM Mg(OAc)_2_ was maintained throughout. First, the fork DNA stock solution was added to CMG and incubated for 1 h. Subsequently, Csm3/Tof1 and Ctf4 were pre-mixed and added to the CMG:DNA reaction mixture. After 10 min incubation, Mrc1 was added to mixture for a further 45 min.

Before loading onto the glycerol gradient, the reaction volume was diluted 2.5-fold using buffer D (25 mM HEPES-NaOH, pH 7.5, 150 mM NaOAc, 0.5 mM TCEP, 500 μM AMP-PNP, 3 mM Mg(OAc)_2_). The complex was separated by centrifugation (Beckman TLS-55 rotor at 200,000*g* and 4 °C for 2 h) and 100 μL fractions were manually collected. The fraction containing the complex was identified by SDS-PAGE. Relevant fractions were pooled (total 200 μL) and buffer exchanged with cryo-EM buffer (buffer D except 100 μM AMP-PNP + 0.005% v/v TWEEN 20 [Protein Grade, Sigma]) during six rounds of ultrafiltration in 0.5 mL 30K MWCO centrifugal filters (Amicon) using a tabletop centrifuge (21,000*g,* 4 °C, 1 min/round,). Sample was finally concentrated down to ∼25 μL and immediately used for cryo-EM grid preparation.

### Co-expression and purification of CMG-Csm3/Tof1-Mrc1-Ctf4 complexes for cryo-EM

Cultures of yJY74 (see Table S4 for details) were grown in YEP with 2% w/v raffinose (15 L) at 30 °C, to a density of ∼6 x 10^7^ cells/mL before inducing overexpression by addition of 2% w/v galactose for 3 h under the same conditions. Cells were harvested by centrifugation (3,000*g*, 8 min, 4 °C), washed and resuspended with Lysis buffer (40 mM HEPES-NaOH, pH 7.5, 150 mM NaOAc, 10 % glycerol, 0.005% v/v TWEEN 20, 0.5 mM TCEP, 1 protease-inhibitor tablet [cOmplete EDTA-free, Roche] per 25 mL buffer), and then flash-frozen as pellets in liquid nitrogen.

Cells were lysed using a Freezer/Mill 6870D SPEX Sample Prep (2 cycles, 1 min pre-cool, 2 min run-time, 1 min cool-time, rate 10 cps) before thawing in Lysis buffer. All subsequent steps were performed at 4 °C unless specified otherwise. The lysate was clarified by centrifugation (160,000*g*, 45 min) and the supernatant filtered (0.45 μm PVDF syringe filters, Elkay Laboratory Products UK). The supernatant was then incubated with 8-10 mL anti-FLAG M2 affinity agarose gel (Sigma), rotating at 7 rpm for 60-90 min. The next affinity chromatography steps were done at room temperature using ice-cold buffers, unless stated differently. The supernatant was split between gravity flow columns (14 cm Econo-Pac, BioRad) and the flow-through re-applied once and each column washed twice with 30 mL Wash buffer A (40 mM HEPES-NaOH, pH 7.5, 150 mM NaOAc, 10 % v/v glycerol, 0.005% v/v TWEEN 20, 0.5 mM TCEP). Each column was washed once with 12.5 mL ATP buffer (40 mM HEPES-NaOH, pH 7.5, 150 mM NaOAc, 10 % v/v glycerol, 0.005% v/v TWEEN 20, 0.5 mM TCEP, 500 μM ATP and 5 mM Mg(OAc)_2_), incubating for 5 min before a final wash with 20 mL of Wash buffer B (40 mM HEPES-NaOH, pH 7.5, 150 mM NaOAc, 10 % glycerol, 0.005% v/v TWEEN 20, 0.5 mM TCEP, 2 mM CaCl_2_). Protein was eluted by successive addition of FLAG-elution buffer (40 mM HEPES-NaOH, pH 7.5, 150 mM NaOAc, 10 % glycerol, 0.005% v/v TWEEN 20, 0.5 mM TCEP, 2 mM CaCl_2_, 0.1-0.2 mg/mL 3xFLAG peptide [Sigma]). The first elution was done with 1 bed volume (BV) of Elution buffer with 0.2 mg/mL 3xFLAG peptide followed by 2 BV of the same buffer containing 0.1 mg/mL 3xFLAG peptide. The resin was then washed with 1 BV of Wash buffer B.

The FLAG-eluate was pooled and incubated with up to 1.2 mL Calmodulin Sepharose 4B affinity resin (GE Healthcare), rotating at 7 rpm for 1 h at 4 °C. The sample was applied to a gravity flow column (9 cm Poly-Prep Chromatography Columns, Bio-Rad) and the flow-through reapplied twice. The column was washed twice with 20 mL Wash buffer C (25 mM HEPES-NaOH, pH 7.5, 150 mM NaOAc, 0.5 mM TCEP, 2 mM CaCl_2_) before elution using 3-5 mL Calmodulin-elution buffer (25 mM HEPES-NaOH, pH 7.5, 150 mM NaOAc, 0.5 mM TCEP, 2 mM EDTA and 2 mM EGTA). The sample was concentrated to 300 μL using 0.5 mL 30K MWCO centrifugal filters (Amicon) in a tabletop centrifuge (21,000*g*, 4 °C) and half was applied to a 10-30% glycerol gradient. The complex was separated by centrifugation in SW40 Ti rotor (Beckman) at 140,000*g* for 15 h at 4 °C. Samples were manually fractionated in 400 μL aliquots and analysed by SDS-PAGE. The relevant fraction was buffer exchanged in cryo-EM buffer without added nucleotide (25 mM HEPES-NaOH, pH 7.5, 150 mM NaOAc, 0.5 mM TCEP, 0.005% v/v TWEEN 20) over six rounds of centrifugation (21,300*g*, 1 min/round, 4 °C) in 0.5 mL 30K MWCO centrifugal filters [Amicon]. The sample was concentrated to a final volume of ∼35 μL.

In early cryo-EM datasets we observed higher compositional heterogeneity of the complex likely arising from endogenous DNA co-purifying with our sample. In an attempt to overcome this, later sample preparations contained 10 μL streptavidin-blocked fork DNA added to the relevant fraction and incubated on ice for 15 min after gradient fixation and before buffer exchange. To prepare streptavidin-blocked fork DNA, 10 μL fork DNA (26.5 μM) was incubated with 12.5 μL of readily available tetravalent streptavidin (21 μM, Pierce) at room temperature for 40 min prior to addition of DNA to a purified replisome. The addition of DNA after the final centrifugation step did not improve DNA homogeneity in our cryo-EM reconstructions, and therefore data obtained from samples prepared with and without added streptavidin-blocked fork DNA were combined during processing.

### Cryo-EM grid preparation

#### Reconstituted and co-expressed complex

Quantifoil R2/2, Cu-400 mesh cryo-EM grids pre-coated with an ultra-thin (3-5 nm) amorphous carbon (produced at the LMB) were glow discharged for 5 s at plasma current of 15 mA (PELCO easiGlow). A 3 μL sample was applied on a grid and incubated for 15-30 s at 4 °C before manually blotting with filter paper for 10 s and plunge-freezing in liquid ethane.

### Data collection

#### Reconstituted sample

A total of 6,878 raw micrographs were acquired within two data sets on the same 300 keV FEI Titan Krios microscope (LMB Krios3) at a calibrated pixel size of 1.049 Å/pixel (nominal magnification of 130,000 X). The K2 Summit direct electron detector (Gatan) was used in electron counting mode with the slit width of the GIF Quantum energy filter set to 20 eV. The EPU (Thermo Fisher) software was used for automated data collection, with defocus range set at −1.4 to −2.6 μm and dose-fractionation into 20 frames with a total exposure time of 7 or 8 s to achieve dose of 37-38 e^-^/Å^2^ per micrograph.

#### Co-expressed sample

Six datasets were collected totalling 11,637 raw micrographs. The 300 keV FEI Titan Krios microscopes (LMB Krios1 and Krios2, eBIC Krios M03 and ESRF Krios1) were used with either Falcon III direct electron detector (FEI) or a K2 Summit direct electron detector (Gatan) in electron counting mode. The data were acquired at several magnifications ranging from 1.05-1.07 Å/pixel. The EPU (Thermo Fisher) was used for automated data collection with a set defocus range −1.5 to −3 μm. For data acquired with Falcon III camera the acquisition was dose-fractionated into 180 frames with an exposure time of 44 s per micrograph and a dose of 0.82 - 0.84 e^-^/pix/s. The micrographs collected with K2 camera were dose-fractionated into 20 frames with a total exposure time of 6 - 8 s to achieve a dose per micrograph of 37 - 43 e^-^/Å^2^, and the slit width of the GIF Quantum energy filter was set to 20 eV.

### Data processing and 3D-reconstruction for the reconstituted sample

The gain-corrected 20-frame movies were aligned and dose-weighted (0.25-0.27 e^-^/Å^2^/frame) by MotionCor2 (Zheng et al., 2017). The contrast transfer function (CTF) parameters were calculated using Gctf (Zhang, 2016). Gautomatch-v0.53 (https://www.mrc-lmb.cam.ac.uk/kzhang/Gautomatch/) was used for automated particle picking on the remaining 6682 micrographs after manually discarding those containing contamination, no particles, significant drift or damaged holes. RELION 3.0-alpha was used for the entire data processing (Nakane et al., 2018; Scheres, 2012a, b; Zivanov et al., 2018). 632,000 particles were extracted with binning-factor 2 and submitted for four rounds of 2D classification from which 472,000 particles selected particles were unbinned in box size of 360 pixels and submitted for 3D classification using a regularization parameter (T) of 4. Four of eight good 3D classes were included in further processing (Figure S2).

Two classes containing the best Csm3/Tof1 density (nearly 300,000 particles, 60%) were combined and 3D-refined before performing further rounds of CTF refinement, Bayesian polishing (Zivanov et al., 2019) and 3D-refinement to yield a map at an overall 3.1 Å resolution (all resolutions hereafter calculated with Gold standard Fourier shell correlation of 0.143). The precise pixel size of 1.049 Å was determined after maximising the cross-correlation coefficient between our 3.1 Å map and the CMG:DNA model (PDB ID: 5U8S) using Chimera (Pettersen et al., 2004). This pixel size was then used for postprocessing all maps obtained in this data set. To further improve the Csm3/Tof1 density, multi-body refinement (Nakane et al., 2018) was performed for (i) the Tof1 Head including N-tier regions of Mcm2, 3, 5, and (ii) the Tof1 Body/Csm3 including the N-tier regions of Mcm4, 6, 7 and dsDNA (Figure S2). Resulting maps were sharpened with B-factor −20 Å^2^ to give final maps of 3.3 and 3.2 Å resolution, respectively. These maps were used for building the models of Csm3, Tof1 and dsDNA.

For the remainder of the complex, the above 3D classification identified two conformations differing in the position of the MCM C-tier and bound ssDNA (conformations 1 and 2). One class represented complexes in conformation 1 and containing Csm3/Tof1 (124,000 particles; 26%). One class represented complexes in conformation 2 and containing Csm3/Tof1 (159,000 particles, 34%). A third class represented a mixed population of particles in both conformations, lacking clear density for Csm3/Tof1; this class was separated into conformation 1 (74,000 particles; 16%) and conformation 2 (23,000 particles; 5%) using 3D subclassification without alignment.

For conformation 1 (Figure S2, grey maps), all classes in this conformation were combined irrespective of Csm3/Tof1 occupancy (198,000 particles; 42%) and 3D-refined, before performing CTF refinement, Bayesian polishing, another round of 3D refinement and map sharpening to yield a map at 3.2 Å resolution. Multi-body refinement was performed masking more rigidly-associated regions of the complex as described in Figure S2. After map sharpening, the resulting maps were used to build the atomic models of CMG and Ctf4 for conformation 1. The above multi-body refinement maps were sharpened with the following B-factors: −40 Å^2^ for the Mcm2356 map, −5 Å^2^ for the Mcm47 map, −20 Å^2^ for the remaining bodies.

For conformation 2 (Figure S2, yellow maps), a similar approach was taken as for conformation 1. After multi-body refinement, fitting of models to the conformation 2 density confirmed the only major differences between conformations was the position of the C-tier and bound ssDNA. Consequently, the maps for the MCM C-tier, Mcm3467 and Mcm25 were sharpened with B-factors −20, −10 and −10 Å^2^ respectively, and used to build the model of the MCM C-tier in conformation 2.

After model building, it was clear there was a mixed population differing in AMP-PNP occupancy. To resolve these populations, the good particles from the original 3D-classification were combined before performing a further round of 3D-subclassification using a higher value T of 10 and limiting the Fourier components used in alignment to 10 Å resolution. Of 12 classes, one represented complexes in conformation 2 with five AMP-PNP molecules bound to the C-tier (Figure S2, magenta map), and a second with particles in conformation 2 with three AMP-PNP molecules bound and a shorter region of ssDNA resolved (Figure S2, cyan map). Models were fitted to these and the AMP-PNP occupancy and ssDNA length adjusted accordingly. This 3D-subclassification additionally yielded a 3.7 Å resolution sharpened map of conformation 1 with more homogenous resolution across all subunits in the complex.

Cryo-EM density maps best illustrating regions of unassigned density as presented in Figure 2A and Figure S5D were produced as follows. To produce the map presented in Figure 2A, the subset of particles in conformation 2, which produced the 3.3 Å map of the whole complex (see Figure S2, yellow map), was subjected to a further round of 3D sub-classification without alignment, this time utilising a higher value T of 100 in addition to providing a mask which encompassed Csm3/Tof1, dsDNA and the N-tier regions primarily belonging to Mcm4 and 6. Of six classes, four classes (50% of input particles) contained good Csm3/Tof1 and dsDNA density; these were recombined, 3D-refined and finally sharpened with a B-factor of −5 Å^2^.

To produce the map presented in Figure S5D, again using the 3.3 Å map mentioned above for conformation 2 as a starting point, signal subtraction was performed to leave primarily regions of unassigned density described in Figure 2A. The resulting signal-subtracted particles were subjected to 3D sub-classification without alignment, using regularisation parameter T of 10 and providing a mask over the regions remaining after signal-subtraction. One of the six classes with the best-defined features (23% of input particles) was selected, and the particles within this class were reverted to their pre-signal-subtraction state. These particles were 3D refined before being subjected to a further round of 3D sub-classification without alignment, this time using a higher value T of 200 in addition to providing a mask encompassing the Tof1 Body and Csm3. Of six classes, three (76% of input particles) were combined and 3D refined. The resulting refined map was then subjected to multi-body refinement, treating the complex as two bodies encompassing either the C-tier regions primarily of Mcm4 and 6 in addition to the region of low-resolution density observed beside Mcm6 C-tier at high map thresholds, or else the remainder of the complex. The former was sharpened with a B-factor of −5 Å^2^ to yield a map at 4.5 Å resolution, and is presented in Figure S5D. The local resolution was calculated using RELION and maps colored accordingly with Chimera (Pettersen et al., 2004).

### Data processing and 3D-reconstruction for the co-expressed sample

A total of six datasets totalling 11,647 raw movies were collected and processed independently using first RELION-2.1 and then RELION 3.0-alpha (Scheres, 2012a, b; Zivanov et al., 2018). In general, raw movies were aligned and dose-weighted by MotionCor2(Zheng et al., 2017) and CTF parameters were estimated using Gctf (Zhang, 2016). Poor micrographs (containing contamination, no particles, significant drift or damaged holes) were manually excluded from each data set. All particles were picked using Gautomatch (https://www.mrc-lmb.cam.ac.uk/kzhang/Gautomatch/). After one to two rounds of 2D-classification, followed by 3D-classification and 3D-refinement, the sharpened maps for the best classes from all six data sets were compared in Chimera in order to calculate scaling factors necessary for combining the data sets initially acquired at different microscope magnifications. Refined particles from five data sets were rescaled to the relative pixel size of the sixth dataset at 1.11 Å/pixel after re-estimation of the CTF parameters followed by particle re-extraction using a box size of 360 pixels (Wilkinson et al., 2019). The combined dataset comprised 412,000 particles, which were submitted for 3D-refinement. The resulting 3.4 Å map was sharpened with a B-factor of −20 Å^2^ and is presented in Figure S 5a (see also Figure S3E,F). To improve the resolution of the complex, a three-body multi-body refinement was performed for MCM C-tier, Csm3-Tof1-dsDNA and the remainder; the sharpened maps are presented in Figure S3G.

The maps with additional unassigned density beside Mcm6 and Cdc45 (Figure S5B,C) were calculated after focused classification with signal subtraction (Bai et al., 2015). Two regions were identified after closer inspection of the map obtained from the combined dataset (that presented in Figure S3G); fragmented lower-resolution density was observed beside Mcm6, Ctf4 and Cdc45. Two masks were created covering these regions with one mask covering more of Mcm6 and the other one including additional density around Cdc45. After subtracting the majority of signal from refined particles in the combined dataset, except for regions defined within these masks, focused classification was performed without alignment and with the regularisation parameter T set to 10. Two classes with additional density by the side of Mcm6 (41,338 particles, 10% of input) or over Cdc45 (74,641 particles, 18% of input) emerged from these 3D sub-classifications and were selected and then reverted to particles with original signal before being subjected to 3D refinement. The refined map with additional globular density beside Mcm6 was sharpened with a B-factor of −20 Å^2^ at an overall resolution of 4.5 Å (Figure S5B). A more extended density on Cdc45 became clearer in map refined to 4.2 Å resolution also sharpened with B-factor of −20 Å^2^ (Figure S5C).

### Model building and refinement

Model building was carried out in *Coot* (Emsley et al., 2010) using maps generated by multi-body refinement, as detailed in the data processing section for the reconstituted sample. The summary of proteins and DNA is given in Table S2. An atomic model was built for conformation 1. As initial template models, CMG:DNA (PDB ID: 5U8S) (Georgescu et al., 2017) and the crystal structure of the C-terminal regions of Ctf4 (PDB ID: 4C8H) (Simon et al., 2014) were used where individual subunits were fitted as rigid bodies to the relevant multi-body refinement maps using Chimera (UCSF) (Pettersen et al., 2004), with N- and C-tier regions of MCM subunits fitted separately. These subunits were then morphed and jiggle-fitted to the relevant maps in *Coot*, prior to adjusting the models to density manually using local refinement. Once rebuilt, individual subunits were refined in the relevant maps using either Refmac (Kovalevskiy et al., 2018), phenix.real_space_refine (Afonine et al., 2018) or ISOLDE (Croll, 2018). It was notable that the resolution of MCM C-tier subunits was variable, with Mcm2, 3, 5 and 6 (those binding AMP-PNP and ssDNA) better resolved than Mcm4 and 7.

For Mcm subunits, the regions N-terminal to the helical domain (N-terminal extension, NTE) of Mcm2, 4 and 6 were extended, with 28 residues built for the Mcm2 NTE, 12 residues built for the Mcm6 NTE, and remodelling of 6 residues of the Mcm4 NTE. These NTE regions contain Tof1 binding sites. 172 residues were not observed for the NTE of Mcm2, although unassigned density is present in the vicinity and may account for some of these residues.

The Zinc-finger (ZnF) domains were rebuilt with tetrahedral-coordinated Zn^2+^ ions placed in the spherical density that was observed at low contour level between four cysteine residues in each of ZnF domains in Mcm4, 6 and 7. The Cα-backbone and the sequence register of Mcm5 ZnF was remodelled using the MCM double hexamer template model (PDB ID: 5BK4) (Noguchi et al., 2017). The N-terminal hairpin (NTH) loops of Mcm7 (the separation pin, 362-368) and Mcm2 (436-443) were remodelled as *α*-helix. The linkers between N- and C-tier were built fully as loops for Mcm2 (459-476) and 5 (339-363) and an *α*-helix (501-508) was built for mostly disordered N/C-tier linker of Mcm6 (463-509). In the C-tier, the ssDNA-binding regions (helix H2, H2I loops and PS1 loops) showed much improved density, which required significant remodelling in terms of repositioning and extending Cα-backbone, assigning correct sequence register and rebuilding of side-chains.

The observed density in C-tier region of Mcm3 (vicinity of residue 583) remains unassigned (584-647). Additional helical density observed in the vicinity of Mcm6 could be potentially attributed to residues 202-251 and/or its N/C-tier linker, however this region was not included in the final model. The C-terminal winged-helix (WH) domain (851-877) was retained for Mcm4 in the lower-resolution density, as seen in prior structures. AMP-PNP/Mg^2+^ was built in a well resolved density at the Mcm2/6, 2/5 and 3/5 interfaces with side chains visible for WalkerA, WalkerB, Arg-finger and Sensor2 motifs (Figure S1G).

For the remainder of CMG, the model for the N-terminal CIP-box of Sld5 was rigid body fitted and adjusted in clearly visible density using the template coordinates (PDB ID: 4C95) (Simon et al., 2014). The model for Psf2 was extended for additional residues 33-38, which now appear to be ordered, presumably through the interaction of this region with the Ctf4 helical bundle. For Ctf4, the side-chains were adjusted, particularly at the interface with Cdc45 and Psf2. The N-terminal regions of Ctf4 (1-460), known to contain a WD40 domain in human And-1, could not be assigned to particular regions of density in our complex. Five residues of the Psf3 N-terminal His-tag were resolved in the density and are present in the model (N-Ser-His-Met-Ala-Ser-C).

For conformation 2, the largest changes were observed in the MCM C-tier and length of bound ssDNA, therefore a model was built for this region by adjusting our MCM C-tier models for conformation 1 to density of conformation2. The resolution of C-tier subunits varied, with those bound to AMP-PNP/Mg^2+^ and 15-mer ssDNA (built as poly-dT) better resolved (the AMP-PNP-free interface only observed between Mcm2 and 5). The major differences compared to conformation 1 were observed in the relative positions of individual MCM subunits and the positions of the ssDNA-binding loops, particularly ordered interfaces between H2I side chains and sugar/bases of ssDNA were rebuilt for Mcm2, 6, 4, 7 and 5. For Mcm4, density for the WH was no longer observed, while the linker connecting the WH to the AAA+ domain was repositioned away from the MCM central channel.

The Csm3/Tof1 dimer was partially built *de novo*. The N-terminus of Tof1 was identified in our density after rigid-body fitting of the fragment of human Timeless (PDB ID: 5mqi) (Holzer et al., 2017), which was then used for homology modelling of the Tof1 region comprising helical repeats 1-6 using Phyre2 (Kelley et al., 2015). The Tof1 homology model was morphed and jiggle-fitted into our multi-body map of Tof1 Head (Figure S2), which was then manually adjusted and locally refined before building *de novo* the Ω-shaped loop (Ω-loop) and the long MCM-plugin, which extend between helical repeats 3-4 and 4-5, respectively. The Mcm6 NTE packs against the Ω-loop and this region was built as a composite Mcm6-Tof1 β-sheet given good density. The density for the Ω-loop in the region facing the major groove of DNA indicates greater flexibility, however three lysine residues (K225, K226 and K230) could be built into the density. The density and connectivity for the long MCM-plugin was of a good quality, in particular several prominent newly built secondary structure elements (helices Bridge and αW, and the β-hairpin, see Figure S9E) packing against the interface with Mcm6, 4 and 7. The density of MCM-plugin, which protrudes into the minor groove of DNA could be well resolved and the side chains built represent those of the Tof1 DBM (401-404). Repeats 7-9 of Tof1 (the Body encompasses repeats 6-9, Figure S9A) were built *de novo* by first placing idealized alpha helices, which generally matched well the secondary structure prediction for this region (Phyre2). The side chains were next built manually in the density up to residue 781 with certain loops being omitted due to lack of density (Table S2). Following residue 781, the remaining novel density accounting for five helices was observed to have opposite polarity to helices in the Body and the model for this density was built *de novo* by first placing idealised helices. The sequence register was assigned to the core of Csm3; in particular, the side chains of the helix α2 were well resolved with a prominent tryptophan side chain (W98). Given the continuous density between these helices the sequence register for the remaining of Csm3 was assigned with confidence (helices α0, α1, α3 and α4). The secondary structure prediction for Csm3 (Phyre2) was in line with the experimental structure. Additional density extending from a small helix α0 into the minor groove of dsDNA was built as the Csm3 DBM (46-53). This region is predicted to be disordered and is likely stabilised by interaction with DNA.

The dsDNA was built in the density of a body representing Tof1-Head/Csm3/dsDNA by fitting idealised B-form DNA. Sequence register was assigned based on the sequence of our fork DNA.

### Model to cryo-EM map validation for conformation 1 and conformation 2

Fourier Shell Correlation (FSC) was calculated between the refined replisome complex models (conformation 1 and MCM C-tier:ssDNA of conformation 2) and their respective unsharpened sums of the two half maps using XMIPP (Sorzano et al., 2004). The above models were also refined with restraints against the respective half-1 maps and the FSC map to model curves were calculated for the half-1 and half-2 (not used for model refinement) maps (Figure S1I).

### Cross-linking mass spectrometry (XL-MS)

The complex comprising CMG, Ctf4, Csm3/Tof1 and Mrc1 was purified following co-expression as described above. The eluate from the Calmodulin sepharose 4B column (25 mM HEPES pH 7.5, 150 mM sodium acetate, 0.5 mM TCEP, 2 mM EDTA/EGTA) was immediately cross-linked with a 100-fold excess of the N-hydroxysuccinimide (NHS) ester disuccinimidyl dibutyric urea (DSBU, ThermoScientific, USA), with respect to the protein concentration. The cross-linking reactions were incubated for 60 min at room temperature and then quenched by the addition of NH_4_HCO_3_ to a final concentration of 20 mM and incubated for further 15 min. The cross-linked proteins were then precipitated according to the method of Wessel and Flugge (Wessel and Flugge, 1984) and resuspended in 8 M urea in 50 mM NH_4_HCO_3._

The cross-linked proteins were reduced with 10 mM DTT and alkylated with 50 mM iodoacetamide. Following alkylation, the concentration of urea was reduced to 1 M by the addition of 50 mM NH_4_HCO_3_ and the proteins digested with trypsin (Promega, UK) at an enzyme-to-substrate ratio of 1:100, for 1 h at room temperature and then further digested overnight at 37 °C following a subsequent addition of trypsin at a ratio of 1:20.

The peptide digests were then fractionated batch-wise by high pH reverse phase chromatography on micro spin C18 columns (Harvard Apparatus, USA), into five fractions (10 mM NH_4_HCO_3_ /10 % v/v acetonitrile pH 8, 10 mM NH_4_HCO_3_ /20 % v/v acetonitrile pH 8, 10 mM NH_4_HCO_3_ /30 % v/v acetonitrile pH 8, 10 mM NH_4_HCO_3_ /50 % v/v acetonitrile pH 8 and 10 mM NH_4_HCO_3_ /80 % v/v acetonitrile pH 8). The 150 µl fractions were evaporated to dryness on a CoolSafe lyophilizer (ScanVac, Denmark) prior to analysis by LC-MS/MS.

Lyophilized peptides for LC-MS/MS were resuspended in 0.1 % v/v formic acid and 2 % v/v acetonitrile and analyzed by nano-scale capillary LC-MS/MS using an Ultimate U3000 HPLC (ThermoScientific Dionex, USA) to deliver a flow of approximately 300 nl/min. A C18 Acclaim PepMap100 5 µm, 100 µm × 20 mm nanoViper (ThermoScientific Dionex, USA), trapped the peptides before separation on a C18 Acclaim PepMap100 3 µm, 75 µm × 250 mm nanoViper (ThermoScientific Dionex, USA). Peptides were eluted with a gradient of acetonitrile. The analytical column outlet was directly interfaced via a nano-flow electrospray ionisation source, with a quadrupole Orbitrap mass spectrometer (Q-Exactive HFX, ThermoScientific, USA). MS data were acquired in data-dependent mode using a top 10 method, where ions with a precursor charge state of 1+ and 2+ were excluded. High-resolution full scans (R=120 000, m/z 300-1800) were recorded in the Orbitrap followed by higher energy collision dissociation (HCD) (stepped collision energy 26 and 28 % Normalized Collision Energy) of the 10 most intense MS peaks. The fragment ion spectra were acquired at a resolution of 50,000 and dynamic exclusion window of 20 s was applied.

For data analysis, Xcalibur raw files were converted into the MGF format using MSConvert (Proteowizard) (Kessner et al., 2008) and used directly as input files for MeroX (Gotze et al., 2015). Searches were performed against an *ad hoc* protein database containing the sequences of the proteins in the complex and a set of randomized decoy sequences generated by the software. The following parameters were set for the searches: maximum number of missed cleavages 3; targeted residues K, S, Y and T; minimum peptide length 5 amino acids; variable modifications: carbamidomethylation of cysteine (mass shift 57.02146 Da), Methionine oxidation (mass shift 15.99491 Da); DSBU modified fragments: 85.05276 Da and 111.03203 Da (precision: 5 ppm MS and 10 ppm MS/MS); False Discovery Rate cut-off: 5 %. Finally, each fragmentation spectrum was manually inspected and validated.

### Origin-dependent DNA replication assays

Origin-dependent replication assays were performed essentially as described previously (Aria and Yeeles, 2019; Taylor and Yeeles, 2018). MCM loading was performed at 24 °C in reactions (typically 35 μl) containing 25 mM HEPES-KOH pH 7.6, 100 mM K-glutamate, 0.01% v/v Nonidet P40 substitute (NP-40-S) (Roche #11754599001), 1 mM DTT, 10 mM Mg(OAc)_2_, 40 mM KCl, 0.1 mg/ml BSA, 3 mM ATP, 3 nM AhdI-linearized vVA20 template(Aria and Yeeles, 2019), 75 nM Cdt1•Mcm2-7, 40 nM Cdc6, 25 nM DDK, 20 nM ORC. After 10 min S-CDK was added to a final concentration of 80 nM and incubation continued at 24 °C for 5 min. Reactions were diluted 4-fold into replication buffer to give final reactions concentrations (accounting for subsequent addition of replication proteins) of 25 mM HEPES-KOH pH 7.6, 250 mM K-glutamate, 0.01% NP-40-S, 1 mM DTT, 10 mM Mg(OAc)_2_, 10 mM KCl, 0.1 mg/ml BSA, 3 mM ATP, 200 μM C/G/UTP, 30 μM dA/dT/dG/dCTP, 1 µCi [α-^32^P]-dCTP, 0.75 nM AhdI-linearized vVA20 template, 18.75 nM Cdt1•Mcm2-7, 10 nM Cdc6, 6.25 nM DDK, 5 nM ORC. Reactions were equilibrated at 30 °C (∼1 min) and replication initiated by addition of replication proteins from a master mix to the following final concentrations: 30 nM Dpb11, 100 nM GINS, 30 nM Cdc45, 10 nM Mcm10, 20 nM Pol ε, 20 nM Ctf4, 100 nM RPA, 20 nM RFC, 20 nM PCNA, 20 nM Pol α, 10 nM Pol δ, 12.5 nM Sld3/7, 20 nM Sld2, 20 nM Mrc1 and 20 nM Csm3/Tof1 or mutants where indicated. Reactions were quenched by addition of an equal volume of 100 mM EDTA and samples were processed as previously described (Aria and Yeeles, 2019; Taylor and Yeeles, 2018). RFB experiments were performed on vJY30 (Table S4) and Fob1 was added together with the MCM loading proteins to a final concentration of 250 nM.

### Replisome association assays

MCM loading was performed at 30 °C in reactions containing 25 mM HEPES-KOH pH 7.6, 100 mM K-glutamate, 0.01% v/v NP-40-S, 1 mM DTT, 10 mM Mg(OAc)_2_, 0.1 mg/ml BSA, 3 mM ATP, 3 nM vVA20 template(Aria and Yeeles, 2019), 75 nM Cdt1•Mcm2-7, 40 nM Cdc6, 25 nM DDK, 14 nM ORC. After 30 min reactions were diluted 2-fold into replication buffer to give final reactions concentrations (accounting for subsequent addition of replication proteins) of 25 mM HEPES-KOH pH 7.6, 250 mM K-glutamate, 0.01% NP-40-S, 1 mM DTT, 10 mM Mg(OAc)_2_, 0.1 mg/ml BSA, 3 mM ATP, 200 μM C/G/UTP, 30 μM dA/dT/dG/dCTP, 1.5 nM vVA20 template, 37.5 nM Cdt1•Mcm2-7, 20 nM Cdc6, 12.5 nM DDK, 7 nM ORC. Replication was initiated by addition of replication proteins from a master mix to the following final concentrations: 30 nM Dpb11, 100 nM GINS, 30 nM Cdc45, 10 nM Mcm10, 20 nM Pol ε, 20 nM Ctf4, 100 nM RPA, 20 nM RFC, 20 nM PCNA, 20 nM Pol α, 12.5 nM Sld3/7, 20 nM Sld2, 10 nM Mrc1 and 20 nM Csm3/Tof1 or mutants where indicated. After 25 min, samples (13 µl) were directly applied to 400 µl (bed volume) Sephacryl S-400 columns (GE healthcare) equilibrated in 25 mM HEPES-NaOH pH 7.5, 150 mM NaOAC, 10 mM Mg(OAc)_2_, 1 mM DTT, 0.01% v/v NP-40-S and 0.1 mM ATP. Columns were centrifuged (750g, 2 min, 21°C) and the eluate was analysed by SDS-PAGE and western blotting. Cdc45 was detected using its FLAG epitope with an anti-FLAG antibody (A8592, Sigma). RPA was detected with an antibody against the Rpa1 subunit (AS07 214). Mcm7, Psf1, Ctf4, Csm3 and Mrc1 were detected with sheep monoclonal antibodies (Maric et al., 2014; Mukherjee and Labib, 2019).

### Electrophoretic mobility-shift assays

Csm3/Tof1 wild-type or mutant proteins were mixed with fork DNA prepared as for cryo-EM sample preparation (40 nM final [DNA]), at a molar ratio of protein:DNA of 1:1, 2:1, 4:1 and 8:1 in a reaction buffer containing 25 mM HEPES-KOH, pH 7.6, 100 mM KOAc, 2 mM Mg(OAc)_2_, 0.2 % NP40 and 1 mM DTT and incubated on ice for 30 min. Ficoll 400 was added to each 15 μL reaction to a final concentration of 2.3 % v/v before loading onto 4% native polyacrylamide gels for analysis. Gels were imaged on a using a Typhoon fluorescence imager (Amersham) at the Cy3 excitation wavelength of 532 nm.

### Camptothecin sensitivity assays

Saturated cultures of *S. cerevisiae* grown in YEP + 2% w/v glucose were diluted to an A_600_ of 0.2 and were grown to an A_600_ of ∼0.6-0.8 in YEP + 2% w/v glucose at 30°C. Cells were harvested and resuspended in YEP + 2% w/v glucose + 100 μg/ml ampicillin or sterile water to a A_600_ of 0.5. Cells from a 10-fold serial dilution in YEP + 2% w/v glucose + 100 μg/ml ampicillin or sterile water were then plated (8 μl) on YEPD agar plates supplemented with either DMSO or 20 μM camptothecin (Merck or Santa Cruz biotechnology). Plates were incubated at 25°C for 48 hours.

### Multiple sequence alignments

Amino acid sequences were retrieved from relevant databases (NCBI or SGD where stated; UniProt if stated or otherwise) (Cherry et al., 2012; UniProt, 2019). Alignment was performed using MUSCLE (EMBL-EBI) (Edgar, 2004). The alignment was rendered using Espript3.0 (https://espript.ibcp.fr) (Robert and Gouet, 2014).

### Structural analysis and visualization

All figures of structures were plotted in PyMOL (Schrodinger, 2015), Chimera (Pettersen et al., 2004) or ChimeraX (Goddard et al., 2018). Interaction analysis between CMG and Ctf4 was performed using PISA (Krissinel and Henrick, 2007).

