## Supplemental Information for "Cryo-EM Structure of the Fork Protection Complex Bound to CMG at a Replication Fork"

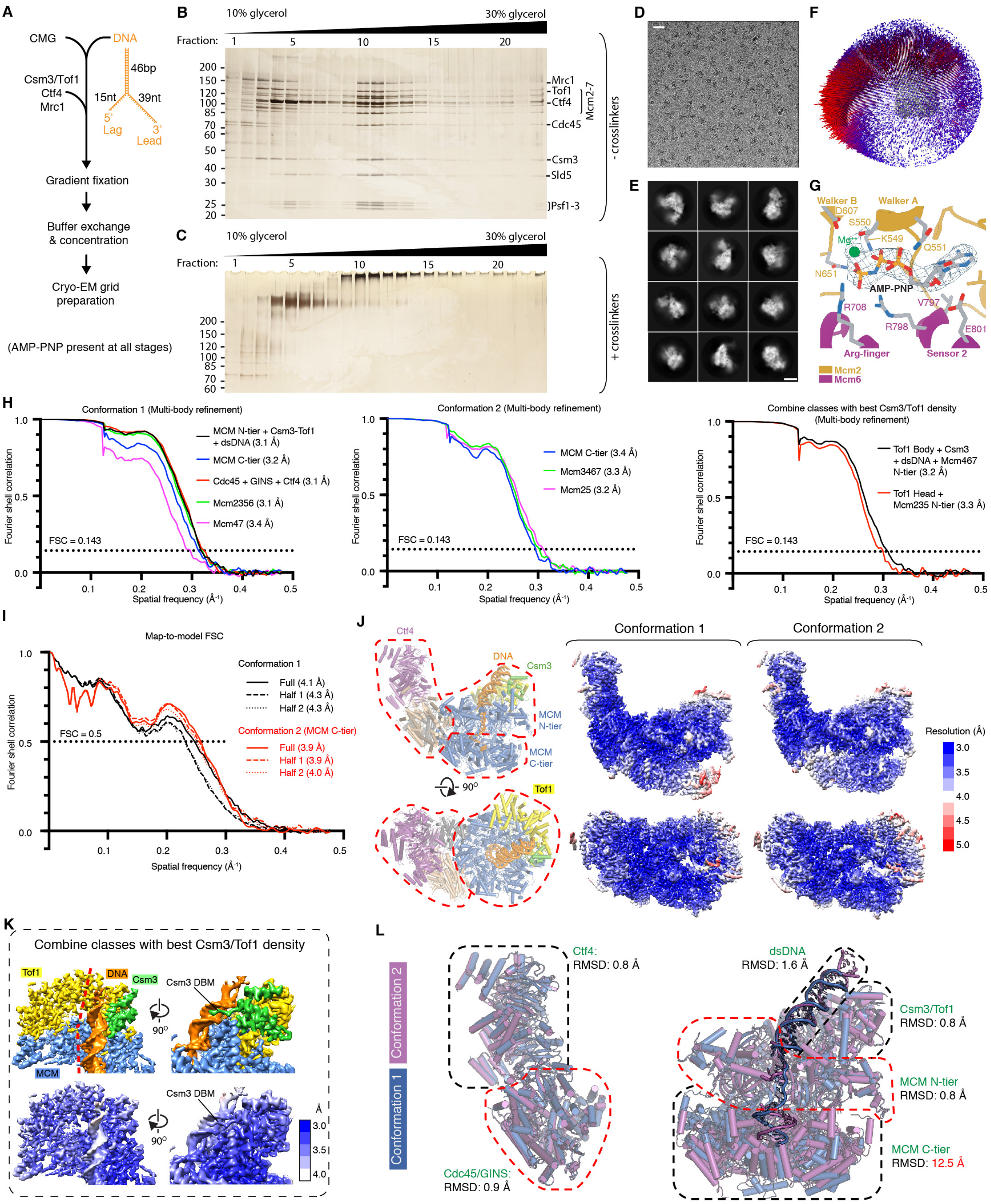

**Figure S1. Samples for cryo-EM experiments prepared by *in vitro* reconstitution**

(A) Schematic illustrating method of sample preparation for cryo-EM experiments.

(B, C) Representative silver-stained SDS-PAGE showing fractions taken across glycerol gradients prepared without (B) and with (C) crosslinkers (bis[sulfosuccinimidyl] suberate and glutaraldehyde). Crosslinked fractions 10+11 were pooled for cryo-EM grid preparation.

(D) Representative cryo-EM micrograph. Scale bar, 30 nm.

(E) Representative 2D class averages. Scale bar, 10 nm.

(F) Representative 3D angular distribution of particle projections.

(G) Representative example of cryo-EM density observed for a bound AMP-PNP•$Mg^{2+}$.

(H) Representative Fourier shell correlation (FSC) curves shown for maps used in model building. The FSC=0.143 criterion used to determine map resolution is indicated on all graphs (dotted black line). Left: conformation 1 (grey maps in Figure S2). Centre: conformation 2 (yellow maps in Figure S2). Right: maps used to build Csm3/Tof1 models (red maps in Figure S2).

(I) Model-to-map FSC curves. Full denotes the correlation between the sum of half1- and half2-maps and refined model. Half 1 denotes correlation between half1-map and model refined in half1-map. Half2 stands for correlation between half2-map and model refined in half1-map.

(J) Representative cryo-EM density maps for conformations 1 and 2, colored by local resolution. A model (conformation 1) is shown on the left for reference. For each of conformation 1 and 2, a composite map is shown comprising three separate regions treated as individual bodies in multi-body refinement, their boundaries roughly illustrated by red dashed lines (left) (refer to Figure S2, grey and yellow colored maps).
(K) Cryo-EM density maps encompassing Csm3/Tof1 and dsDNA, colored by subunit (top) and local resolution (bottom). Displayed is a composite of two maps generated by multi-body refinement, encompassing either the Tof1 Head or Tof1 Body, Csm3 and dsDNA, plus underlying regions of MCM subunits (see Figure S2, red color maps). The rough demarcation of these maps is illustrated by red dashed lines (top left).
(L) Comparison of conformations 1 and 2. Models were built for the entire complex in conformation 1, and the MCM C-tier of conformation 2. To complete the model of conformation 2, remaining subunits were docked as rigid bodies with models fitting the density well. Complete models for conformations 1 and 2 were then aligned on either the MCM N-tier or Cdc45/GINS (red outlines) and root mean square deviation (RMSD) values calculated for the regions indicated (dashed outlines) without further alignment.

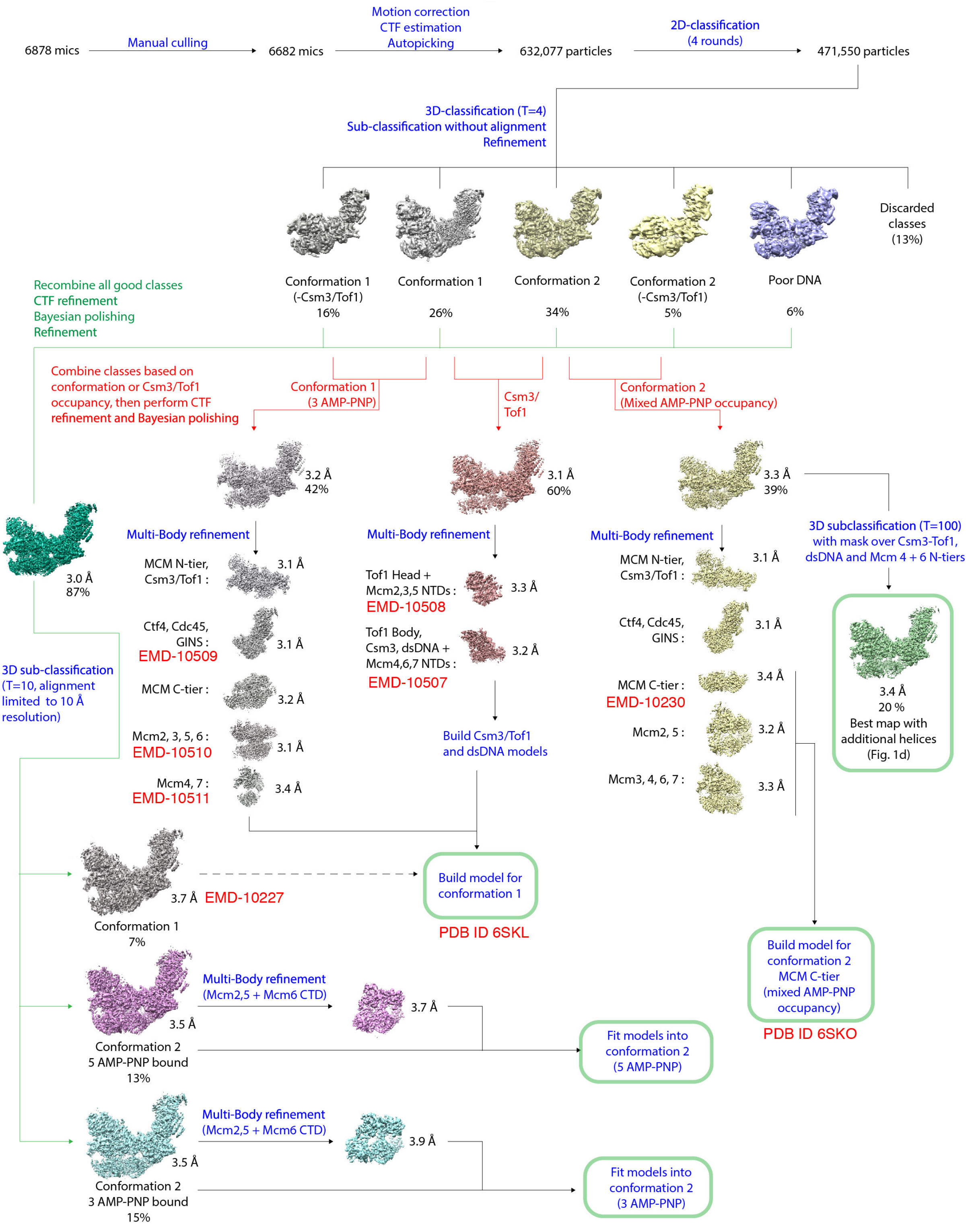

**Figure S2. Schematic of data processing pipeline for cryo-EM samples prepared by *in vitro* reconstitution**
Cryo-EM density maps related to conformation 1 are colored grey. The 3.7 Å conformation 1 map represents the map with best density over all subunits in a single map for this conformation. Maps related to conformation 2 are colored yellow for maps with a mixed AMP-PNP occupancy, or magenta/cyan for maps with five/three AMP-PNP bound. T is the regularization parameter used during 3D classification. Green boxes highlight the final models discussed in the text and the map best illustrating regions of unassigned density (Figure 2A).

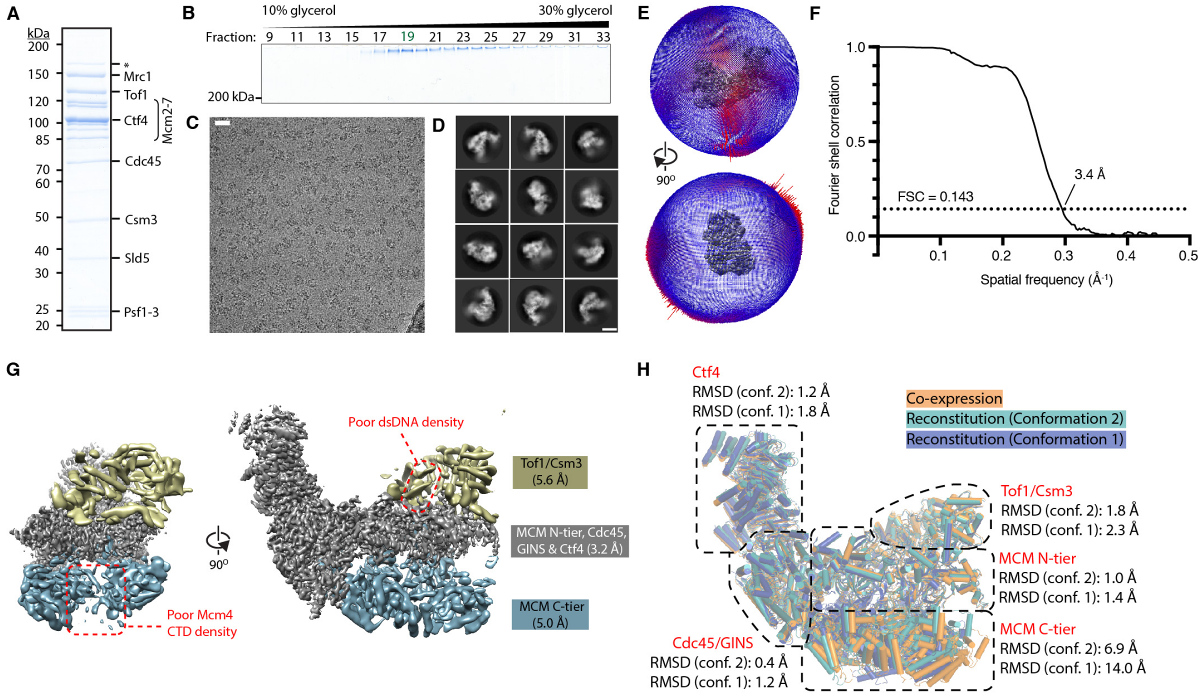

**Figure S3. Samples for cryo-EM experiments prepared by co-expression of 15 proteins in *S. cerevisiae***

(A, B) Representative Coomassie-stained SDS-PAGE of the eluate from calmodulin sepharose 4B resin (A) (see Methods for details) and fractions across a crosslinking glycerol gradient containing glutaraldehyde (B). In (B), fraction 19 was taken for cryo-EM grid preparation. In (A) the * indicates a contaminant with comparable migration to previously identified Pol1, a known interaction partner of Ctf4 shown to co-purify with replisome progression complexes under similar purification conditions (Gambus et al., 2006). Note the 14 mL gradient (400 µL fractions) used in (B) differs from the 2.2 mL gradient (100 µL fractions) used for the reconstituted sample (Figure S1B,C), explaining the discrepancy in the fraction number chosen.

(C) Representative cryo-EM micrograph. Scale bar, 30 nm.

(D) Representative 2D class averages. Scale bar, 10 nm.

(E) 3D angular distribution for particle projections.

(F) Fourier shell correlation curve for the best overall cryo-EM map.

(G) Cryo-EM maps produced using multi-body refinement, with individually refined regions (bodies) colored. Single-stranded DNA was not observed to bind the MCM C-tier, consistent with ssDNA-binding stabilizing either of the two C-tier conformations in the reconstituted sample allowing higher-resolution reconstructions to be attained for this region in the reconstituted sample (Figure S1H,J,K).

(H) Comparison of reconstitution and co-expression datasets illustrating that major differences are only observed in the MCM C-tier. The model for the co-expressed sample was generated by fitting rigid bodies for individual subunits, with N- and C-tier regions fitted independently for each Mcm. The complete model for conformation 2 was produced as described for Figure S1L. Complete models (except DNA) were aligned on the region encompassing MCM N-tier, Cdc45 and GINS, and RMSD values were determined for each region of the complex indicated by dashed outlines without further alignment.

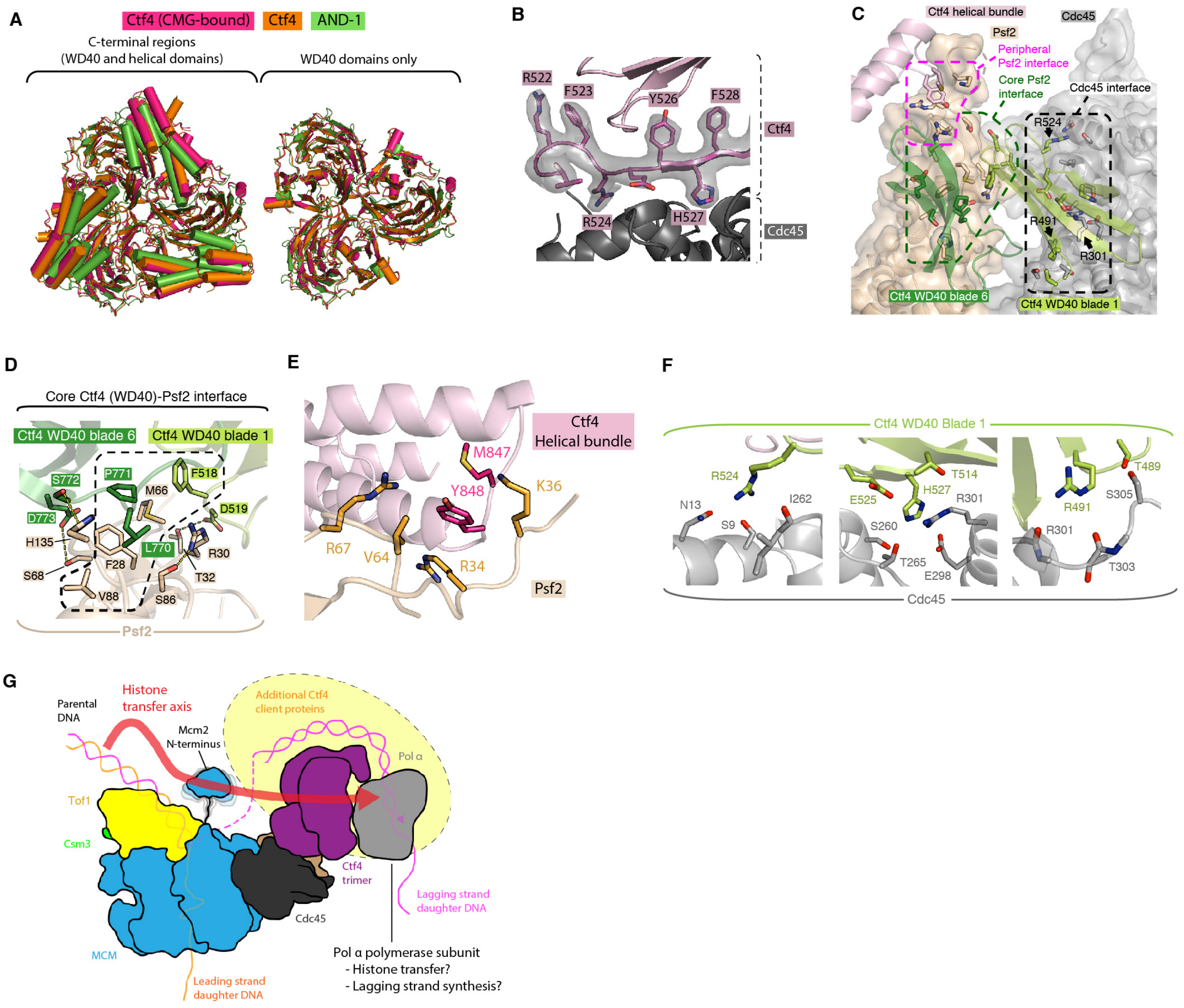

**Figure S4. Ctf4 tethering in the replisome**

**(A)** Comparison of CMG-bound Ctf4 with previously published crystal structures of the C-terminal regions of trimeric Ctf4 (PDB ID: 4C8H) (Simon et al., 2014) and human And-1 (PDB ID: 5OGS) (Kilkenny et al., 2017).

(B) Representative region of cryo-EM density at the CMG:Ctf4 interface, shown for Ctf4 WD40 blade 1.

(C) Overview of the CMG-Ctf4 interface from a "top-down" perspective looking from Ctf4 towards Cdc45/GINS. Surface representations are shown for Cdc45/GINS. Only the regions of Ctf4 involved in the interaction are displayed. Core and peripheral interaction networks at the interface with Psf2 are expanded in (D) and (E), respectively. Arginine residues central to interaction networks at the interface with Cdc45 [expanded in (F)] are labelled.

(D) Ctf4-Psf2 core interface mediated by the Ctf4 WD40 domain. The central hydrophobic network is indicated by the dashed outline. Yellow dashes indicate hydrogen bonds.

(E) Ctf4-Psf2 peripheral interaction network mediated by the Ctf4 helical bundle.

(F) Ctf4-Cdc45 interface, subdivided into three interaction networks each involving a central arginine.

(G) Schematic illustrating the position of Ctf4 in the context of the eukaryotic replisome replicating DNA. The position of Pol α suggested by recent cryo-EM investigation of CMG-Ctf4 complexes in the absence of DNA is indicated (Yuan et al., 2019). The N-terminus of Mcm2 (not resolved) is also illustrated as flexibly tethered. The yellow halo indicates additional Ctf4 interacting partners ("clients") are expected to interact with the replisome, including Chl1, Dia2 and Dna2 (Morohashi et al., 2009; Villa et al., 2016). The role suggested of Ctf4 in parental histone transfer to the lagging strand along the Mcm2-Ctf4-Pol α axis is illustrated (Gan et al., 2018).

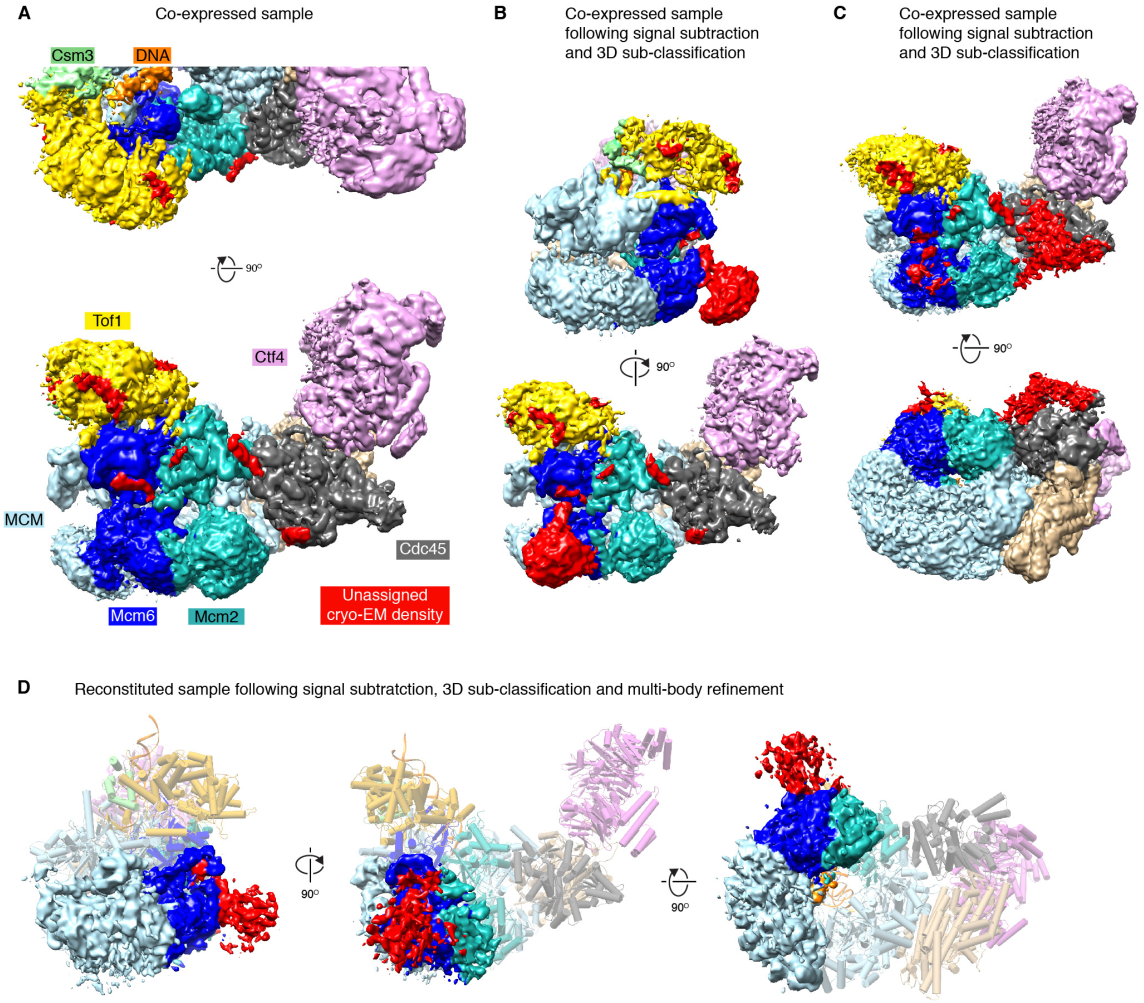

**Figure S5. Regions of unassigned density in co-expressed and reconstituted cryo-EM samples**

(A) Cryo-EM density map reconstructed for the co-expressed complex, colored according to subunit identity. Regions of unassigned density are colored red. Unassigned density is observed in regions comparable to the reconstituted sample (Figure 2A). Additional fragmented density was observed at low resolution. To improve the reconstruction of these regions, signal subtraction and focused 3D sub-classification was performed, yielding the maps displayed in panels B-D.

(B, C) Cryo-EM density maps for the co-expressed sample produced by signal subtraction and focused 3D sub-classification, revealing additional unassigned density beside Mcm6 (B) and Cdc45 (C).

(D) Cryo-EM density map for the reconstituted sample encompassing the C-tier regions of Mcm2, 6, 4 and 7, produced by signal subtraction, focused 3D-subcassification and focused refinement using multi-body refinement. Additional unassigned density is revealed beside Mcm6, as seen in co-expressed complexes (B). The model for the whole complex is shown in the background for context.

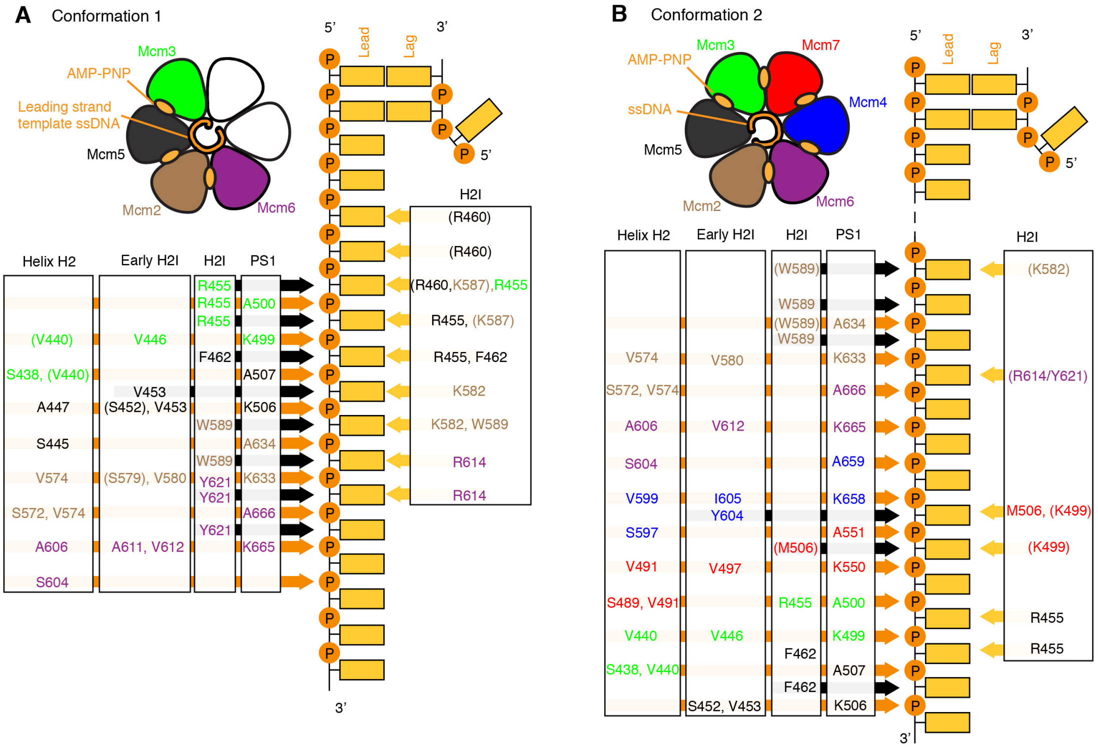

**Figure S6. Supporting figure 1 for MCM:ssDNA interactions**

(A, B) Representation of contacts made by the MCM C-tier with leading-strand template ssDNA for conformations 1 and 2 respectively. Arrows indicating interaction with phosphate ("P"), ribose and nucleobase moieties are colored orange, black and yellow, respectively. Parentheses denote residues where cryo-EM density could not unambiguously identify the position of side chains.

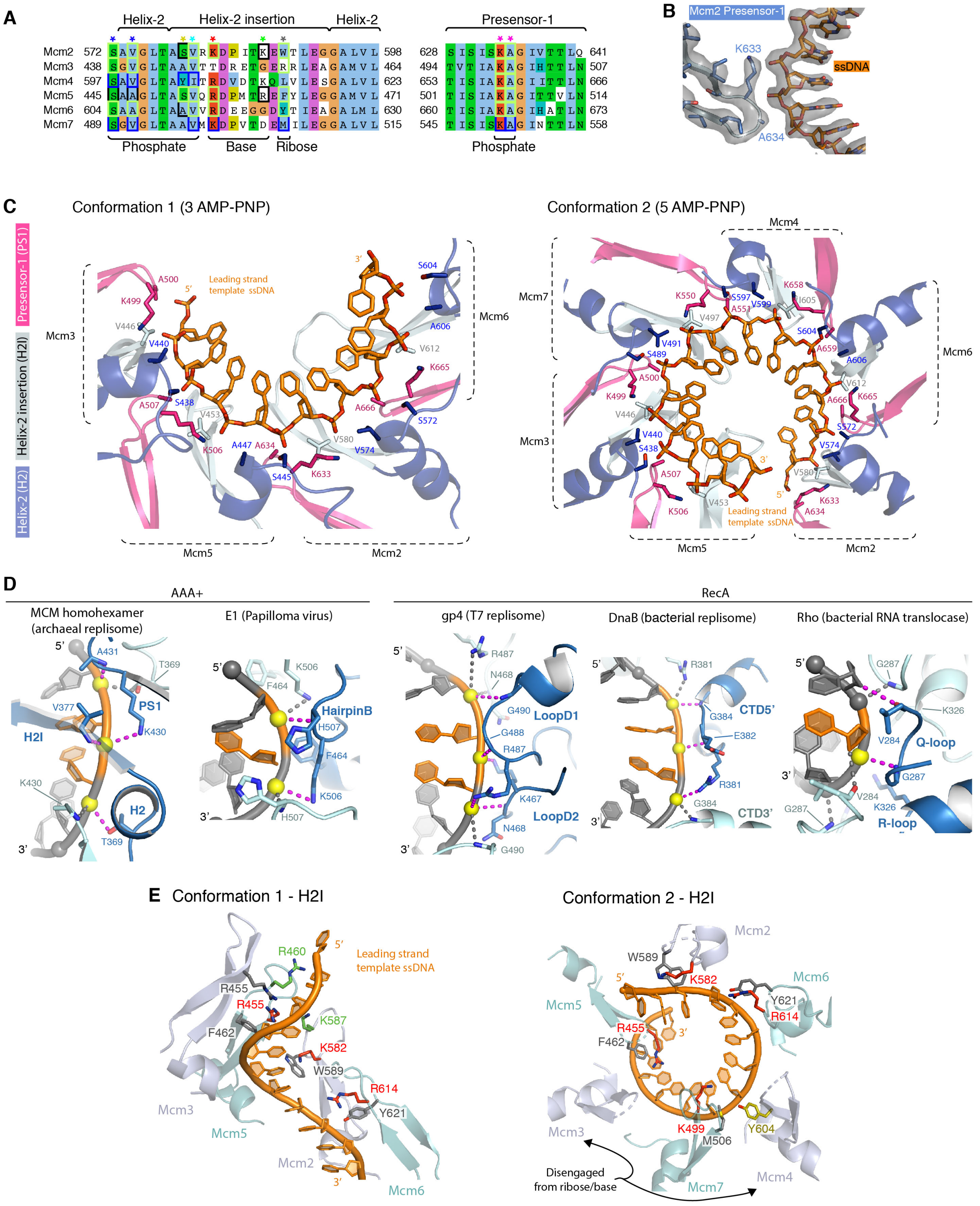

**Figure S7. Supporting figure 2 for MCM:ssDNA interactions**

(A) Multiple sequence alignment for MCM C-tier DNA binding loops for *S. cerevisiae* MCM subunits. Boxes indicate residues involved in ssDNA interactions, with interactions observed in conformation 1, 2 or both colored black, blue or lime-green, respectively. Asterisks above the alignment are colored according to the scheme used in other panels.

(B) Cryo-EM density observed for C-tier-bound ssDNA and a representative interacting loop.

(C) Repetitive interactions of MCM C-tier AAA+ domains with the leading-strand template phosphate moieties.

(D) Comparison of repetitive phosphate contacts mediated by diverse homohexameric nucleic acid helicases (PDB IDs: archaeal MCM, 6MII; E1, 2GXA; T7, 6N7N; DnaB, 4ESV; Rho, 5JJI) (Enemark and Joshua-Tor, 2006; Gao et al., 2019; Itsathitphaisarn et al., 2012; Meagher et al., 2019; Thomsen et al., 2016).

(E) Interactions of MCM H2I loop residues with sugar/base moieties of leading strand template ssDNA. Residues are colored according to their position in the H2I primary sequence, corresponding to the panel (A) asterisks. The parentheses around Mcms3 and 4 in conformation indicate these loops are disengaged from ssDNA, except for a unique tyrosine (colored yellow) at a distinct position in the Mcm4 H2I loop, which approaches the sugar-phosphate backbone of ssDNA.

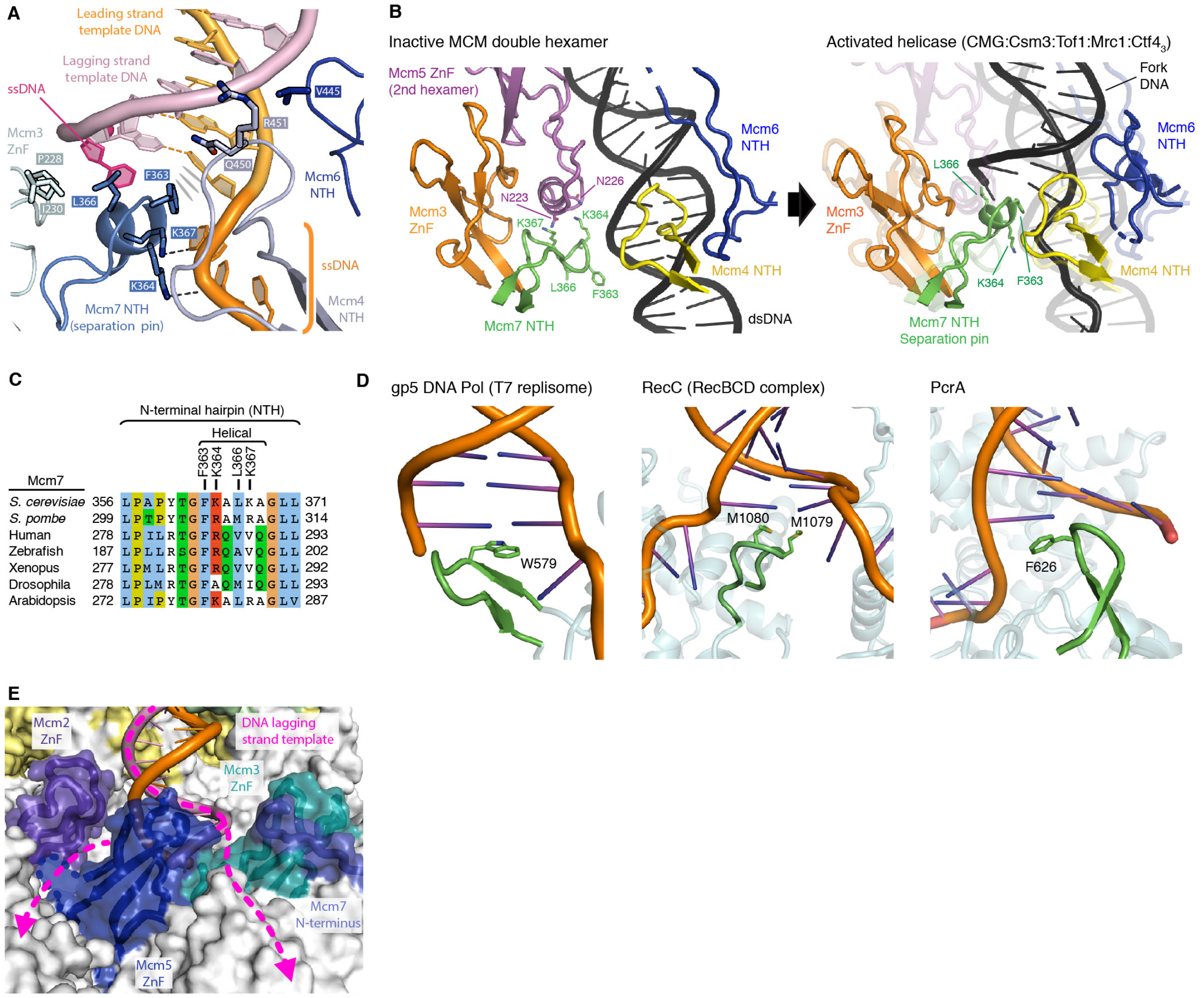

**Figure S8. Supporting figure for CMG:fork junction interactions**

(A) Detailed view of the residues involved in interactions with DNA at the fork junction. The Mcm5 ZnF which contacts the unwound lagging strand via R184 is omitted for clarity (see Movie S2). F363 is shown to make $\pi$-$\pi$ interactions with the last base-pair.

(B) Comparison of the MCM loops surrounding the fork junction in CMG with the corresponding region of the inactive MCM double hexamer on dsDNA (PDB ID: 5BK4) (Noguchi et al., 2017). Right-hand side: the model for the active helicase is shown overlaid on the model of the inactive double hexamer having aligned globally on the MCM N-tier, for ease of comparison.

(C) Multiple sequence alignment for the Mcm7 N-terminal hairpin (strand separation pin). The internal helix and DNA-interaction residues are labelled.

(D) Comparison of previously described strand separation pins from diverse helicases (PDB IDs: T7, 6N7W; RecBCD, 1W36; PcrA, 3PJR) (Gao et al., 2019; Singleton et al., 2004; Velankar et al., 1999). The characteristic hydrophobic residues observed to abut the last base-pair are indicated.

(E) Schematic for hypothesized lagging-strand template ssDNA exit pathways post-unwinding.

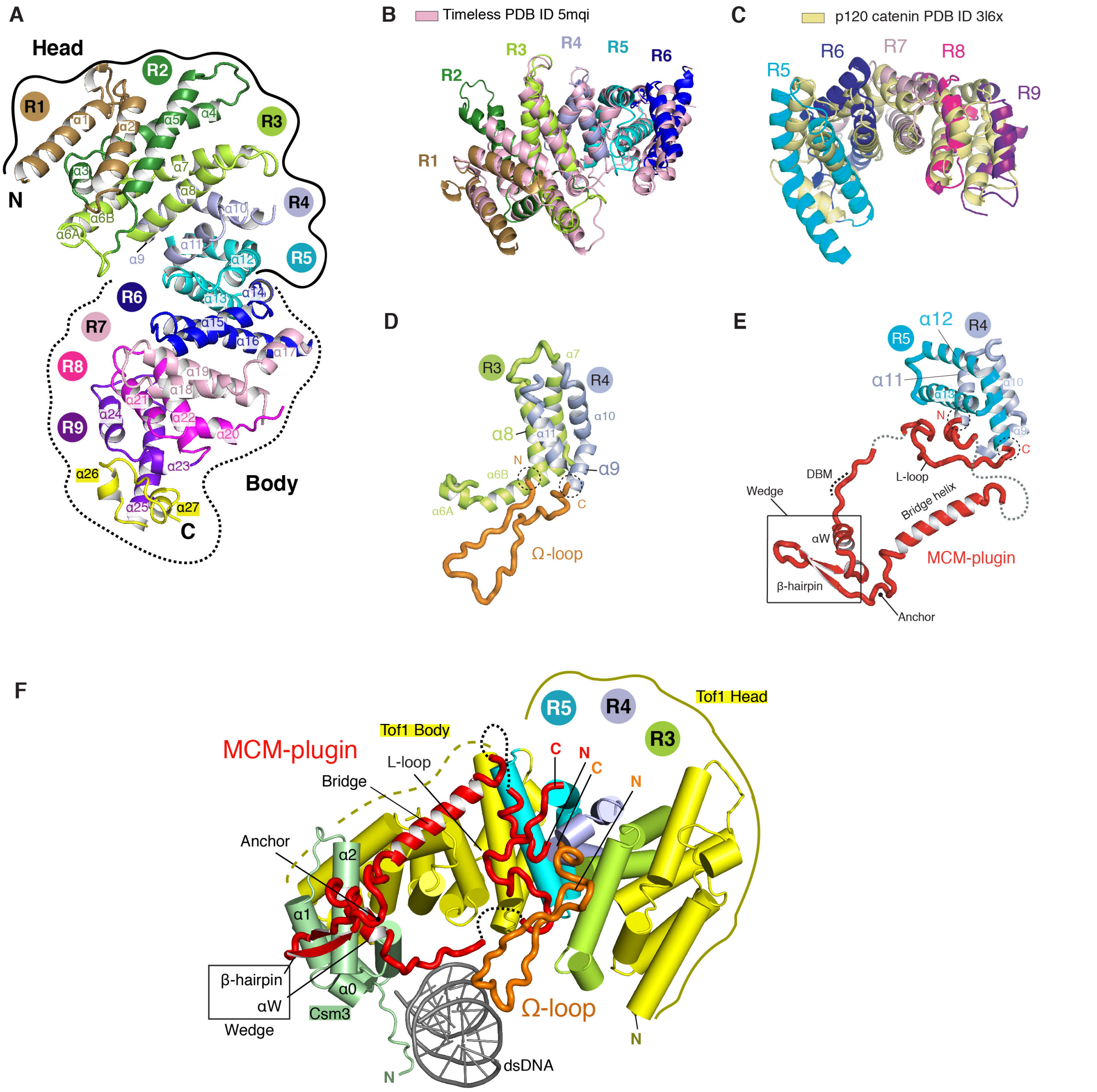

**Figure S9. Supporting figure 1 for Csm3/Tof1 structure**
(A) Assignment of Tof1 helical repeats. The model is coloured by helical repeat (R1-9) and the α-helices are numbered. The positions of the Head (repeats 1-5) and Body (repeats 6-9) are highlighted by solid and dashed black lines respectively.
(B) Superposition of Tof1 helical repeats 1-6 with the crystal structure of the human Timeless N-terminus (Holzer et al., 2017) (RMSD = 2 Å).
(C) Superposition of Tof1 helical repeats 5-9 with repeats 5-9 of p120 Catenin (Ishiyama et al., 2010) (RMSD = 6 Å). p120 Catenin was returned as the second highest scoring protein after Timeless in a Dali search (Holm and Sander, 1995) against Tof1 (13-781).
(D) Location of the Tof1 Ω-loop between helical repeats 3 and 4.
(E) Location of the MCM-plugin between helical repeats 4 and 5. The N- and C-termini of the Ω-loop and MCM-plugin are highlighted by dashed black circles.
(F) Bottom view of Csm3/Tof1. Tof1 repeats 3, 4 and 5 are highlighted to emphasize the positions of the Ω-loop and MCM-plugin.
DBM, DNA binding motif.

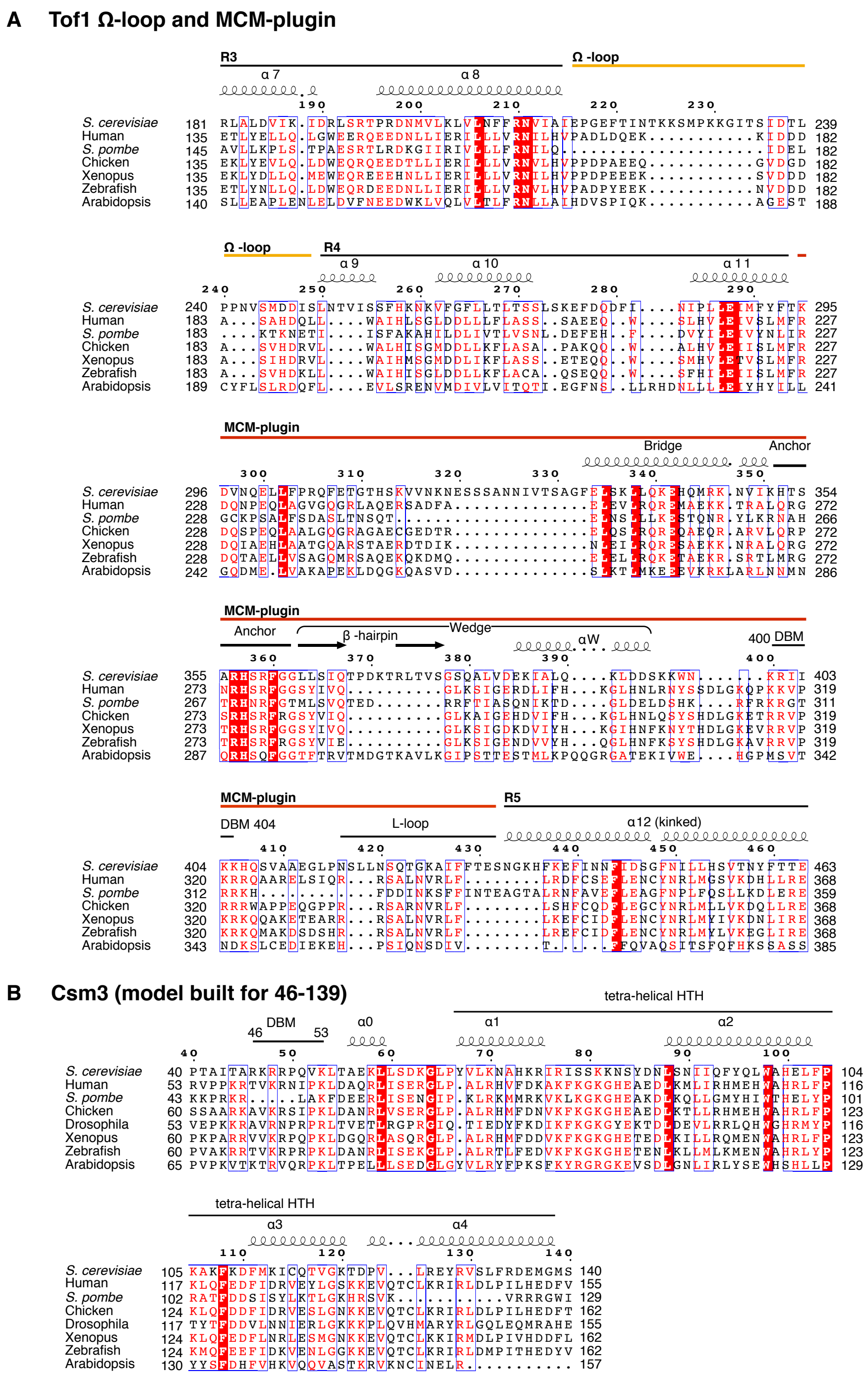

**Figure S10. Multiple sequence alignment of Csm3 and Tof1**

(A) The sequences of Tof1 (*S. cerevisiae*, 1238 residues, SGD YNL273W) and its orthologues: human Timeless (*H. sapiens*, 1208 residues, UniProtKB Q9UNS1), Swi1 (*S. pombe*, 971 residues, UniProtKB Q9UUM2), chicken Timeless (*G. gallus*, 1217 residues, NCBI ref XP_015155764.1), Xenopus Timeless (*X. tropicalis,* 1204 residues, UniProtKB F6Z6K7), zebrafish Timeless (*D. rerio*, 1278 residues, NCBI ref NP_001265529.1) and Arabidopsis Timeless (*A. thaliana*, 1141 residues, NCBI ref NP_200103.1).

(B) The sequence of Csm3 (*S. cerevisiae*, 317 residues, SGD YMR048W) and its orthologues: human Tipin (*H. sapiens*, 301 residues, UniProtKB Q9BVW5), Swi3 (*S. pombe*, 181 residues, UniProtKB O14350), chicken Tipin (*G. gallus*, 283 residues, UniProtKB Q5F416), Xenopus Tipin (*X. laevis,* 368 residues, UniProtKB Q0IHI4), zebrafish Tipin (*D. rerio*, 294 residues, UniProtKB G1K2L6) and Arabidopsis Tipin (*A. lyrata*, 287 residues, UniProtKB D7KZL5).

DBM, DNA binding motif; HTH, helix-turn-helix motif.

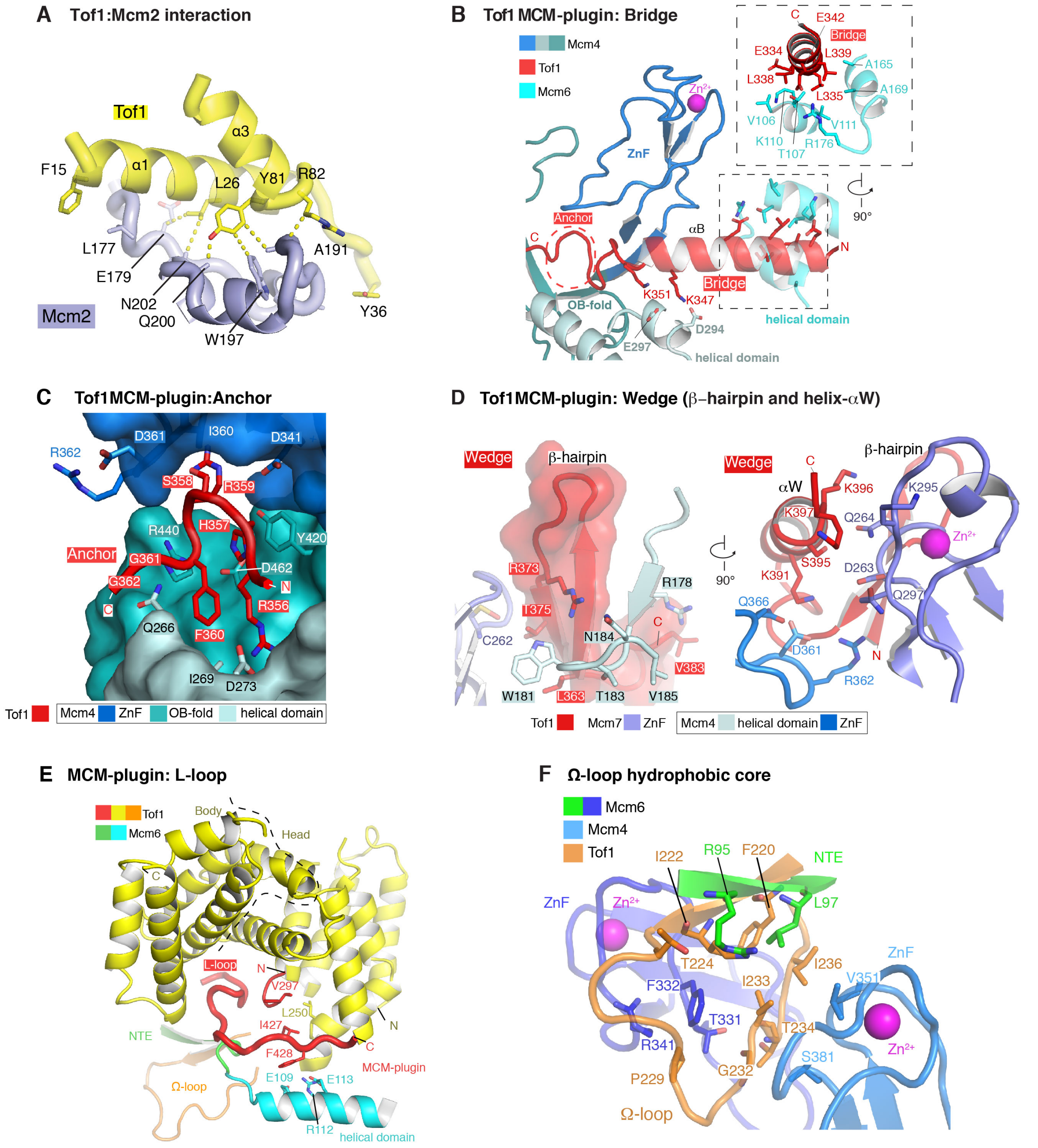

**Figure S11. Supporting figure 2 for Csm3/Tof1 structure**
(A) Interface between Tof1 (α1 and α3) and the Mcm2 NTE.
(B) Detailed view of interactions between the Bridge and Mcm6 and 4. The position of the Anchor is highlighted with a dashed red line and the inset shows details of the Bridge interface with the Mcm6 helical domain.
(C) Details of Anchor binding sites on Mcm4. The Mcm4 ZnF, OB-fold and helical domain are denoted by different shades of blue.
(D) Details of Wedge binding to Mcm4 and 7.
(E) Interactions of the L-loop with the Mcm6 helical domain and NTE. The dashed black line denotes the boundary of the Tof1 Head and Body.
(F) Details of the interactions between the hydrophobic core of the Ω-loop and Mcm6 and 4.
DBM, DNA binding motif.

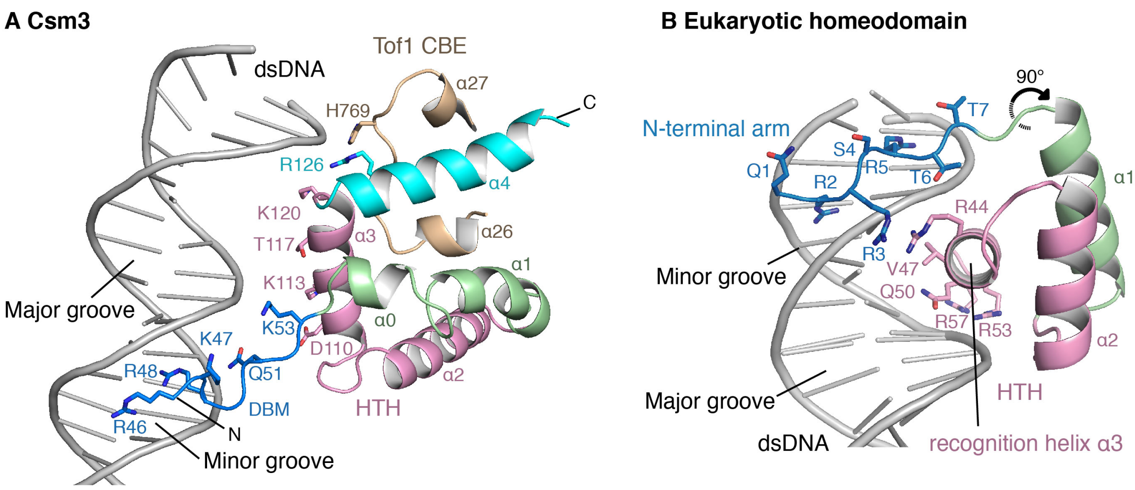

**Figure S12. Supporting figure 3 for Csm3/Tof1 structure**

(A) Overview of the Csm3 structure. Amino acid side chains for residues in the DBM and close to dsDNA are shown.

(B) Binding of a typical homeodomain transcription factor to dsDNA (Drosophila paired protein, PDB ID: 1FJL) (Wilson et al., 1995) emphasizing the minor groove contacts made by the N-terminal arm.

CBE, Csm3-binding element; DBM, DNA binding motif; HTH, helix-turn-helix motif.

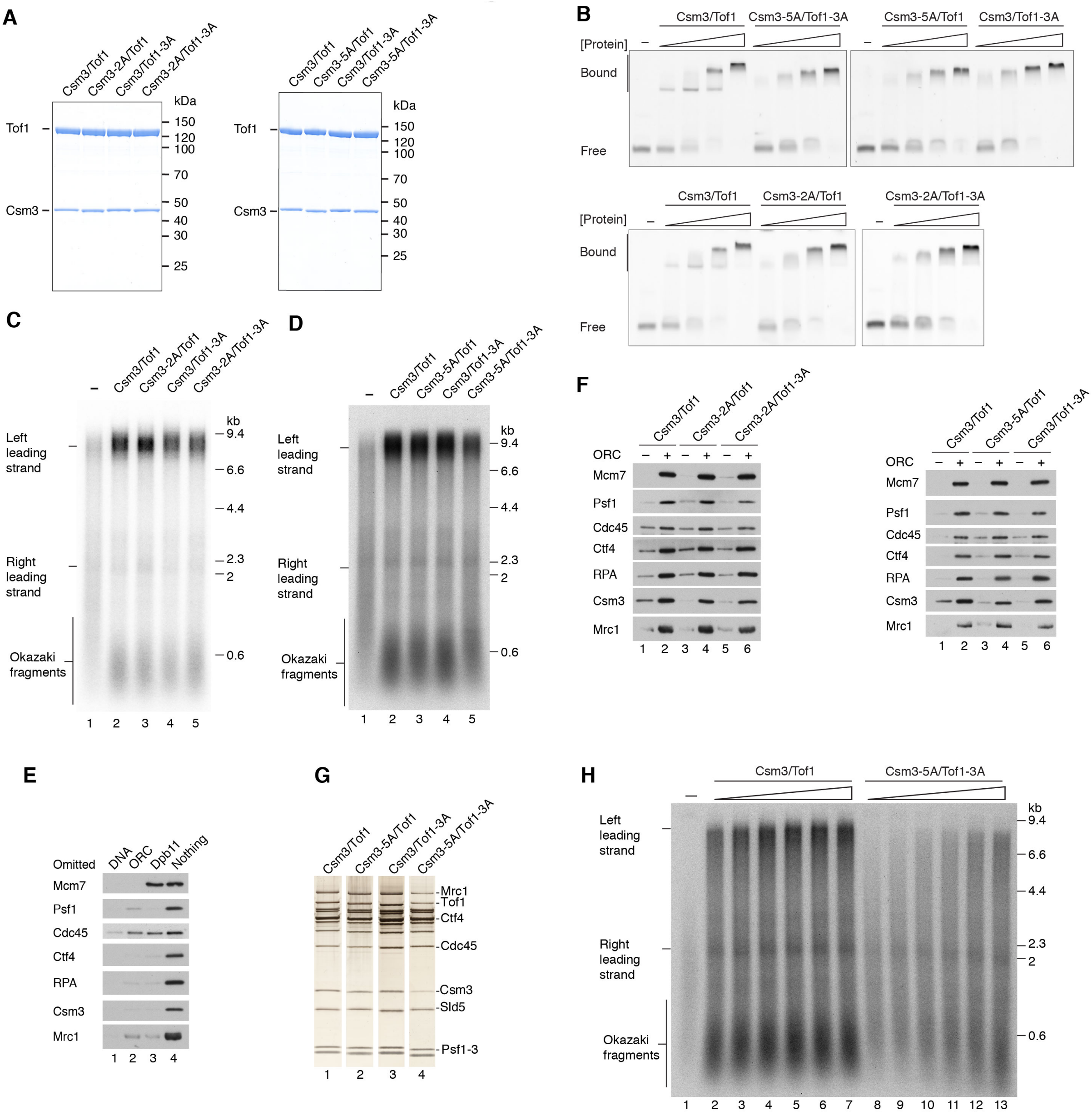

**Figure S13. Supporting figure 1 for Csm3/Tof1 functional analysis**

(A) Coomassie strained SDS-PAGE of Csm3/Tof1 DBM mutants.

(B) Electrophoretic mobility shift assays (EMSA) with the fork DNA substrate in Figure S1A and increasing concentrations (40, 80, 160, 320 nM) of the indicated Csm3/Tof1 proteins.

(C, D) Denaturing gel analysis of origin-dependent replication reactions performed for 15 (C) and 20 (D) min.

(E, F) Protein association assays performed as in Figure 6D.

(G) Peak fractions from native glycerol gradients performed as in Figure S1B with the indicated Csm3/Tof1 proteins.

(H) Denaturing gel analysis of an origin-dependent replication reaction performed for 7 min with increasing concentrations (2.5, 5, 10, 20, 40, 80 nM) of the indicated Csm3/Tof1 proteins.

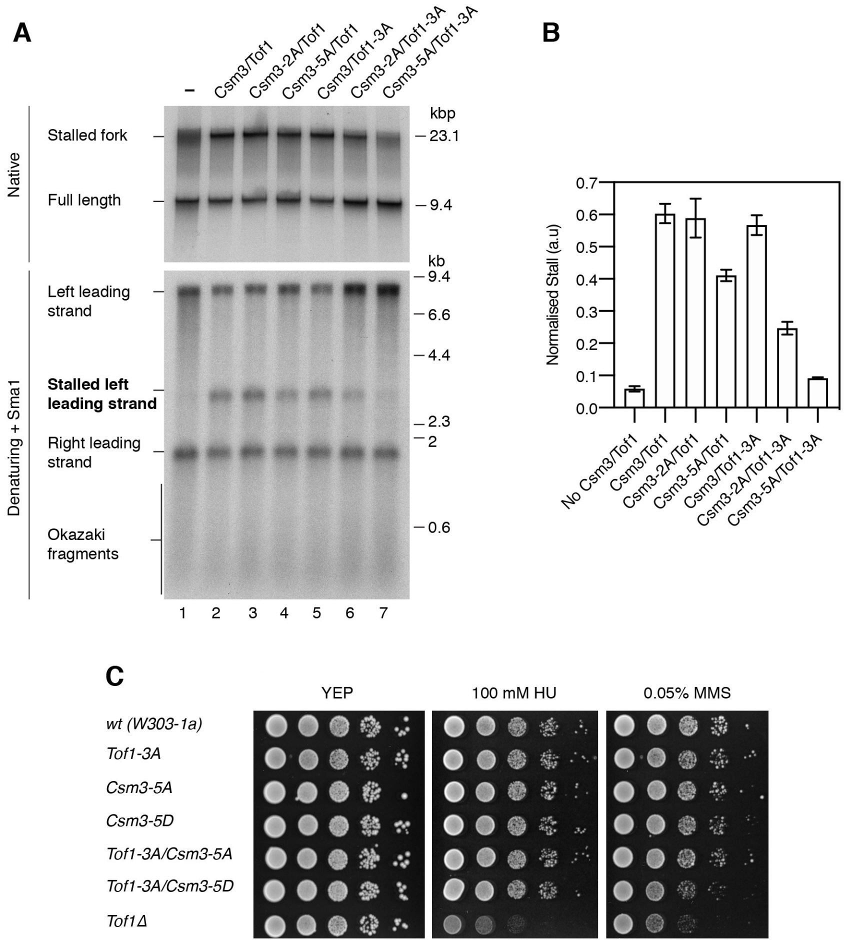

**Figure S14. Supporting figure 2 for Csm3/Tof1 functional analysis**
(A) Experiment performed and analyzed as in Figure 7B but with 100 mM potassium glutamate for 30 min. Pol δ was omitted as it catalyzes extensive strand-displacement synthesis in lower ionic strength buffers (Devbhandari et al., 2017).
(B) Quantitation of experiments performed as in (I). Error bars represent the SEM from a minimum of 3 experiments.
(C) Spot dilution assay performed for 3 days at 30ºC.
SEM, standard error of the mean.

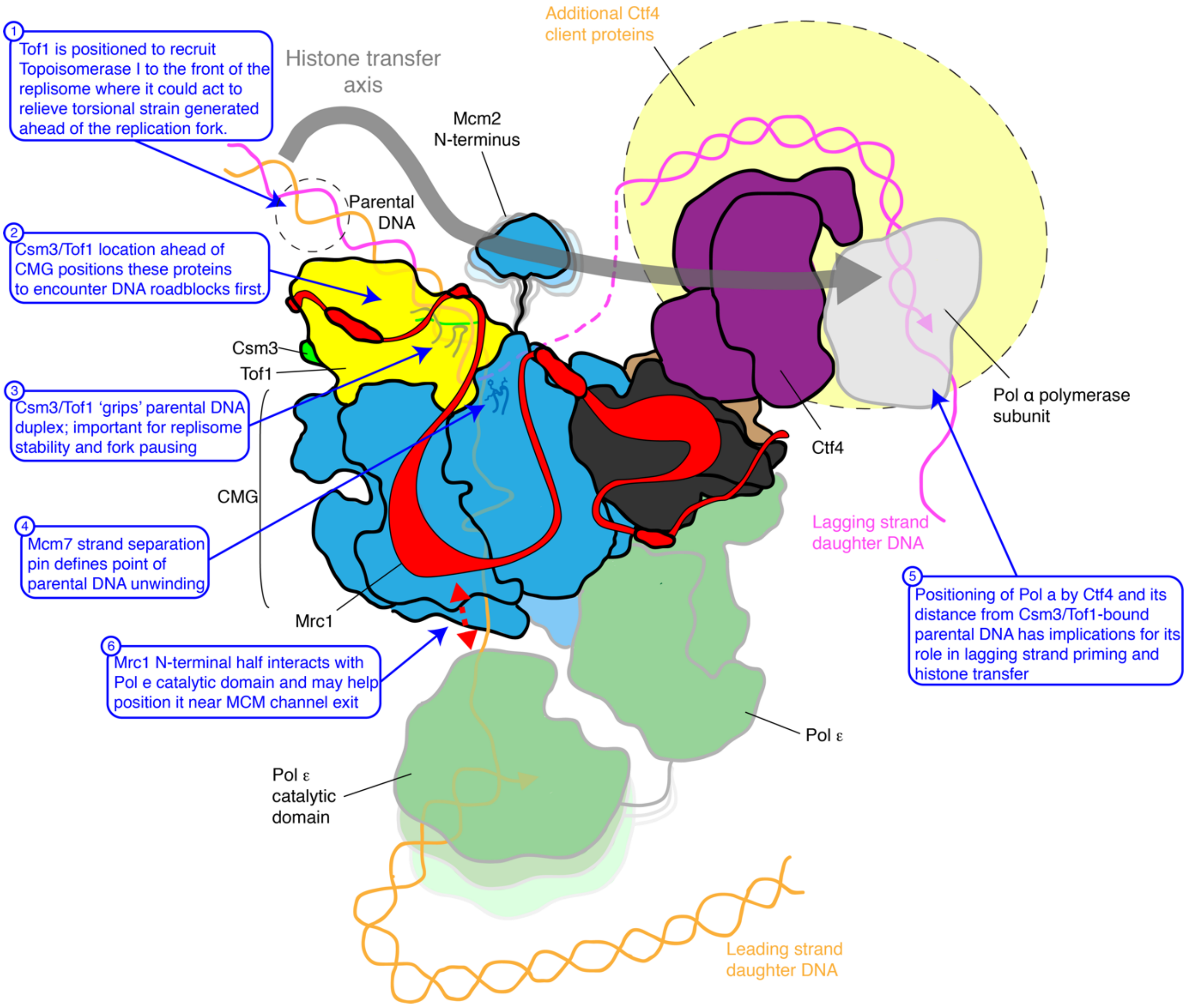

**Figure S15. Schematic representing the current view of the structure of the eukaryotic replisome**
The key highlights from the current work are indicated by blue boxes 1-4. The positions of the non-catalytic regions of DNA Pol ε and the polymerase subunit of DNA Pol α were derived from the structures of CMG-Pol ε (PDB-ID: 6HV9) (Goswami et al., 2018) and Ctf4-Pol α (Yuan et al., 2019), respectively. The N-terminus of Mcm2 - reportedly involved in a histone transfer axis with Ctf4 and Pol α (Gan et al., 2018) - was not resolved in the current structure and has been represented as flexibly tethered. The catalytic domain of Pol ε which was not resolved in prior structures is represented as flexibly tethered.

**Cryo-EM data collection, refinement and validation statistics**

| | Reconstitution: Conformation 1 [CMG-Csm3-Tof1-Ctf4-DNA] (EMD-10227) (PDB 6SKL) | Reconstitution: Conformation 2 [MCM C-tier] (EMD-10230) (PDB 6SKO) |
|---|---|---|
| **Data collection and processing** | | |
| Microscope / Detector | Titan Krios (Thermo Fisher) / K2 Summit (Gatan) | |
| Datasets | 2 | |
| Micrographs (used in processing) | 6,682 (20-frames per micrograph) | |
| Voltage (kV) | 300 | |
| GIF energy filter slit width (eV) | 20 | |
| Electron exposure ($e^{-}/Å^{2}$) | 37 | |
| Defocus range (μm) | -1.4 to -2.6 | |
| Pixel size (Å) | 1.049 | |
| Initial particle number | 632,077 | |
| Final particle number | Csm3/Tof1 (MBRs): 282,761<br>CMG + Ctf4 (MBRs): 198,120<br>Whole complex: 34,647 | 181,957 |
| Map resolution (Å)<br>0.143 FSC threshold | Csm3/Tof1 (MBRs): 3.2, 3.3 Å<br>CMG + Ctf4 (MBRs): 3.1 – 3.4 Å<br>Whole complex: 3.7 Å | C-tier (MBR): 3.4 Å<br>Mcm3467 (MBR): 3.3 Å<br>Mcm25 (MBR): 3.2 Å |
| **Refinement** | | |
| Model resolution (Å)<br>0.5 FSC threshold | 3.9 | 4.1 |
| Map sharpening *B* factor ($Å^{2}$) | -5 | -20 |
| Model composition | | |
| Non-hydrogen atoms | 58,210 | 15,491 |
| Protein residues | 7,359 | 2,012 |
| Ligands | 3 AMP-PNP, 3 $Mg^{2+}$, 3 $Zn^{2+}$ | 5 AMP-PNP, 5 $Mg^{2+}$ |
| R.m.s. deviations | | |
| Bond lengths (Å) | 0.009 | 0.004 |
| Bond angles (°) | 0.869 | 0.814 |
| Validation# | | |
| MolProbity score | 2.36 | 2.19 |
| Clashscore | 18.5 | 13.05 |
| Poor rotamers (%) | 0.79 | 0.13 |
| Ramachandran plot | | |
| Favored (%) | 88.15 | 89.46 |
| Allowed (%) | 11.24 | 9.98 |
| Outliers (%) | 0.61 | 0.56 |

#MolProbity validation server http://molprobity.biochem.duke.edu/

**Table S1. Cryo-EM statistics.** Maps mentioned are those useful for model building. For conformation 1, MBR maps derived from the same particles grouped together; for details about their derivation and individual resolutions quoted at the 0.143 FSC correlation coefficient, refer to Figure S2. MBR, multi-body refinement.

**Conformation 1:**

| Subunit | chain ID | Region | Residues | Residues built | model template (PDB ID) | Model building |
|---|---|---|---|---|---|---|
| Tof1 | X | repeats 1-6 | 1-550 | 13-213; 250-275; 278-294; 433-550 | 5MQI | Homology model docked and rebuilt |
| | | Ω-loop | 214-249 | 214-249 | - | *de novo* |
| | | MCM-plugin | 295-432 | 295-303; 329-405; 411-432 | - | *de novo* |
| | | repeats 7-8 | 551-704 | 551-607; 655-704 | - | *de novo* |
| | | CBE | 705-781 | 705-781 | - | *de novo* |
| Csm3 | Y | N-terminus (DBM, α0) | 1-63 | 46-63 | - | *de novo* |
| | | tetrahelical-HTH (α1- α4) | 64-139 | 64-139 | - | *de novo* |
| CMG | | | | | | |
| Mcm2 | 2 | N-tier | 173-471 | 173-471 | 5U8S | Docked and rebuilt |
| | | C-tier | 472-868 | 472-710; 738-868 | | Docked and rebuilt |
| Mcm3 | 3 | N-tier | 18-336 | 18-56; 90-333 | | Docked and rebuilt |
| | | C-tier | 337-740 | 337-583; 648-689; 696-740 | | Docked and rebuilt |
| Mcm4 | 4 | N-tier | 174-505 | 174-469 | | Docked and rebuilt |
| | | C-tier | 505-852 | 505-550; 557-595; 598-733; 739-780; 790-852 | | Docked and rebuilt |
| | | WH | 853-933 | 853-929 | | Docked |
| Mcm5 | 5 | N-tier | 20-343 | 20-107; 131-198; 226-305; 319-343 | | Docked and rebuilt |
| | | C-tier | 344-694 | 344-694 | | Docked and rebuilt |
| Mcm6 | 6 | N-tier | 92-496 | 92-201; 252-419; 434-463 | | Docked and rebuilt |
| | | C-tier | 497-840 | 497-737; 744-840 | | Docked and rebuilt |
| Mcm7 | 7 | N-tier | 1-393 | 2-34; 60-157; 188-210; 218-386 | | Docked and rebuilt |
| | | C-tier | 394-730 | 394-443; 450-675; 679-730 | | Docked and rebuilt |
| Psf1 | A | - | 1-208 | 1-110, 119-208 | | Docked and rebuilt |
| Psf2 | B | - | 1-213 | 1-38, 47-201 | | Docked and rebuilt |
| Psf3 | C | - | 1-194 | 1-58, 68-144, 158-194 | | Docked and rebuilt |
| Sld5 | D | - | 1-294 | 1-15, 55-107, 120-294 | | Docked and rebuilt |
| | | CIP-box | 2-13 | 2-13 | | Docked and rebuilt (using PDB ID: 4C95) |

| | | | | | | |
|---|---|---|---|---|---|---|
| Cdc45 | E | - | 1-650 | 1-108, 111-167, 224-436, 460-593, 598-650 | | Docked and rebuilt |
| **Ctf4 trimer** | | | | | | |
| Ctf4 monomer (CMG interface) | F | C-terminal regions | 473-927 | 473-664, 670-791, 814-923 | 4C8H | Docked and rebuilt |
| Ctf4 monomer (GINS-facing) | G | | | 474-663, 669-790, 814-924 | | Docked and rebuilt |
| Ctf4 monomer (Cdc45-facing) | H | | | 473-665, 670-790, 814-925 | | Docked and rebuilt |
| **Fork DNA** | | | | | | |
| Leading-strand template | I | - | - | 37 nt | - | *de novo* |
| Lagging-strand template | J | - | - | 22 nt | - | *de novo* |

**Table S2. Summary of cryo-EM model building.** See Methods for further details. For CMG and Ctf4, boundaries for the regions listed were approximated from the structure.

| Score | m/z | Charge | Deviation in ppm | Peptide 1 | Protein 1 | From | To | Peptide 2 | Protein 2 | From | To | best linkage position peptide 1 | best linkage position peptide 2 |
|---|---|---|---|---|---|---|---|---|---|---|---|---|---|
| 131 | 616.0022312 | 3 | -0.448750556 | [ADESLPKR] | >MRC1 | 619 | 626 | [KSHHVK] | >MRC1 | 594 | 599 | K7 | K1 |
| 130 | 778.0758058 | 3 | 0.451348166 | [YVKFNBLK] | >MCM2 | 336 | 343 | [VNGEKTVYR] | >MCM2 | 375 | 383 | K3 | K5 |
| 129 | 778.0752824 | 3 | -0.221939582 | [YVKFNBLK] | >MCM2 | 336 | 343 | [VNGEKTVYR] | >MCM2 | 375 | 383 | K3 | K5 |
| 129 | 682.7434816 | 3 | 0.739697119 | [MLSLLFKK] | >CDC45 | 44 | 51 | [KVELLNR] | >CDC45 | 488 | 494 | K7 | K1 |
| 128 | 760.3948426 | 3 | -0.013860309 | [ASITYMGKTR] | >MRC1 | 1042 | 1051 | [SYKTVGSSK] | >MRC1 | 1033 | 1041 | K8 | K3 |
| 128 | 682.7428589 | 3 | -0.173246784 | [MLSLLFKK] | >CDC45 | 44 | 51 | [KVELLNR] | >CDC45 | 488 | 494 | K7 | K1 |
| 127 | 759.4246792 | 3 | -0.608840956 | [NILKEVSNLR] | >PSF1 | 39 | 48 | [KNTEYLK] | >PSF1 | 49 | 55 | K4 | K1 |
| 127 | 759.423286 | 3 | -2.444967393 | [NILKEVSNLR] | >PSF1 | 39 | 48 | [KNTEYLK] | >PSF1 | 49 | 55 | K4 | K1 |
| 127 | 616.0027955 | 3 | 0.468192247 | [ADESLPKR] | >MRC1 | 619 | 626 | [KSHHVK] | >MRC1 | 594 | 599 | K7 | K1 |
| 126 | 928.1456164 | 3 | -3.172552208 | [SFKNFILEFR] | >MCM5 | 30 | 39 | [IQYEKTHPDR] | >PSF2 | 102 | 111 | K3 | K5 |
| 126 | 759.426266 | 3 | 1.482512414 | [NILKEVSNLR] | >PSF1 | 39 | 48 | [KNTEYLK] | >PSF1 | 49 | 55 | S7 | K1 |
| 125 | 759.423575 | 3 | -2.064161344 | [NILKEVSNLR] | >PSF1 | 39 | 48 | [KNTEYLK] | >PSF1 | 49 | 55 | K4 | K1 |
| 123 | 778.0754983 | 3 | 0.055741075 | [YVKFNBLK] | >MCM2 | 336 | 343 | [VNGEKTVYR] | >MCM2 | 375 | 383 | K3 | K5 |
| 123 | 750.0462687 | 3 | -0.081641395 | [SYLVEKYK] | >MCM6 | 771 | 778 | [KDDAQGFSR] | >MCM6 | 782 | 790 | K6 | K1 |
| 123 | 1048.538204 | 3 | 0.200092247 | [EDESPTTKIFTIIK] | >MCM7 | 736 | 749 | [ESLYQETNKSK] | >MCM7 | 725 | 735 | T7 | K9 |
| 122 | 759.4240578 | 3 | -1.427804116 | [NILKEVSNLR] | >PSF1 | 39 | 48 | [KNTEYLK] | >PSF1 | 49 | 55 | K4 | K1 |
| 121 | 1048.537788 | 3 | -0.196729938 | [EDESPTTKIFTIIK] | >MCM7 | 736 | 749 | [ESLYQETNKSK] | >MCM7 | 725 | 735 | K8 | T7 |
| 121 | 734.5976753 | 5 | -1.82647435 | [VHKTHSFPISLANTGK] | >CTF4 | 458 | 473 | [VHKTHSFPISLANTGK] | >CTF4 | 458 | 473 | K3 | K3 |
| 121 | 950.8394048 | 3 | 1.42585679 | [SITTSTSPEQTER] | >MCM6 | 249 | 261 | [FLPFLQKGLR] | >MCM6 | 177 | 186 | S1 | K7 |
| 121 | 778.0763207 | 3 | 1.113666904 | [YVKFNBLK] | >MCM2 | 336 | 343 | [VNGEKTVYR] | >MCM2 | 375 | 383 | K3 | K5 |
| 121 | 760.0672274 | 3 | 0.499612163 | [NILKEVSNLR] | >PSF1 | 39 | 48 | {MYGDLGNK] | >PSF1 | 0 | 8 | S7 | {0 |
| 120 | 989.5287485 | 3 | -1.271258675 | [SKEDESPTTKIFTIIK] | >MCM7 | 734 | 749 | [KMLQETGK] | >MCM7 | 750 | 757 | T8 | K1 |
| 120 | 1037.556624 | 3 | 1.847086679 | [DNMVLKLVLNFFR] | >TOF1 | 198 | 210 | [WFKLVDDQQK] | >TOF1 | 61 | 70 | K6 | K3 |
| 120 | 751.0535147 | 3 | -0.048441994 | [LEEMEKEIYK] | >PSF3 | 169 | 178 | [LKBAPR] | >MCM5 | 608 | 613 | K6 | K2 |
| 120 | 751.0530282 | 3 | -0.696752242 | [LEEMEKEIYK] | >PSF3 | 169 | 178 | [LKBAPR] | >MCM5 | 608 | 613 | K6 | K2 |
| 120 | 989.5311194 | 3 | 1.126378303 | [SKEDESPTTKIFTIIK] | >MCM7 | 734 | 749 | [KMLQETGK] | >MCM7 | 750 | 757 | T8 | K1 |
| 119 | 805.7938345 | 3 | -0.425413621 | [LLEENKAILNK] | >CTF4 | 786 | 796 | [MPVFVKSK] | >CTF4 | 778 | 785 | K6 | K6 |
| 119 | 760.0651294 | 3 | -2.263160992 | [NILKEVSNLR] | >PSF1 | 39 | 48 | {MYGDLGNK] | >PSF1 | 0 | 8 | S7 | {0 |
| 119 | 1037.556185 | 3 | 1.424112099 | [DNMVLKLVLNFFR] | >TOF1 | 198 | 210 | [WFKLVDDQQK] | >TOF1 | 61 | 70 | K6 | K3 |
| 119 | 750.0460795 | 3 | -0.334056353 | [SYLVEKYK] | >MCM6 | 771 | 778 | [KDDAQGFSR] | >MCM6 | 782 | 790 | K6 | K1 |
| 119 | 760.0670188 | 3 | 0.224933296 | [NILKEVSNLR] | >PSF1 | 39 | 48 | {MYGDLGNK] | >PSF1 | 0 | 8 | S7 | {0 |
| 118 | 914.7819484 | 3 | -0.107900895 | [VSKESLYQETNK] | >MCM7 | 722 | 733 | [SKEDESPTTK] | >MCM7 | 734 | 743 | K3 | S1 |
| 118 | 760.3951961 | 3 | 0.451321987 | [ASITYMGKTR] | >MRC1 | 1042 | 1051 | [SYKTVGSSK] | >MRC1 | 1033 | 1041 | K8 | K3 |
| 118 | 1223.659187 | 3 | -0.807726868 | [VHKTHSFPISLANTGK] | >CTF4 | 458 | 473 | [VHKTHSFPISLANTGK] | >CTF4 | 458 | 473 | K3 | K3 |
| 118 | 1278.983188 | 3 | 2.035815464 | [KDANLDYYNFNPMGIK] | >CTF4 | 670 | 685 | [VHKTHSFPISLANTGK] | >CTF4 | 458 | 473 | K1 | K3 |
| 117 | 889.4765791 | 3 | -1.121760788 | [NTLSYENIVKTVR] | >MCM7 | 758 | 770 | [KMLQETGK] | >MCM7 | 750 | 757 | K10 | T6 |
| 117 | 734.5985257 | 5 | -0.667553652 | [VHKTHSFPISLANTGK] | >CTF4 | 458 | 473 | [VHKTHSFPISLANTGK] | >CTF4 | 458 | 473 | K3 | K3 |
| 117 | 841.1777981 | 3 | -1.585445222 | [TYVDVVHVKK] | >MCM4 | 459 | 468 | [LINLKGLVLR] | >MCM4 | 325 | 334 | K9 | K5 |
| 116 | 644.3695068 | 3 | 1.011421522 | [ALTAAVKISER] | >CTF4 | 894 | 904 | [TKSNK] | >MRC1 | 1081 | 1085 | K7 | T1 |
| 116 | 989.5273516 | 3 | -2.683913309 | [SKEDESPTTKIFTIIK] | >MCM7 | 734 | 749 | [KMLQETGK] | >MCM7 | 750 | 757 | T8 | K1 |
| 116 | 989.5269047 | 3 | -3.135837697 | [SKEDESPTTKIFTIIK] | >MCM7 | 734 | 749 | [KMLQETGK] | >MCM7 | 750 | 757 | T8 | K1 |
| 116 | 769.1094971 | 3 | 0.592786942 | [SQILKYVEK] | >MCM2 | 550 | 558 | [STVEGIKLR] | >MCM6 | 558 | 566 | Y6 | K7 |
| 116 | 708.1370719 | 4 | 0.249621274 | [ALYALEKHETIQLR] | >MCM5 | 751 | 764 | [FEQELKR] | >MCM5 | 714 | 720 | T10 | K6 |
| 116 | 742.3984197 | 4 | -1.218443437 | [SKEDESPTTKIFTIIK] | >MCM7 | 734 | 749 | [KMLQETGK] | >MCM7 | 750 | 757 | T9 | K1 |
| 115 | 750.0463305 | 3 | 8.23E-04 | [SYLVEKYK] | >MCM6 | 771 | 778 | [KDDAQGFSR] | >MCM6 | 782 | 790 | K6 | K1 |
| 115 | 769.1085426 | 3 | -0.649248605 | [SQILKYVEK] | >MCM2 | 550 | 558 | [STVEGIKLR] | >MCM6 | 558 | 566 | Y6 | K7 |
| 115 | 785.0797517 | 3 | -2.210872593 | [SKIDNSADIK] | >TOF1 | 490 | 499 | [KVFSFFHR] | >TOF1 | 692 | 699 | S1 | K1 |
| 115 | 917.8203589 | 3 | -0.708281983 | [EDESPTTKIFTIIK] | >MCM7 | 736 | 749 | [KMLQETGK] | >MCM7 | 750 | 757 | T7 | K1 |
| 115 | 682.7428968 | 3 | -0.11771984 | [MLSLLFKK] | >CDC45 | 44 | 51 | [KVELLNR] | >CDC45 | 488 | 494 | K7 | K1 |
| 115 | 595.0061035 | 3 | -1.943153504 | [NINKLSK] | >MCM3 | 349 | 355 | [SPQKSPK] | >MCM3 | 761 | 767 | K4 | K4 |
| 115 | 940.8386425 | 3 | 0.393107413 | [LAKNNPDHEEFR] | >TOF1 | 105 | 116 | [LALDVIKIDR] | >TOF1 | 182 | 191 | K3 | K7 |
| 115 | 995.1963888 | 3 | 0.085407618 | [IBQTVGKTDPVLR] | >CSM3 | 114 | 126 | [IGSKYSLQFMK] | >TOF1 | 579 | 589 | K7 | S3 |
| 114 | 708.1364304 | 4 | -0.657375612 | [ALYALEKHETIQLR] | >MCM5 | 751 | 764 | [FEQELKR] | >MCM5 | 714 | 720 | T10 | K6 |
| 114 | 897.4655052 | 4 | -0.542223767 | [LFASABSDQNVEKALSLAHELK] | >CTF4 | 869 | 890 | [SKLLEENK] | >CTF4 | 784 | 791 | K13 | S1 |
| 114 | 548.6787109 | 3 | 0.54272402 | [ILKSYK] | >MRC1 | 1030 | 1035 | [LIAPKR] | >MRC1 | 1053 | 1058 | K3 | K5 |
| 114 | 742.3982599 | 4 | -1.433864114 | [SKEDESPTTKIFTIIK] | >MCM7 | 734 | 749 | [KMLQETGK] | >MCM7 | 750 | 757 | T8 | K1 |
| 114 | 1145.95216 | 3 | -1.828480511 | [GISKTTEGLNSGVTGLR] | >MCM6 | 425 | 441 | [SLYKTYVDVVHVK] | >MCM4 | 455 | 467 | K4 | S1 |
| 114 | 989.530055 | 3 | 0.049980186 | [SKEDESPTTKIFTIIK] | >MCM7 | 734 | 749 | [KMLQETGK] | >MCM7 | 750 | 757 | K10 | T6 |
| 113 | 940.8383118 | 3 | 0.041379182 | [LAKNNPDHEEFR] | >TOF1 | 105 | 116 | [LALDVIKIDR] | >TOF1 | 182 | 191 | K3 | K7 |
| 113 | 897.4668595 | 4 | 0.968070661 | [LFASABSDQNVEKALSLAHELK] | >CTF4 | 869 | 890 | [SKLLEENK] | >CTF4 | 784 | 791 | K13 | S1 |
| 113 | 708.1380067 | 4 | 1.571105339 | [ALYALEKHETIQLR] | >MCM5 | 751 | 764 | [FEQELKR] | >MCM5 | 714 | 720 | T10 | K6 |
| 113 | 989.5306391 | 3 | 0.640647375 | [SKEDESPTTKIFTIIK] | >MCM7 | 734 | 749 | [KMLQETGK] | >MCM7 | 750 | 757 | T8 | K1 |
| 113 | 897.4667028 | 4 | 0.793337882 | [LFASABSDQNVEKALSLAHELK] | >CTF4 | 869 | 890 | [SKLLEENK] | >CTF4 | 784 | 791 | K13 | S1 |
| 113 | 989.5299635 | 3 | -0.042628279 | [SKEDESPTTKIFTIIK] | >MCM7 | 734 | 749 | [KMLQETGK] | >MCM7 | 750 | 757 | T8 | K1 |
| 112 | 989.5295081 | 3 | -0.503162913 | [SKEDESPTTKIFTIIK] | >MCM7 | 734 | 749 | [KMLQETGK] | >MCM7 | 750 | 757 | K10 | T6 |
| 112 | 760.066805 | 3 | -0.056562549 | [NILKEVSNLR] | >PSF1 | 48 | | {MYGDLGNK] | >PSF1 | 0 | 8 | K4 | {0 |
| 112 | 1182.954493 | 3 | -3.247699646 | [THSFPISLANTGKFR] | >CTF4 | 461 | 475 | [THSFPISLANTGKFR] | >CTF4 | 461 | 475 | T11 | K13 |
| 112 | 1109.558948 | 3 | -0.370900287 | [YFLYFKPLSQNBAR] | >MCM7 | 200 | 213 | [YMAMLQKVANR] | >MCM7 | 63 | 73 | K6 | K7 |
| 112 | 1201.650429 | 3 | 1.053096482 | [SVYTSGKASSAAGLTAAVVR] | >MCM6 | 595 | 614 | [SQFLKYVVGFAPR] | >MCM6 | 582 | 594 | K7 | K5 |
| 112 | 989.530566 | 3 | 0.566737326 | [SKEDESPTTKIFTIIK] | >MCM7 | 734 | 749 | [KMLQETGK] | >MCM7 | 750 | 757 | K10 | T6 |
| 112 | 817.1154737 | 3 | 0.894109136 | [KFNISEGDITK] | >TOF1 | 622 | 632 | [TLVIEGKSR] | >TOF1 | 610 | 618 | K1 | K7 |
| 112 | 841.1804286 | 3 | 1.544155749 | [TYVDVVHVKK] | >MCM4 | 459 | 468 | [LINLKGLVLR] | >MCM4 | 325 | 334 | K9 | K5 |
| 111 | 780.1287702 | 3 | 0.571662737 | [ALTAAVKISER] | >CTF4 | 894 | 904 | [AELPSLVKK] | >CTF4 | 905 | 913 | K7 | S5 |
| 111 | 1037.556626 | 3 | 1.849484883 | [DNMVLKLVLNFFR] | >TOF1 | 198 | 210 | [WFKLVDDQQK] | >TOF1 | 61 | 70 | K6 | K3 |

| | | | | | | | | | | | | | |
|---|---|---|---|---|---|---|---|---|---|---|---|---|---|
| 111 | 682.743171 | 3 | 0.284310305 | [MLSLLFKK] | >CDC45 | 44 | 51 | [KVELLNR] | >CDC45 | 488 | 494 | K7 | K1 |
| 111 | 701.8911872 | 4 | 0.898727555 | [SKEDESPTTKIFTIIK] | >MCM7 | 734 | 749 | [KSPITAR] | >MCM3 | 694 | 700 | T8 | K1 |
| 111 | 1145.955344 | 3 | 0.951828104 | [GISKTTEGLNSGVTGLR] | >MCM6 | 425 | 441 | [SLYKTYVDVVHVK] | >MCM4 | 455 | 467 | K4 | S1 |
| 111 | 897.4672941 | 4 | 1.452679609 | [LFASABSDQNVEKALSLAHELK] | >CTF4 | 869 | 890 | [SKLLEENK] | >CTF4 | 784 | 791 | K13 | S1 |
| 110 | 981.1855227 | 3 | -2.439057461 | [SLYKTYVDVVHVK] | >MCM4 | 455 | 467 | [BSFADKQVIK] | >MCM4 | 389 | 398 | K4 | K6 |
| 110 | 1139.621453 | 3 | -1.118720523 | [ALYALEKHETIQLR] | >MCM5 | 751 | 764 | [NTLSYENIVKTVR] | >MCM7 | 758 | 770 | K7 | K10 |
| 110 | 780.128542 | 3 | 0.278903553 | [ALTAAVKISER] | >CTF4 | 894 | 904 | [AELPSLVKK] | >CTF4 | 905 | 913 | K7 | S5 |
| 110 | 778.0749512 | 3 | -0.648016176 | [YVKFNBLK] | >MCM2 | 336 | 343 | [VNGEKTVYR] | >MCM2 | 375 | 383 | K3 | K5 |
| 110 | 989.5304539 | 3 | 0.453347439 | [SKEDESPTTKIFTIIK] | >MCM7 | 734 | 749 | [KMLQETGK] | >MCM7 | 750 | 757 | T8 | K1 |
| 110 | 1201.651032 | 3 | 1.555252728 | [SVYTSGKASSAAGLTAAVVR] | >MCM6 | 595 | 614 | [SQFLKYVVGFAPR] | >MCM6 | 582 | 594 | K7 | K5 |
| 110 | 1067.529239 | 3 | 0.533448202 | [EMDSKFSFGQATPR] | >MCM7 | 674 | 687 | [TGKIDMNLVQTGK] | >MCM4 | 842 | 854 | K5 | T1 |
| 109 | 708.1367188 | 4 | -0.249677946 | [ALYALEKHETIQLR] | >MCM5 | 751 | 764 | [FEQELKR] | >MCM5 | 714 | 720 | T10 | K6 |
| 109 | 917.8197632 | 3 | -1.357824837 | [EDESPTTKIFTIIK] | >MCM7 | 736 | 749 | [KMLQETGK] | >MCM7 | 750 | 757 | T7 | K1 |
| 109 | 711.7266857 | 3 | -2.231650564 | [LSLWNENGKK] | >CDC45 | 348 | 357 | [TKIYPK] | >MCM2 | 772 | 777 | K9 | T1 |
| 109 | 1095.950866 | 3 | 0.884170175 | [IDMNLVQTGKSVIQR] | >MCM4 | 845 | 859 | [KGILLQLFGGTNK] | >MCM4 | 538 | 550 | K10 | K1 |
| 109 | 742.3995234 | 4 | 0.269856538 | [SKEDESPTTKIFTIIK] | >MCM7 | 734 | 749 | [KMLQETGK] | >MCM7 | 750 | 757 | T9 | K1 |
| 109 | 1201.650308 | 3 | 0.95229825 | [SVYTSGKASSAAGLTAAVVR] | >MCM6 | 595 | 614 | [SQFLKYVVGFAPR] | >MCM6 | 582 | 594 | K7 | K5 |
| 109 | 1182.959895 | 3 | 1.321620593 | [THSFPISLANTGKFR] | >CTF4 | 461 | 475 | [THSFPISLANTGKFR] | >CTF4 | 461 | 475 | T11 | K13 |
| 108 | 1395.752299 | 3 | 1.288025772 | [LYEILTNSIAPSIFGNEDIKK] | >MCM5 | 366 | 386 | [FVEKVSPIAVYTSGK] | >MCM5 | 428 | 442 | S12 | K4 |
| 108 | 1145.957591 | 3 | 2.913525344 | [GISKTTEGLNSGVTGLR] | >MCM6 | 425 | 441 | [SLYKTYVDVVHVK] | >MCM4 | 455 | 467 | K4 | S1 |
| 107 | 700.3287128 | 3 | 0.292033491 | [DYATDPKTGK] | >MCM4 | 835 | 844 | [KMGDDSR] | >MCM4 | 779 | 785 | K7 | K1 |
| 107 | 1040.916909 | 3 | -0.436361783 | [LTVSGSQALVDEKIALQK] | >TOF1 | 374 | 391 | [NVIKHTSAR] | >TOF1 | 348 | 356 | K13 | K4 |
| 107 | 759.4246552 | 3 | -0.640401525 | [NILKEVSNLR] | >PSF1 | 39 | 48 | [KNTEYLK] | >PSF1 | 49 | 55 | K4 | K1 |
| 107 | 1203.307369 | 3 | -1.567537393 | [VHKTHSFPISLANTGK] | >CTF4 | 458 | 473 | [THSFPISLANTGKFR] | >CTF4 | 461 | 475 | K3 | S7 |
| 107 | 913.2113434 | 4 | -0.822417841 | [SBIELTHSVLKVLEQYSDDK] | >TOF1 | 590 | 609 | [SKIDNSADIK] | >TOF1 | 490 | 499 | K11 | S1 |
| 106 | 928.1483991 | 3 | -0.172273836 | [SFKNFILEFR] | >MCM5 | 30 | 39 | [IQYEKTHPDR] | >PSF2 | 102 | 111 | K3 | K5 |
| 106 | 660.3783496 | 3 | -0.696723622 | [LLQKEHQMR] | >TOF1 | 338 | 346 | [KNVIK] | >TOF1 | 347 | 351 | K4 | K1 |
| 106 | 1095.238532 | 3 | 2.025516303 | [VLEQYSDDKTLVIEGK] | >TOF1 | 601 | 616 | [KFNISEGDITK] | >TOF1 | 622 | 632 | K9 | K1 |
| 105 | 1401.724075 | 3 | 1.921774206 | [YDDLKSPGDNIDFQTTILSR] | >MCM5 | 530 | 549 | [FSQLALDKALYALEK] | >MCM5 | 743 | 757 | Y1 | K8 |
| 105 | 901.4890953 | 4 | 0.447174373 | [SVYTSGKASSAAGLTAAVVR] | >MCM6 | 595 | 614 | [SQFLKYVVGFAPR] | >MCM6 | 582 | 594 | S5 | K5 |
| 105 | 1072.929 | 3 | -0.458335187 | [VHKTHSFPISLANTGK] | >CTF4 | 458 | 473 | [LLEENKAILNK] | >CTF4 | 786 | 796 | K3 | K6 |
| 105 | 1182.959257 | 3 | 0.782217929 | [THSFPISLANTGKFR] | >CTF4 | 461 | 475 | [THSFPISLANTGKFR] | >CTF4 | 461 | 475 | T11 | K13 |
| 104 | 682.7431912 | 3 | 0.313921066 | [MLSLLFKK] | >CDC45 | 44 | 51 | [KVELLNR] | >CDC45 | 488 | 494 | K7 | K1 |
| 104 | 759.4255935 | 3 | 0.596209867 | [NILKEVSNLR] | >PSF1 | 39 | 48 | [KNTEYLK] | >PSF1 | 49 | 55 | K4 | K1 |
| 104 | 1389.341471 | 3 | 0.802343276 | [NTEYLKEQQQLGMLDDK] | >PSF1 | 50 | 66 | [VAKBQYFVTLLBMER] | >PSF1 | 67 | 81 | K6 | K3 |
| 103 | 680.7363892 | 3 | 1.086857099 | [ATILNLKAR] | >MRC1 | 439 | 447 | [LSKQNQK] | >MRC1 | 448 | 454 | K7 | K3 |
| 103 | 967.5238286 | 3 | -2.045465797 | [FNPLLQAGAKLAK] | >MCM3 | 625 | 637 | [NKGNYNGTEIPK] | >MCM3 | 638 | 649 | K10 | K2 |
| 103 | 1182.955582 | 3 | -2.326513638 | [THSFPISLANTGKFR] | >CTF4 | 461 | 475 | [THSFPISLANTGKFR] | >CTF4 | 461 | 475 | T11 | K13 |
| 103 | 841.1794139 | 3 | 0.336887575 | [TYVDVVHVKK] | >MCM4 | 459 | 468 | [LINLKGLVLR] | >MCM4 | 325 | 334 | K9 | K5 |
| 103 | 1109.561115 | 3 | 1.582934861 | [YFLYFKPLSQNBAR] | >MCM7 | 200 | 213 | [YMAMLQKVANR] | >MCM7 | 63 | 73 | S9 | K7 |
| 103 | 734.5980956 | 5 | -1.25368995 | [VHKTHSFPISLANTGK] | >CTF4 | 458 | 473 | [VHKTHSFPISLANTGK] | >CTF4 | 458 | 473 | K3 | K3 |
| 103 | 981.1890168 | 3 | 1.124467438 | [SLYKTYVDVVHVK] | >MCM4 | 455 | 467 | [BSFADKQVIK] | >MCM4 | 389 | 398 | K4 | K6 |
| 103 | 1102.264415 | 3 | 1.204933937 | [KQLLINELESTER] | >MCM5 | 631 | 643 | [NTLSYENIVKTVR] | >MCM7 | 758 | 770 | S10 | K10 |
| 102 | 875.4336922 | 3 | -0.265691263 | [LFASABSDQNVEK] | >CTF4 | 869 | 881 | [SKLLEENK] | >CTF4 | 784 | 791 | S4 | K2 |
| 102 | 750.0457764 | 3 | -0.738598451 | [SYLVEKYK] | >MCM6 | 771 | 778 | [KDDAQGFSR] | >MCM6 | 782 | 790 | K6 | K1 |
| 102 | 553.0859973 | 4 | 0.387960622 | [LINLKGLVLR] | >MCM4 | 325 | 334 | [VGDGMKIR] | >MCM7 | 444 | 451 | K5 | K6 |
| 102 | 1109.559171 | 3 | -0.169713616 | [YFLYFKPLSQNBAR] | >MCM7 | 200 | 213 | [YMAMLQKVANR] | >MCM7 | 63 | 73 | K6 | K7 |
| 102 | 875.4340377 | 3 | 0.129216022 | [LFASABSDQNVEK] | >CTF4 | 869 | 881 | [SKLLEENK] | >CTF4 | 784 | 791 | S4 | K2 |
| 102 | 1095.952266 | 3 | 2.162045366 | [IDMNLVQTGKSVIQR] | >MCM4 | 845 | 859 | [KGILLQLFGGTNK] | >MCM4 | 538 | 550 | K10 | K1 |
| 101 | 595.007019 | 3 | -0.402729882 | [NINKLSK] | >MCM3 | 349 | 355 | [SPQKSPK] | >MCM3 | 761 | 767 | K4 | K4 |
| 101 | 682.74121 | 3 | -2.59067368 | [MLSLLFKK] | >CDC45 | 44 | 51 | [KVELLNR] | >CDC45 | 488 | 494 | K7 | K1 |
| 101 | 680.74403 | 3 | -0.005911134 | [ALTAAVKISER] | >CTF4 | 894 | 904 | [AILNKK] | >CTF4 | 792 | 797 | K7 | K5 |
| 101 | 608.5762879 | 4 | -0.305645168 | [LSFKGSFGAHALSPR] | >MCM3 | 155 | 169 | [STGVAAR] | >MCM3 | 333 | 339 | K4 | S1 |
| 101 | 593.663089 | 3 | -0.679099028 | [KQILDHQK] | >MRC1 | 482 | 489 | [SKDPK] | >MRC1 | 462 | 466 | K1 | S1 |
| 101 | 805.7940508 | 3 | -0.156771168 | [LLEENKAILNK] | >CTF4 | 786 | 796 | [MPVFVKSK] | >CTF4 | 778 | 785 | K6 | K6 |
| 101 | 1401.723077 | 3 | 1.209429676 | [YDDLKSPGDNIDFQTTILSR] | >MCM5 | 530 | 549 | [FSQLALDKALYALEK] | >MCM5 | 743 | 757 | Y1 | K8 |
| 100 | 593.6637023 | 3 | 0.355187368 | [KQILDHQK] | >MRC1 | 482 | 489 | [SKDPK] | >MRC1 | 462 | 466 | K1 | S1 |
| 100 | 617.0230103 | 3 | 0.165160088 | [SYKTVGSSK] | >MRC1 | 1033 | 1041 | [LIAPKR] | >MRC1 | 1053 | 1058 | K3 | K5 |
| 100 | 1146.595712 | 4 | -1.410779115 | [FSAFQEBKIQELSQQVPVGHIPR] | >MCM7 | 307 | 329 | [GISKTTEGLNSGVTGLR] | >MCM6 | 425 | 441 | K8 | S3 |
| 100 | 956.1922854 | 3 | -0.3337168 | [LQQEDKVIVLGEGVR] | >MCM4 | 910 | 924 | [KLQEDLSR] | >MCM4 | 860 | 867 | K6 | K1 |
| 100 | 989.5288007 | 3 | -1.218473775 | [SKEDESPTTKIFTIIK] | >MCM7 | 734 | 749 | [KMLQETGK] | >MCM7 | 750 | 757 | T8 | K1 |
| 100 | 708.1363473 | 4 | -0.774844229 | [ALYALEKHETIQLR] | >MCM5 | 751 | 764 | [FEQELKR] | >MCM5 | 714 | 720 | T10 | K6 |
| 99 | 759.4259524 | 3 | 1.069160143 | [NILKEVSNLR] | >PSF1 | 39 | 48 | [KNTEYLK] | >PSF1 | 49 | 55 | K4 | K1 |
| 99 | 1196.286877 | 3 | 1.099330697 | [LFASABSDQNVEKALSLAHELK] | >CTF4 | 869 | 890 | [SKLLEENK] | >CTF4 | 784 | 791 | K13 | S1 |
| 99 | 928.1492613 | 3 | 0.757365883 | [SFKNFILEFR] | >MCM5 | 30 | 39 | [IQYEKTHPDR] | >PSF2 | 102 | 111 | K3 | K5 |
| 98 | 687.0187327 | 3 | 0.101974625 | [KLQEDLSR] | >MCM4 | 860 | 867 | [VDEKNDR] | >MCM4 | 712 | 718 | K1 | K4 |
| 98 | 1067.528691 | 3 | 0.01970898 | [EMDSKFSFGQATPR] | >MCM7 | 674 | 687 | [TGKIDMNLVQTGK] | >MCM4 | 842 | 854 | K5 | T11 |
| 98 | 989.530833 | 3 | 0.836660976 | [SKEDESPTTKIFTIIK] | >MCM7 | 734 | 749 | [KMLQETGK] | >MCM7 | 750 | 757 | T8 | K1 |
| 98 | 1102.263181 | 3 | 0.08543093 | [KQLLINELESTER] | >MCM5 | 631 | 643 | [NTLSYENIVKTVR] | >MCM7 | 758 | 770 | K1 | K10 |
| 98 | 1580.703967 | 3 | 1.918673676 | [KYNFEDEEDFIDDDDGAGYISGK] | >CTF4 | 425 | 447 | [KDANLDYYNFNPMGIK] | >CTF4 | 670 | 685 | K1 | Y7 |
| 97 | 697.0303955 | 3 | -0.248190991 | [SKEDESPTTK] | >MCM7 | 734 | 743 | [KSPITAR] | >MCM3 | 694 | 700 | T8 | K1 |
| 97 | 679.0072632 | 3 | 1.074610429 | [VDDVTGEKVR] | >MCM6 | 103 | 112 | [SSASASGR] | >MCM4 | 96 | 103 | K8 | S6 |
| 97 | 956.1934189 | 3 | 0.852602121 | [LQQEDKVIVLGEGVR] | >MCM4 | 910 | 924 | [KLQEDLSR] | >MCM4 | 860 | 867 | K6 | S7 |
| 97 | 734.5965597 | 5 | -3.346921501 | [VHKTHSFPISLANTGK] | >CTF4 | 458 | 473 | [VHKTHSFPISLANTGK] | >CTF4 | 458 | 473 | K3 | K3 |
| 97 | 682.7421206 | 3 | -1.255730218 | [MLSLLFKK] | >CDC45 | 44 | 51 | [KVELLNR] | >CDC45 | 488 | 494 | K7 | K1 |
| 97 | 1198.23245 | 3 | -1.081147493 | [SDAAYFKDLDNNASDK] | >TOF1 | 984 | 999 | [KYVSQFSDYFLAR] | >TOF1 | 736 | 748 | K7 | K1 |
| 97 | 647.0092103 | 3 | -0.008983952 | [QLIDKEK] | >MRC1 | 687 | 693 | [EKEHEAK] | >MRC1 | 700 | 706 | K5 | K2 |

| | | | | | | | | | | | | |
|---|---|---|---|---|---|---|---|---|---|---|---|---|
| 97 | 556.981785 | 3 | -0.677062224 | [ILKSYK] | >MRC1 | 1030 | 1035 | [FKEGNK] | >MRC1 | 1021 | 1026 | K3 | K2 |
| 97 | 780.128884 | 3 | 0.717665935 | [ALTAAVKISER] | >CTF4 | 894 | 904 | [AELPSLVKK] | >CTF4 | 905 | 913 | K7 | K8 |
| 97 | 575.3585205 | 3 | 1.666321543 | [LIKHLR] | >CDC45 | 518 | 523 | [TKIYPK] | >MCM2 | 772 | 777 | K3 | T1 |
| 97 | 548.6793803 | 3 | 1.764226512 | [ILKSYK] | >MRC1 | 1030 | 1035 | [LIAPKR] | >MRC1 | 1053 | 1058 | K3 | K5 |
| 97 | 759.4242658 | 3 | -1.153625695 | [NILKEVSNLR] | >PSF1 | 39 | 48 | [KNTEYLK] | >PSF1 | 49 | 55 | K4 | K1 |
| 97 | 928.1490687 | 3 | 0.549680869 | [SFKNFILEFR] | >MCM5 | 30 | 39 | [IQYEKTHPDR] | >PSF2 | 102 | 111 | K3 | K5 |
| 97 | 1011.846447 | 3 | 1.163356242 | [KDANLDYYNFNPMGIK] | >CTF4 | 670 | 685 | [MPVFVKSK] | >CTF4 | 778 | 785 | K1 | K6 |
| 96 | 1072.927771 | 3 | -1.604653734 | [VHKTHSFPISLANTGK] | >CTF4 | 458 | 473 | [LLEENKAILNK] | >CTF4 | 786 | 796 | K3 | K6 |
| 96 | 1217.279273 | 3 | -0.898385966 | [SBIELTHSVLKVLEQYSDDK] | >TOF1 | 590 | 609 | [SKIDNSADIK] | >TOF1 | 490 | 499 | K11 | S1 |
| 96 | 950.8400394 | 3 | 2.09375841 | [SITTSTSPEQTER] | >MCM6 | 249 | 261 | [FLPFLQKGLR] | >MCM6 | 177 | 186 | S1 | K7 |
| 96 | 765.4006852 | 3 | 2.875597566 | [NILKEVSNLR] | >PSF1 | 39 | 48 | {mYGDLGNK] | >PSF1 | 0 | 8 | K4 | {0 |
| 96 | 540.3236694 | 3 | 2.339624384 | [FKEGNK] | >MRC1 | 1021 | 1026 | [TVKILK] | >MRC1 | 1027 | 1032 | K2 | K3 |
| 95 | 700.3293905 | 3 | 1.26064948 | [DYATDPKTGK] | >MCM4 | 835 | 844 | [KMGDDSR] | >MCM4 | 779 | 785 | K7 | K1 |
| 95 | 1389.339178 | 3 | -0.849224069 | [NTEYLKEQQQLGMLDDK] | >PSF1 | 50 | 66 | [VAKBQYFVTLLBMER] | >PSF1 | 67 | 81 | K6 | K3 |
| 95 | 1190.292313 | 3 | 2.53078405 | [FSQLALDKALYALEK] | >MCM5 | 743 | 757 | [SPGDNIDFQTTILSR] | >MCM5 | 535 | 549 | K8 | S1 |
| 95 | 1196.286962 | 3 | 1.170390256 | [LFASABSDQNVEKALSLAHELK] | >CTF4 | 869 | 890 | [SKLLEENK] | >CTF4 | 784 | 791 | K13 | S1 |
| 95 | 765.400611 | 3 | 2.778465409 | [NILKEVSNLR] | >PSF1 | 39 | 48 | {mYGDLGNK] | >PSF1 | 0 | 8 | S7 | {0 |
| 94 | 1094.299391 | 4 | 0.718144443 | [GKHIWPEFPLPLPSEMEIR] | >CTF4 | 759 | 777 | [KDANLDYYNFNPMGIK] | >CTF4 | 670 | 685 | K2 | K1 |
| 94 | 729.9425375 | 4 | -0.898538273 | [SQILQYVHKITPR] | >MCM4 | 575 | 587 | [LINLKGLVLR] | >MCM4 | 325 | 334 | K9 | K5 |
| 94 | 584.0817594 | 4 | 0.174636391 | [ALSLAHELKQDR] | >CTF4 | 882 | 893 | [KINNIR] | >CTF4 | 913 | 918 | K9 | K1 |
| 94 | 736.1435168 | 4 | 1.036204614 | [SLYKTYVDVVHVK] | >MCM4 | 455 | 467 | [BSFADKQVIK] | >MCM4 | 389 | 398 | Y6 | K6 |
| 94 | 1182.960565 | 3 | 1.888683655 | [THSFPISLANTGKFR] | >CTF4 | 461 | 475 | [THSFPISLANTGKFR] | >CTF4 | 461 | 475 | T11 | K13 |
| 94 | 553.0864258 | 4 | 1.163699913 | [LINLKGLVLR] | >MCM4 | 325 | 334 | [VGDGMKIR] | >MCM7 | 444 | 451 | K5 | K6 |
| 93 | 919.7383995 | 4 | 2.431826966 | [EIDYKDDVLDVILNQR] | >MCM7 | 131 | 146 | [FGGLLSIQTPDKTR] | >TOF1 | 360 | 373 | Y4 | K12 |
| 93 | 1395.751194 | 3 | 0.49627273 | [LYEILTNSIAPSIFGNEDIKK] | >MCM5 | 366 | 386 | [FVEKVSPIAVYTSGK] | >MCM5 | 428 | 442 | S12 | K4 |
| 93 | 1102.262328 | 3 | -0.689297084 | [KQLLINELESTER] | >MCM5 | 631 | 643 | [NTLSYENIVKTVR] | >MCM7 | 758 | 770 | K1 | K10 |
| 92 | 570.3017511 | 4 | -0.358157892 | [NILKEVSNLR] | >PSF1 | 39 | 48 | {MYGDLGNK] | >PSF1 | 0 | 8 | S7 | {0 |
| 92 | 912.513277 | 3 | -1.787890625 | [GISKTTEGLNSGVTGLR] | >MCM6 | 425 | 441 | [VLKSLYK] | >MCM4 | 452 | 458 | S3 | K3 |
| 92 | 1089.589478 | 3 | 0.978772848 | [DNMVLKLVLNFFR] | >TOF1 | 198 | 210 | [WFKLVDDQQKR] | >TOF1 | 61 | 71 | K6 | K3 |
| 92 | 591.3342018 | 3 | -1.797692708 | [STVEGIKLR] | >MCM6 | 558 | 566 | [SKDPK] | >MRC1 | 462 | 466 | K7 | S1 |
| 92 | 738.022717 | 3 | -0.650205764 | [SAIKDYATDPK] | >MCM4 | 831 | 841 | [KMGDDSR] | >MCM4 | 779 | 785 | K4 | K1 |
| 92 | 565.9795532 | 3 | -2.045056135 | [EKEHEAK] | >MRC1 | 700 | 706 | [IKELK] | >MRC1 | 707 | 711 | K2 | K2 |
| 92 | 854.1533014 | 3 | -0.140483373 | [ALSLAHELKQDR] | >CTF4 | 882 | 893 | [AELPSLVKK] | >CTF4 | 905 | 913 | K9 | K8 |
| 92 | 729.9431377 | 4 | -0.075402398 | [SQILQYVHKITPR] | >MCM4 | 575 | 587 | [LINLKGLVLR] | >MCM4 | 325 | 334 | K9 | K5 |
| 91 | 556.9816911 | 3 | -0.845912986 | [ILKSYK] | >MRC1 | 1030 | 1035 | [FKEGNK] | >MRC1 | 1021 | 1026 | K3 | K2 |
| 91 | 633.3628509 | 3 | -2.078138155 | [GLKLEDMAK] | >MRC1 | 496 | 504 | [LVKNPK] | >MRC1 | 839 | 844 | K3 | K3 |
| 91 | 970.8568536 | 3 | -1.996893371 | [FGGLLSIQTPDKTR] | >TOF1 | 360 | 373 | [LLQKEHQMR] | >TOF1 | 338 | 346 | K12 | K4 |
| 91 | 1067.526318 | 3 | -2.204689272 | [EMDSKFSFGQATPR] | >MCM7 | 674 | 687 | [TGKIDMNLVQTGK] | >MCM4 | 842 | 854 | K5 | T11 |
| 91 | 984.3229134 | 5 | 2.36187456 | [TAIHEVMEQQTISISKAGINTTLNAR] | >MCM7 | 535 | 560 | [SVYTSGKASSAAGLTAAVVR] | >MCM6 | 595 | 614 | K16 | S5 |
| 91 | 901.4874555 | 4 | -1.373396862 | [SVYTSGKASSAAGLTAAVVR] | >MCM6 | 595 | 614 | [SQFLKYVVGFAPR] | >MCM6 | 582 | 594 | Y3 | K5 |
| 91 | 817.1142454 | 3 | -0.610334707 | [KFNISEGDITK] | >TOF1 | 622 | 632 | [TLVIEGKSR] | >TOF1 | 610 | 618 | K1 | K7 |
| 91 | 1037.55605 | 3 | 1.293467715 | [DNMVLKLVLNFFR] | >TOF1 | 198 | 210 | [WFKLVDDQQK] | >TOF1 | 61 | 70 | K6 | K3 |
| 91 | 917.8663961 | 3 | -1.207736792 | [FNPLLQAGAKLAK] | >MCM3 | 625 | 637 | [NILKEVSNLR] | >PSF1 | 39 | 48 | K10 | K4 |
| 91 | 919.7377841 | 4 | 1.762136717 | [EIDYKDDVLDVILNQR] | >MCM7 | 131 | 146 | [FGGLLSIQTPDKTR] | >TOF1 | 360 | 373 | Y4 | K12 |
| 90 | 708.1365832 | 4 | -0.441240215 | [ALYALEKHETIQLR] | >MCM5 | 751 | 764 | [FEQELKR] | >MCM5 | 714 | 720 | T10 | K6 |
| 90 | 1150.866473 | 3 | 0.397251496 | [GVBLIDEFDKMNDQDR] | >MCM2 | 602 | 617 | [BSIIAAANPNGGR] | >MCM2 | 644 | 656 | K10 | S2 |
| 90 | 844.4688816 | 3 | -2.346839124 | [FLPFLQKGLR] | >MCM6 | 177 | 186 | [VDDVTGEKVR] | >MCM6 | 103 | 112 | K7 | K8 |
| 90 | 631.1351677 | 4 | -1.585409529 | [TYVDVVHVKK] | >MCM4 | 459 | 468 | [LINLKGLVLR] | >MCM4 | 325 | 334 | K9 | K5 |
| 90 | 1037.556379 | 3 | 1.610706 | [DNMVLKLVLNFFR] | >TOF1 | 198 | 210 | [WFKLVDDQQK] | >TOF1 | 61 | 70 | K6 | K3 |
| 90 | 649.6802222 | 3 | -1.695149324 | [EKADESLPK] | >MRC1 | 617 | 625 | [KSHHVK] | >MRC1 | 594 | 599 | K2 | K1 |
| 90 | 759.4254363 | 3 | 0.38903698 | [NILKEVSNLR] | >PSF1 | 39 | 48 | [KNTEYLK] | >PSF1 | 49 | 55 | K4 | K1 |
| 90 | 802.1130622 | 3 | 0.483670127 | [FLPFLQKGLR] | >MCM6 | 177 | 186 | [KVDDVTGEK] | >MCM6 | 102 | 110 | K7 | T6 |
| 90 | 707.9094735 | 4 | 1.972692607 | [DPKVDHNVLLNTLR] | >MRC1 | 464 | 477 | [ATILNLKAR] | >MRC1 | 439 | 447 | K3 | K7 |
| 89 | 811.4095436 | 3 | 0.607796005 | [EMVKDEHIYDK] | >MCM6 | 518 | 528 | [KIAEVDR] | >MCM6 | 919 | 925 | K4 | K1 |
| 89 | 522.6836133 | 3 | 0.646166381 | [TVKILK] | >MRC1 | 1027 | 1032 | [KLIAPK] | >MRC1 | 1052 | 1057 | K3 | K1 |
| 89 | 608.3384 | 3 | -0.817251493 | [EKEHEAK] | >MRC1 | 700 | 706 | [LQLKQK] | >MRC1 | 694 | 699 | K2 | K4 |
| 89 | 995.1974904 | 3 | 1.193124529 | [IBQTVGKTDPVLR] | >CSM3 | 114 | 126 | [IGSKYSLQFMK] | >TOF1 | 579 | 589 | K7 | S3 |
| 89 | 1225.980414 | 3 | 1.049958371 | [EIDYKDDVLDVILNQR] | >MCM7 | 131 | 146 | [FGGLLSIQTPDKTR] | >TOF1 | 360 | 373 | Y4 | K12 |
| 88 | 684.051692 | 3 | 0.078711677 | [KQILDHQK] | >MRC1 | 482 | 489 | [LSKQNQK] | >MRC1 | 448 | 454 | K1 | K3 |
| 88 | 591.6479025 | 3 | 0.102920074 | [YVEKTAHR] | >MCM2 | 555 | 562 | [SKDPK] | >MRC1 | 462 | 466 | K4 | S1 |
| 88 | 676.9995229 | 3 | -0.372772774 | [SHESYKDTK] | >PSF3 | 180 | 188 | [WMFKK} | >PSF3 | 190 | 195 | Y5 | K4 |
| 88 | 759.4255471 | 3 | 0.534982332 | [NILKEVSNLR] | >PSF1 | 39 | 48 | [KNTEYLK] | >PSF1 | 49 | 55 | K4 | K1 |
| 88 | 783.4184311 | 3 | -2.075570837 | [VPLLSSYANNLKR] | >MRC1 | 383 | 395 | [EIDSSK] | >MRC1 | 396 | 401 | K12 | S4 |
| 88 | 1011.843094 | 3 | -2.153211896 | [KDANLDYYNFNPMGIK] | >CTF4 | 670 | 685 | [MPVFVKSK] | >CTF4 | 778 | 785 | K1 | K6 |
| 88 | 680.7348022 | 3 | -1.246624178 | [ATILNLKAR] | >MRC1 | 439 | 447 | [LSKQNQK] | >MRC1 | 448 | 454 | K7 | K3 |
| 88 | 1040.917745 | 3 | 0.367467487 | [LTVSGSQALVDEKIALQK] | >TOF1 | 374 | 391 | [NVIKHTSAR] | >TOF1 | 348 | 356 | K13 | K4 |
| 88 | 765.4011207 | 3 | 3.445008522 | [NILKEVSNLR] | >PSF1 | 39 | 48 | {mYGDLGNK] | >PSF1 | 0 | 8 | S7 | {0 |
| 88 | 1395.752088 | 3 | 1.137064554 | [LYEILTNSIAPSIFGNEDIKK] | >MCM5 | 366 | 386 | [FVEKVSPIAVYTSGK] | >MCM5 | 428 | 442 | S12 | K4 |
| 87 | 538.9820313 | 3 | -0.65625692 | [FKEGNK] | >MRC1 | 1021 | 1026 | [LIAPKR] | >MRC1 | 1053 | 1058 | K2 | K5 |
| 87 | 817.1141853 | 3 | -0.684012104 | [KFNISEGDITK] | >TOF1 | 622 | 632 | [TLVIEGKSR] | >TOF1 | 610 | 618 | K1 | K7 |
| 87 | 1118.260509 | 3 | 1.04363352 | [ELKSFLLEYTDETGR] | >MCM2 | 210 | 224 | [RLEIGDDAKLVK] | >MRC1 | 830 | 841 | S4 | K9 |
| 87 | 1483.359238 | 3 | -0.566983855 | [YDDLKSPGDNIDFQTTILSR] | >MCM5 | 530 | 549 | [GVBLIDEFDKMNDQDR] | >MCM2 | 602 | 617 | Y1 | K10 |
| 87 | 420.4942017 | 4 | -2.919012163 | [KTEGSHR] | >MRC1 | 1059 | 1065 | [KLIAPK] | >MRC1 | 1052 | 1057 | S5 | K1 |
| 87 | 742.4002822 | 4 | 1.292953807 | [SKEDESPTTKIFTIIK] | >MCM7 | 734 | 749 | [KMLQETGK] | >MCM7 | 750 | 757 | T8 | K1 |
| 87 | 482.6061168 | 3 | -2.26075627 | [SDEKR] | >MCM4 | 786 | 790 | [KLSLR] | >MCM6 | 692 | 696 | K4 | K1 |
| 87 | 696.3637528 | 4 | 0.739812914 | [SFKNFILEFR] | >MCM5 | 30 | 39 | [IQYEKTHPDR] | >PSF2 | 102 | 111 | K3 | K5 |
| 87 | 1099.255198 | 3 | 0.496596976 | [FDLVYLVLDKVDEK] | >MCM4 | 702 | 715 | [TGKIDMNLVQTGK] | >MCM4 | 842 | 854 | K10 | T1 |

| | | | | | | | | | | | | | |
|---|---|---|---|---|---|---|---|---|---|---|---|---|---|
| 86 | 747.8569076 | 4 | 0.538146459 | [KAPEQNHNNGKDR] | >MRC1 | 71 | 83 | [SEVKDNSYSEK] | >MRC1 | 106 | 116 | K11 | K4 |
| 86 | 586.0040344 | 3 | 0.698759517 | [LSKTVNK] | >MCM3 | 717 | 723 | [SPQKSPK] | >MCM3 | 761 | 767 | T4 | K4 |
| 86 | 1102.263181 | 3 | 0.085370411 | [KQLLINELESTER] | >MCM5 | 631 | 643 | [NTLSYENIVKTVR] | >MCM7 | 758 | 770 | K1 | K10 |
| 86 | 1025.195823 | 3 | -2.597722656 | [AIKVVVDSFVDAQK] | >MCM2 | 837 | 850 | [LHQMDMDKVSR] | >MCM2 | 778 | 788 | K3 | K8 |
| 86 | 765.3969151 | 3 | -2.054484749 | [NILKEVSNLR] | >PSF1 | 39 | 48 | {mYGDLGNK] | >PSF1 | 0 | 8 | S7 | {0 |
| 86 | 676.9996893 | 3 | -0.126822062 | [SHESYKDTK] | >PSF3 | 180 | 188 | [WMFKK} | >PSF3 | 190 | 195 | Y5 | K4 |
| 86 | 470.0241099 | 4 | 0.197690597 | [TDINDKFK] | >MRC1 | 1015 | 1022 | [TVKILK] | >MRC1 | 1027 | 1032 | K6 | K3 |
| 85 | 964.8546127 | 3 | -2.936969836 | [VHKTHSFPISLANTGK] | >CTF4 | 458 | 473 | [SKLLEENK] | >CTF4 | 784 | 791 | K3 | S1 |
| 85 | 776.1046169 | 3 | -2.469905745 | [TVGSSKASITYMGK] | >MRC1 | 1036 | 1049 | [TVKILK] | >MRC1 | 1027 | 1032 | K6 | K3 |
| 85 | 1225.981373 | 3 | 1.832423576 | [EIDYKDDVLDVILNQR] | >MCM7 | 131 | 146 | [FGGLLSIQTPDKTR] | >TOF1 | 360 | 373 | Y4 | K12 |
| 85 | 596.3187726 | 3 | -3.485239487 | [LVDDQQKR] | >TOF1 | 64 | 71 | [KSMPK] | >TOF1 | 226 | 230 | K7 | K1 |
| 85 | 649.6815196 | 3 | 0.303955254 | [EKADESLPK] | >MRC1 | 617 | 625 | [KSHHVK] | >MRC1 | 594 | 599 | K2 | K1 |
| 85 | 538.9819203 | 3 | -0.862624979 | [FKEGNK] | >MRC1 | 1021 | 1026 | [LIAPKR] | >MRC1 | 1053 | 1058 | K2 | K5 |
| 85 | 780.128769 | 3 | 0.570144588 | [ALTAAVKISER] | >CTF4 | 894 | 904 | [AELPSLVKK] | >CTF4 | 905 | 913 | K7 | S5 |
| 85 | 528.3348621 | 3 | 0.619176356 | [LQLKQK] | >MRC1 | 694 | 699 | [IKELK] | >MRC1 | 707 | 711 | K4 | K2 |
| 84 | 1001.215105 | 3 | -1.643699839 | [ELGIIFDKNLDR] | >CDC45 | 387 | 398 | [LYPLLQDEVKR] | >CDC45 | 291 | 301 | K8 | K10 |
| 84 | 800.8995548 | 4 | 1.540938068 | [EMDSKFSFGQATPR] | >MCM7 | 674 | 687 | [TGKIDMNLVQTGK] | >MCM4 | 842 | 854 | K5 | T11 |
| 84 | 631.1371282 | 4 | 1.524600166 | [TYVDVVHVKK] | >MCM4 | 459 | 468 | [LINLKGLVLR] | >MCM4 | 325 | 334 | K9 | K5 |
| 83 | 540.6367723 | 3 | 1.22926428 | [LKBAPR] | >MCM5 | 608 | 613 | [EIYKK] | >PSF3 | 175 | 179 | K2 | K4 |
| 83 | 663.3318535 | 3 | 0.832553225 | {MYGDLGNK] | >PSF1 | 0 | 8 | [KNTEYLK] | >PSF1 | 49 | 55 | {0 | K1 |
| 83 | 797.4105295 | 4 | 0.35669766 | [VHKTHSFPISLANTGK] | >CTF4 | 458 | 473 | [KPHNEHSYSR] | >CTF4 | 448 | 457 | T4 | K1 |
| 83 | 800.8965578 | 4 | -2.204661152 | [EMDSKFSFGQATPR] | >MCM7 | 674 | 687 | [TGKIDMNLVQTGK] | >MCM4 | 842 | 854 | K5 | T11 |
| 83 | 811.6353245 | 4 | 1.831618418 | [ESLYQETNKSKEDESPTTK] | >MCM7 | 725 | 743 | [NDDNTKK] | >MCM3 | 688 | 694 | T17 | K6 |
| 83 | 1094.2997 | 4 | 1.000366475 | [GKHIWPEFPLPLPSEMEIR] | >CTF4 | 759 | 777 | [KDANLDYYNFNPMGIK] | >CTF4 | 670 | 685 | K2 | K1 |
| 82 | 806.1381836 | 4 | -3.422832288 | [EMDSKFSFGQATPR] | >MCM7 | 674 | 687 | [VSKESLYQETNK] | >MCM7 | 722 | 733 | K5 | S2 |
| 82 | 490.6297183 | 3 | -3.561535108 | [LIAPKR] | >MRC1 | 1053 | 1058 | [TKSNK] | >MRC1 | 1081 | 1085 | K5 | K2 |
| 82 | 618.3636216 | 3 | 1.118523823 | [SYKTVGSSK] | >MRC1 | 1033 | 1041 | [TVKILK] | >MRC1 | 1027 | 1032 | K3 | T1 |
| 82 | 575.3579712 | 3 | 0.710468577 | [LIKHLR] | >CDC45 | 518 | 523 | [TKIYPK] | >MCM2 | 772 | 777 | K3 | T1 |
| 82 | 688.0750523 | 3 | 0.635754659 | [mLSLLFKK] | >CDC45 | 44 | 51 | [KVELLNR] | >CDC45 | 488 | 494 | K7 | K1 |
| 82 | 841.1793202 | 3 | 0.225482737 | [TYVDVVHVKK] | >MCM4 | 459 | 468 | [LINLKGLVLR] | >MCM4 | 325 | 334 | K9 | K5 |
| 82 | 546.5888672 | 3 | -1.663084471 | [KMGDDSR] | >MCM4 | 779 | 785 | [SDEKR] | >MCM4 | 786 | 790 | K1 | K4 |
| 82 | 928.1500908 | 3 | 1.651736271 | [SFKNFILEFR] | >MCM5 | 30 | 39 | [IQYEKTHPDR] | >PSF2 | 102 | 111 | K3 | K5 |
| 81 | 570.5434502 | 4 | 0.259644819 | [DYATDPKTGK] | >MCM4 | 835 | 844 | [KLQEDLSR] | >MCM4 | 860 | 867 | K7 | K1 |
| 81 | 764.1776758 | 4 | 1.124335236 | [SKDPKVDHNVLLNTLR] | >MRC1 | 462 | 477 | [KQILDHQK] | >MRC1 | 482 | 489 | T14 | K1 |
| 81 | 804.9476412 | 4 | -1.612043148 | [VHKTHSFPISLANTGK] | >CTF4 | 458 | 473 | [LLEENKAILNK] | >CTF4 | 786 | 796 | K3 | K6 |
| 81 | 759.4256344 | 3 | 0.650096418 | [NILKEVSNLR] | >PSF1 | 39 | 48 | [KNTEYLK] | >PSF1 | 49 | 55 | K4 | K1 |
| 81 | 841.1794047 | 3 | 0.325941815 | [TYVDVVHVKK] | >MCM4 | 459 | 468 | [LINLKGLVLR] | >MCM4 | 325 | 334 | K9 | K5 |
| 81 | 1395.751868 | 3 | 0.979456129 | [LYEILTNSIAPSIFGNEDIKK] | >MCM5 | 366 | 386 | [FVEKVSPIAVYTSGK] | >MCM5 | 428 | 442 | S12 | K4 |
| 81 | 977.8712098 | 3 | 0.63923816 | {MYGDLGNKLVLEAK] | >PSF1 | 0 | 14 | [NILKEVSNLR] | >PSF1 | 39 | 48 | K8 | K4 |
| 80 | 540.6352539 | 3 | -1.582692705 | [LKBAPR] | >MCM5 | 608 | 613 | [EIYKK] | >PSF3 | 175 | 179 | K2 | K4 |
| 80 | 585.6657858 | 3 | 1.629423372 | [TKQLYAR] | >PSF1 | 16 | 22 | [EIYKK] | >PSF3 | 175 | 179 | Y5 | K4 |
| 80 | 540.6357313 | 3 | -0.698509254 | [LKBAPR] | >MCM5 | 608 | 613 | [EIYKK] | >PSF3 | 175 | 179 | K2 | K4 |
| 80 | 765.4000159 | 3 | 2.000348276 | [NILKEVSNLR] | >PSF1 | 39 | 48 | {mYGDLGNK] | >PSF1 | 0 | 8 | K4 | {0 |
| 80 | 652.7138872 | 3 | -2.34441309 | [TFKPILTK] | >MCM6 | 760 | 767 | [YIKYAR] | >MCM6 | 754 | 759 | K3 | K3 |
| 80 | 1203.307331 | 3 | -1.598752212 | [VHKTHSFPISLANTGK] | >CTF4 | 458 | 473 | [THSFPISLANTGKFR] | >CTF4 | 461 | 475 | T4 | K13 |
| 80 | 928.1461876 | 3 | -2.55662328 | [SFKNFILEFR] | >MCM5 | 30 | 39 | [IQYEKTHPDR] | >PSF2 | 102 | 111 | K3 | K5 |
| 80 | 1133.370921 | 4 | 1.708534416 | [SVLHEVMEQQTISIAKAGIITTLNAR] | >MCM4 | 643 | 668 | [SQFLKYVVGFAPR] | >MCM6 | 582 | 594 | K16 | K5 |
| 80 | 715.7597479 | 3 | 0.245004732 | [STVEGIKLR] | >MCM6 | 558 | 566 | [TFKPILTK] | >MCM6 | 760 | 767 | K7 | K3 |
| 80 | 759.425995 | 3 | 1.125370624 | [NILKEVSNLR] | >PSF1 | 39 | 48 | [KNTEYLK] | >PSF1 | 49 | 55 | K4 | K1 |
| 80 | 1139.624041 | 3 | 1.153647736 | [ALYALEKHETIQLR] | >MCM5 | 751 | 764 | [NTLSYENIVKTVR] | >MCM7 | 758 | 770 | K7 | K10 |
| 80 | 1051.546215 | 4 | 3.196556951 | [YDDLKSPGDNIDFQTTILSR] | >MCM5 | 530 | 549 | [FSQLALDKALYALEK] | >MCM5 | 743 | 757 | Y1 | K8 |
| 79 | 660.3784298 | 3 | -0.575119031 | [LLQKEHQMR] | >TOF1 | 338 | 346 | [KNVIK] | >TOF1 | 347 | 351 | K4 | K1 |
| 79 | 470.0237122 | 4 | -0.649862838 | [TDINDKFK] | >MRC1 | 1015 | 1022 | [TVKILK] | >MRC1 | 1027 | 1032 | K6 | K3 |
| 79 | 540.3237101 | 3 | 2.414894911 | [FKEGNK] | >MRC1 | 1021 | 1026 | [TVKILK] | >MRC1 | 1027 | 1032 | K2 | K3 |
| 79 | 898.9261567 | 4 | -1.081125226 | [SDAAYFKDLDNNASDK] | >TOF1 | 984 | 999 | [KYVSQFSDYFLAR] | >TOF1 | 736 | 748 | K7 | K1 |
| 79 | 553.0855569 | 4 | -0.409388335 | [LINLKGLVLR] | >MCM4 | 325 | 334 | [VGDGMKIR] | >MCM7 | 444 | 451 | K5 | K6 |
| 79 | 1099.239976 | 3 | 1.903701723 | [DNMVLKLVLNFFR] | >TOF1 | 198 | 210 | [EIQSELASEDSKR] | >MRC1 | 293 | 305 | K6 | S8 |
| 79 | 630.040218 | 3 | 0.329112861 | [KPQKPIPTK] | >MRC1 | 315 | 323 | [KFFSK] | >MRC1 | 324 | 328 | K4 | K1 |
| 79 | 663.8763724 | 4 | 0.18888712 | [ALYALEKHETIQLR] | >MCM5 | 751 | 764 | [KSPITAR] | >MCM3 | 694 | 700 | K7 | K1 |
| 78 | 764.1759699 | 4 | -1.110187618 | [SKDPKVDHNVLLNTLR] | >MRC1 | 462 | 477 | [KQILDHQK] | >MRC1 | 482 | 489 | K5 | K1 |
| 78 | 1023.857388 | 3 | -0.766494817 | [FGGLLSIQTPDKTR] | >TOF1 | 360 | 373 | [NDIHTSDLSSPR] | >MCM4 | 136 | 147 | K12 | T5 |
| 78 | 943.8470883 | 3 | 0.339229878 | [ALYALEKHETIQLR] | >MCM5 | 751 | 764 | [FEQELKR] | >MCM5 | 714 | 720 | K7 | K6 |
| 78 | 759.4254539 | 3 | 0.412131854 | [NILKEVSNLR] | >PSF1 | 39 | 48 | [KNTEYLK] | >PSF1 | 49 | 55 | K4 | K1 |
| 78 | 941.724635 | 4 | 0.267968395 | [TSEVRPELYKASFTBDMBR] | >MCM6 | 297 | 315 | [FLPFLQKGLR] | >MCM6 | 177 | 186 | T1 | K7 |
| 78 | 1081.901307 | 3 | -0.478313372 | [THSFPISLANTGKFR] | >CTF4 | 461 | 475 | [THSFPISLANTGK] | >CTF4 | 461 | 473 | K13 | S7 |
| 77 | 1036.032556 | 4 | -0.095501002 | [YDDLKSPGDNIDFQTTILSR] | >MCM5 | 530 | 549 | [VVVDSFVDAQKVSVR] | >MCM2 | 840 | 854 | S19 | K11 |
| 77 | 797.4113365 | 4 | 1.369700555 | [VHKTHSFPISLANTGK] | >CTF4 | 458 | 473 | [KPHNEHSYSR] | >CTF4 | 448 | 457 | T4 | K1 |
| 77 | 902.734257 | 4 | 0.551642677 | [VHKTHSFPISLANTGK] | >CTF4 | 458 | 473 | [THSFPISLANTGKFR] | >CTF4 | 461 | 475 | T4 | K13 |
| 77 | 727.3694738 | 5 | 0.828662427 | [SKEDESPTTKIFTIIK] | >MCM7 | 734 | 749 | [EMDSKFSFGQATPR] | >MCM7 | 674 | 687 | K10 | S4 |
| 77 | 1395.748718 | 3 | -1.278930418 | [LYEILTNSIAPSIFGNEDIKK] | >MCM5 | 366 | 386 | [FVEKVSPIAVYTSGK] | >MCM5 | 428 | 442 | S12 | K4 |
| 76 | 715.7584241 | 3 | -1.606187056 | [STVEGIKLR] | >MCM6 | 558 | 566 | [TFKPILTK] | >MCM6 | 760 | 767 | K7 | K3 |
| 76 | 897.4658173 | 4 | -0.194148532 | [LFASABSDQNVEKALSLAHELK] | >CTF4 | 869 | 890 | [SKLLEENK] | >CTF4 | 784 | 791 | K13 | S1 |
| 76 | 977.8683991 | 3 | -2.237014234 | {MYGDLGNKLVLEAK] | >PSF1 | 0 | 14 | [NILKEVSNLR] | >PSF1 | 39 | 48 | K8 | K4 |
| 76 | 913.2128455 | 4 | 0.82387409 | [SBIELTHSVLKVLEQYSDDK] | >TOF1 | 590 | 609 | [SKIDNSADIK] | >TOF1 | 490 | 499 | K11 | S1 |
| 76 | 1168.134752 | 4 | -0.056757474 | [SAYDLKNWTVTHAGMIAFNELLNLVSR] | >TOF1 | 519 | 545 | [IQLLSNLPKIGSK] | >TOF1 | 570 | 582 | S26 | K9 |
| 76 | 427.496964 | 4 | 3.81E-04 | [KTEGSHR] | >MRC1 | 1059 | 1065 | [LIAPKR] | >MRC1 | 1053 | 1058 | S5 | K5 |
| 76 | 760.3962771 | 3 | 1.874253338 | [ASITYMGKTR] | >MRC1 | 1042 | 1051 | [SYKTVGSSK] | >MRC1 | 1033 | 1041 | K8 | K3 |

| | | | | | | | | | | | | |
|---|---|---|---|---|---|---|---|---|---|---|---|---|
| 76 | 1203.309954 | 3 | 0.582131181 | [VHKTHSFPISLANTGK] | >CTF4 | 458 | 473 | [THSFPISLANTGKFR] | >CTF4 | 461 | 475 | K3 | S7 |
| 76 | 1067.530193 | 3 | 1.42738056 | [EMDSKFSFGQATPR] | >MCM7 | 674 | 687 | [TGKIDMNLVQTGK] | >MCM4 | 842 | 854 | K5 | T11 |
| 76 | 1001.215142 | 3 | -1.606969745 | [ELGIIFDKNLDR] | >CDC45 | 387 | 398 | [LYPLLQDEVKR] | >CDC45 | 291 | 301 | K8 | K10 |
| 75 | 783.764 | 3 | -0.484889514 | [QNQKLSQRPNK] | >MRC1 | 451 | 461 | [YIKYAR] | >MCM6 | 754 | 759 | K4 | K3 |
| 75 | 786.654605 | 4 | -0.90319477 | [EDESPTTKIFTIIK] | >MCM7 | 736 | 749 | [ESLYQETNKSK] | >MCM7 | 725 | 735 | K8 | T7 |
| 75 | 777.1024296 | 3 | -2.540264651 | [VPLLSSYANNLKR] | >MRC1 | 383 | 395 | [INEKR] | >MRC1 | 378 | 382 | K12 | K4 |
| 75 | 625.3237849 | 3 | -4.597293807 | [SYKTVGSSK] | >MRC1 | 1033 | 1041 | [FKEGNK] | >MRC1 | 1021 | 1026 | K3 | K2 |
| 75 | 687.0172729 | 3 | -2.024922069 | [KLQEDLSR] | >MCM4 | 860 | 867 | [VDEKNDR] | >MCM4 | 712 | 718 | K1 | K4 |
| 75 | 747.8569483 | 4 | 0.592683866 | [KAPEQNHNNGKDR] | >MRC1 | 71 | 83 | [SEVKDNSYSEK] | >MRC1 | 106 | 116 | K11 | K4 |
| 75 | 1196.28634 | 3 | 0.649662962 | [LFASABSDQNVEKALSLAHELK] | >CTF4 | 869 | 890 | [SKLLEENK] | >CTF4 | 784 | 791 | K13 | S1 |
| 75 | 1248.671616 | 3 | 1.638215683 | [FSAFQEBKIQELSQQVPVGHIPR] | >MCM7 | 307 | 329 | [VLKSLYK] | >MCM4 | 452 | 458 | S13 | K3 |
| 74 | 699.7439167 | 3 | 2.784022565 | [ATILNLKAR] | >MRC1 | 439 | 447 | [LHKMFAR] | >CDC45 | 359 | 365 | K7 | K3 |
| 74 | 964.8557965 | 3 | -1.709161519 | [VHKTHSFPISLANTGK] | >CTF4 | 458 | 473 | [SKLLEENK] | >CTF4 | 784 | 791 | S10 | K2 |
| 74 | 794.7782207 | 3 | -2.13778156 | [SQILQYVHKITPR] | >MCM4 | 575 | 587 | [VSDKR] | >MCM4 | 469 | 473 | K9 | K4 |
| 74 | 590.0043288 | 3 | 0.333603237 | [ILFNKAK] | >PSF2 | 125 | 131 | [WMFKK} | >PSF3 | 190 | 195 | K5 | K4 |
| 74 | 637.3807373 | 3 | -2.318897367 | [EVIETKGLK] | >MRC1 | 490 | 498 | [LVKNPK] | >MRC1 | 839 | 844 | T5 | K3 |
| 74 | 1193.261319 | 3 | 1.745769303 | [SDAAYFKDLDNNASDK] | >TOF1 | 984 | 999 | [DNMVLKLVLNFFR] | >TOF1 | 198 | 210 | K7 | K6 |
| 74 | 777.1046702 | 3 | 0.345498728 | [VPLLSSYANNLKR] | >MRC1 | 383 | 395 | [INEKR] | >MRC1 | 378 | 382 | K12 | K4 |
| 74 | 1008.174556 | 3 | 2.600749809 | [LGDDBLABLKDLK] | >TOF1 | 47 | 59 | [WFKLVDDQQK] | >TOF1 | 61 | 70 | K10 | K3 |
| 74 | 913.2139484 | 4 | 2.032500676 | [SBIELTHSVLKVLEQYSDDK] | >TOF1 | 590 | 609 | [SKIDNSADIK] | >TOF1 | 490 | 499 | K11 | S1 |
| 73 | 642.9946407 | 3 | 1.326066745 | [KDDAQGFSR] | >MCM6 | 782 | 790 | [YKELR] | >MCM6 | 777 | 781 | K1 | K2 |
| 73 | 897.1389811 | 3 | 0.785940403 | [EIQSELASEDSKR] | >MRC1 | 293 | 305 | [TLVIEGKSR] | >TOF1 | 610 | 618 | S8 | K7 |
| 73 | 1048.537048 | 3 | -0.9032234 | [EDESPTTKIFTIIK] | >MCM7 | 736 | 749 | [ESLYQETNKSK] | >MCM7 | 725 | 735 | K8 | T7 |
| 73 | 1193.988172 | 3 | -0.761714557 | [IDMNLVQTGKSVIQR] | >MCM4 | 845 | 859 | [LQQEDKVIVLGEGVR] | >MCM4 | 910 | 924 | K10 | K6 |
| 73 | 574.6636963 | 3 | -1.785529556 | [EGNKTVK] | >MRC1 | 1023 | 1029 | [ILKSYK] | >MRC1 | 1030 | 1035 | K4 | K3 |
| 73 | 913.2139043 | 4 | 1.984243782 | [SBIELTHSVLKVLEQYSDDK] | >TOF1 | 590 | 609 | [SKIDNSADIK] | >TOF1 | 490 | 499 | K11 | S1 |
| 73 | 1168.13392 | 4 | -0.769277084 | [SAYDLKNWTVTHAGMIAFNELLNLVSR] | >TOF1 | 519 | 545 | [IQLLSNLPKIGSK] | >TOF1 | 570 | 582 | S26 | K9 |
| 73 | 1037.555726 | 3 | 0.9811028 | [DNMVLKLVLNFFR] | >TOF1 | 198 | 210 | [WFKLVDDQQK] | >TOF1 | 61 | 70 | K6 | K3 |
| 73 | 759.4249446 | 3 | -0.259056491 | [NILKEVSNLR] | >PSF1 | 39 | 48 | [KNTEYLK] | >PSF1 | 49 | 55 | K4 | K1 |
| 73 | 826.9403831 | 4 | -1.204445844 | [VHKTHSFPISLANTGK] | >CTF4 | 458 | 473 | [THSFPISLANTGK] | >CTF4 | 461 | 473 | K3 | T1 |
| 73 | 1123.953846 | 3 | 1.577914376 | [SIAPSIYELEDVKK] | >MCM4 | 525 | 538 | [SQILQYVHKITPR] | >MCM4 | 575 | 587 | K13 | K9 |
| 72 | 797.4091905 | 4 | -1.324069613 | [VHKTHSFPISLANTGK] | >CTF4 | 458 | 473 | [KPHNEHSYSR] | >CTF4 | 448 | 457 | S6 | K1 |
| 72 | 1081.899936 | 3 | -1.746408654 | [THSFPISLANTGKFR] | >CTF4 | 461 | 475 | [THSFPISLANTGK] | >CTF4 | 461 | 473 | K13 | S7 |
| 72 | 943.2201973 | 3 | 0.575804035 | [VIISEDFVQKSLSFLK] | >MRC1 | 908 | 923 | [LIKHLR] | >CDC45 | 518 | 523 | K10 | K3 |
| 72 | 642.9928553 | 3 | -1.453540378 | [KDDAQGFSR] | >MCM6 | 782 | 790 | [YKELR] | >MCM6 | 777 | 781 | K1 | K2 |
| 72 | 713.7354736 | 3 | -0.091071775 | [IQYEKTHPDR] | >PSF2 | 102 | 111 | [VLKGLK] | >PSF2 | 156 | 161 | K5 | K3 |
| 72 | 1047.0434 | 4 | 1.91962214 | [VAWHPKGLHFALPBADDTVK] | >CTF4 | 235 | 254 | [GYSLQKTLSTNLSSTK] | >CTF4 | 260 | 275 | K6 | K6 |
| 72 | 630.040498 | 3 | 0.773955102 | [KPQKPIPTK] | >MRC1 | 315 | 323 | [KFFSK] | >MRC1 | 324 | 328 | K4 | K1 |
| 72 | 680.7362991 | 3 | 0.954418879 | [ATILNLKAR] | >MRC1 | 439 | 447 | [LSKQNQK] | >MRC1 | 448 | 454 | K7 | S2 |
| 72 | 826.9410268 | 4 | -0.42536146 | [VHKTHSFPISLANTGK] | >CTF4 | 458 | 473 | [THSFPISLANTGK] | >CTF4 | 461 | 473 | K3 | T1 |
| 72 | 1395.723457 | 3 | 2.886747954 | [VAWHPKGLHFALPBADDTVK] | >CTF4 | 235 | 254 | [GYSLQKTLSTNLSSTK] | >CTF4 | 260 | 275 | K6 | Y2 |
| 72 | 1193.993324 | 3 | 3.55570753 | [IDMNLVQTGKSVIQR] | >MCM4 | 845 | 859 | [LQQEDKVIVLGEGVR] | >MCM4 | 910 | 924 | K10 | K6 |
| 71 | 617.0222427 | 3 | -1.080196669 | [SYKTVGSSK] | >MRC1 | 1033 | 1041 | [LIAPKR] | >MRC1 | 1053 | 1058 | K3 | K5 |
| 71 | 614.6941876 | 3 | -0.802570802 | [SKLLEENK] | >CTF4 | 784 | 791 | [AILNKK] | >CTF4 | 792 | 797 | S1 | K5 |
| 71 | 778.074646 | 3 | -1.040575523 | [YVKFNBLK] | >MCM2 | 336 | 343 | [VNGEKTVYR] | >MCM2 | 375 | 383 | K3 | K5 |
| 71 | 913.2097725 | 4 | -2.54398216 | [SBIELTHSVLKVLEQYSDDK] | >TOF1 | 590 | 609 | [SKIDNSADIK] | >TOF1 | 490 | 499 | K11 | S1 |
| 71 | 711.7255354 | 3 | -3.849482503 | [LSLWNENGKK] | >CDC45 | 348 | 357 | [TKIYPK] | >MCM2 | 772 | 777 | K9 | T1 |
| 71 | 652.7133507 | 3 | -3.167082072 | [TFKPILTK] | >MCM6 | 760 | 767 | [YIKYAR] | >MCM6 | 754 | 759 | K3 | K3 |
| 71 | 902.7315139 | 4 | -2.489628171 | [VHKTHSFPISLANTGK] | >CTF4 | 458 | 473 | [THSFPISLANTGKFR] | >CTF4 | 461 | 475 | K3 | T11 |
| 71 | 1211.605442 | 3 | -3.71096769 | [SKEDESPTTKIFTIIK] | >MCM7 | 734 | 749 | [EMDSKFSFGQATPR] | >MCM7 | 674 | 687 | K10 | S4 |
| 71 | 1008.170618 | 3 | -1.307650465 | [LGDDBLABLKDLK] | >TOF1 | 47 | 59 | [WFKLVDDQQK] | >TOF1 | 61 | 70 | K10 | K3 |
| 71 | 1047.06492 | 4 | 0.214943578 | [LYEILTNSIAPSIFGNEDIKK] | >MCM5 | 366 | 386 | [FVEKVSPIAVYTSGK] | >MCM5 | 428 | 442 | S12 | K4 |
| 71 | 659.0168105 | 3 | 0.160154053 | [EKADESLPKR] | >MRC1 | 617 | 626 | [SHHVK] | >MRC1 | 595 | 599 | K9 | S1 |
| 71 | 504.2876045 | 3 | 0.481087359 | [TTTYKK] | >MRC1 | 17 | 22 | [LEGKK] | >MRC1 | 67 | 71 | K5 | K4 |
| 71 | 1196.287516 | 3 | 1.633278022 | [LFASABSDQNVEKALSLAHELK] | >CTF4 | 869 | 890 | [SKLLEENK] | >CTF4 | 784 | 791 | K13 | S1 |
| 70 | 580.3317415 | 3 | -0.193285645 | [KSPITAR] | >MCM3 | 694 | 700 | [SPQKSPK] | >MCM3 | 761 | 767 | K1 | K4 |
| 70 | 567.3200846 | 3 | 0.43789634 | [LSKQNQK] | >MRC1 | 448 | 454 | [INEKR] | >MRC1 | 378 | 382 | K3 | K4 |
| 70 | 715.7595254 | 3 | -0.066057137 | [STVEGIKLR] | >MCM6 | 558 | 566 | [TFKPILTK] | >MCM6 | 760 | 767 | K7 | K3 |
| 70 | 778.0748368 | 3 | -0.79508279 | [YVKFNBLK] | >MCM2 | 336 | 343 | [VNGEKTVYR] | >MCM2 | 375 | 383 | K3 | K5 |
| 70 | 1395.749621 | 3 | -0.631152237 | [LYEILTNSIAPSIFGNEDIKK] | >MCM5 | 366 | 386 | [FVEKVSPIAVYTSGK] | >MCM5 | 428 | 442 | S12 | K4 |
| 70 | 919.7405197 | 4 | 4.738893676 | [EIDYKDDVLDVILNQR] | >MCM7 | 131 | 146 | [FGGLLSIQTPDKTR] | >TOF1 | 360 | 373 | Y4 | K12 |
| 70 | 913.2139589 | 4 | 2.04409844 | [SBIELTHSVLKVLEQYSDDK] | >TOF1 | 590 | 609 | [SKIDNSADIK] | >TOF1 | 490 | 499 | K11 | S1 |
| 70 | 708.1369393 | 4 | 0.062095208 | [ALYALEKHETIQLR] | >MCM5 | 751 | 764 | [FEQELKR] | >MCM5 | 714 | 720 | K7 | K6 |
| 70 | 532.0187908 | 3 | 0.248632122 | [TVKILK] | >MRC1 | 1027 | 1032 | [LIAPKR] | >MRC1 | 1053 | 1058 | K3 | K5 |
| 69 | 642.9952393 | 3 | 2.257895748 | [KDDAQGFSR] | >MCM6 | 782 | 790 | [YKELR] | >MCM6 | 777 | 781 | K1 | K2 |
| 69 | 751.0539189 | 3 | 0.490243374 | [LEEMEKEIYK] | >PSF3 | 169 | 178 | [LKBAPR] | >MCM5 | 608 | 613 | Y9 | K2 |
| 69 | 1123.9501 | 3 | -1.756542587 | [SIAPSIYELEDVKK] | >MCM4 | 525 | 538 | [SQILQYVHKITPR] | >MCM4 | 575 | 587 | Y7 | K9 |
| 69 | 1066.87263 | 3 | -2.27206666 | [KYVSQFSDYFLAR] | >TOF1 | 736 | 748 | [FRELEDDSIKK] | >TOF1 | 682 | 692 | K1 | K10 |
| 69 | 590.0041917 | 3 | 0.10087681 | [ILFNKAK] | >PSF2 | 125 | 131 | [WMFKK} | >PSF3 | 190 | 195 | K5 | K4 |
| 69 | 682.7435669 | 3 | 0.864805999 | [MLSLLFKK] | >CDC45 | 44 | 51 | [KVELLNR] | >CDC45 | 488 | 494 | K7 | K1 |
| 69 | 632.5687256 | 4 | -0.79161443 | [AKDDFHDPIHELR] | >PSF2 | 130 | 142 | [WMFKK} | >PSF3 | 190 | 195 | K2 | K4 |
| 69 | 897.4672013 | 4 | 1.349232357 | [LFASABSDQNVEKALSLAHELK] | >CTF4 | 869 | 890 | [SKLLEENK] | >CTF4 | 784 | 791 | K13 | S1 |
| 68 | 664.6658512 | 3 | 0.764519576 | [KLQEDLSR] | >MCM4 | 860 | 867 | [KMGDDSR] | >MCM4 | 779 | 785 | K1 | K1 |
| 68 | 897.1386714 | 3 | 0.440451162 | [EIQSELASEDSKR] | >MRC1 | 293 | 305 | [TLVIEGKSR] | >TOF1 | 610 | 618 | S8 | K7 |
| 68 | 854.1527104 | 3 | -0.832996289 | [ALSLAHELKQDR] | >CTF4 | 882 | 893 | [AELPSLVKK] | >CTF4 | 905 | 913 | K9 | K8 |
| 68 | 1196.282525 | 3 | -2.540650747 | [LFASABSDQNVEKALSLAHELK] | >CTF4 | 869 | 890 | [SKLLEENK] | >CTF4 | 784 | 791 | K13 | S1 |
| 68 | 1017.553698 | 4 | 1.910524057 | [KLSDEPSDIIPLFETAITQVAKR] | >MCM5 | 78 | 100 | [SFKNFILEFR] | >MCM5 | 30 | 39 | K22 | K3 |

| | | | | | | | | | | | | | |
|---|---|---|---|---|---|---|---|---|---|---|---|---|---|
| 68 | 614.3412086 | 3 | -1.970940857 | [TVNKVDAK] | >MCM3 | 720 | 727 | [SPQKSPK] | >MCM3 | 761 | 767 | K4 | K4 |
| 68 | 715.7589722 | 3 | -0.839762087 | [STVEGIKLR] | >MCM6 | 558 | 566 | [TFKPILTK] | >MCM6 | 760 | 767 | K7 | K3 |
| 68 | 895.744379 | 4 | 0.837099747 | [IDMNLVQTGKSVIQR] | >MCM4 | 845 | 859 | [LQQEDKVIVLGEGVR] | >MCM4 | 910 | 924 | S11 | K6 |
| 68 | 794.7806104 | 3 | 0.871467165 | [SQILQYVHKITPR] | >MCM4 | 575 | 587 | [VSDKR] | >MCM4 | 469 | 473 | K9 | K4 |
| 68 | 733.6556928 | 4 | 1.275606767 | {MYGDLGNKLVLEAK] | >PSF1 | 0 | 14 | [NILKEVSNLR] | >PSF1 | 39 | 48 | K8 | K4 |
| 67 | 889.1293958 | 3 | 2.146141886 | [FRELEDDSIKK] | >TOF1 | 682 | 692 | [SKIDNSADIK] | >TOF1 | 490 | 499 | K10 | S1 |
| 67 | 649.6815272 | 3 | 0.315629441 | [EKADESLPK] | >MRC1 | 617 | 625 | [KSHHVK] | >MRC1 | 594 | 599 | K2 | K1 |
| 67 | 657.3336524 | 3 | -0.072025878 | [SHESYKDTK] | >PSF3 | 180 | 188 | [EIYKK] | >PSF3 | 175 | 179 | S4 | K4 |
| 67 | 736.1417647 | 4 | -1.3463264 | [SLYKTYVDVVHVK] | >MCM4 | 455 | 467 | [BSFADKQVIK] | >MCM4 | 389 | 398 | T5 | K6 |
| 67 | 977.8702629 | 3 | -0.329721395 | {MYGDLGNKLVLEAK] | >PSF1 | 0 | 14 | [NILKEVSNLR] | >PSF1 | 39 | 48 | K8 | K4 |
| 67 | 884.6673705 | 4 | -0.714344371 | [TSEVRPELYKASFTBDMBR] | >MCM6 | 297 | 315 | [KVDDVTGEK] | >MCM6 | 102 | 110 | K10 | K1 |
| 67 | 977.8708114 | 3 | 0.231580192 | {MYGDLGNKLVLEAK] | >PSF1 | 0 | 14 | [NILKEVSNLR] | >PSF1 | 39 | 48 | K8 | K4 |
| 67 | 1395.722647 | 3 | 2.305640703 | [VAWHPKGLHFALPBADDTVK] | >CTF4 | 235 | 254 | [GYSLQKTLSTNLSSTK] | >CTF4 | 260 | 275 | K6 | Y2 |
| 66 | 1134.517488 | 3 | 0.192230696 | [SNNYEDFETDKELSR] | >MRC1 | 924 | 938 | [LHQMDMDKVSR] | >MCM2 | 778 | 788 | S14 | K8 |
| 66 | 574.6636963 | 3 | -1.785529556 | [EGNKTVK] | >MRC1 | 1023 | 1029 | [ILKSYK] | >MRC1 | 1030 | 1035 | K4 | K3 |
| 66 | 634.049372 | 3 | -2.252141682 | [KVELLNR] | >CDC45 | 488 | 494 | [ILFNKAK] | >PSF2 | 125 | 131 | K1 | K5 |
| 66 | 777.1028735 | 3 | -1.968544918 | [VPLLSSYANNLKR] | >MRC1 | 383 | 395 | [INEKR] | >MRC1 | 378 | 382 | K12 | K4 |
| 66 | 1395.718123 | 3 | -0.937189717 | [VAWHPKGLHFALPBADDTVK] | >CTF4 | 235 | 254 | [GYSLQKTLSTNLSSTK] | >CTF4 | 260 | 275 | K6 | S14 |
| 66 | 1193.988133 | 3 | -0.794396604 | [IDMNLVQTGKSVIQR] | >MCM4 | 845 | 859 | [LQQEDKVIVLGEGVR] | >MCM4 | 910 | 924 | S11 | K6 |
| 66 | 1201.64845 | 3 | -0.59532645 | [SVYTSGKASSAAGLTAAVVR] | >MCM6 | 595 | 614 | [SQFLKYVVGFAPR] | >MCM6 | 582 | 594 | S9 | K5 |
| 66 | 810.3884221 | 3 | -0.281159994 | [EMDSKFSFGQATPR] | >MCM7 | 674 | 687 | [QDSKR] | >MCM7 | 669 | 673 | T12 | K4 |
| 66 | 567.3196515 | 3 | -0.326322437 | [LSKQNQK] | >MRC1 | 448 | 454 | [INEKR] | >MRC1 | 378 | 382 | K3 | K4 |
| 66 | 913.212304 | 4 | 0.2303424 | [SBIELTHSVLKVLEQYSDDK] | >TOF1 | 590 | 609 | [SKIDNSADIK] | >TOF1 | 490 | 499 | K11 | S1 |
| 66 | 913.2122887 | 4 | 0.213664903 | [SBIELTHSVLKVLEQYSDDK] | >TOF1 | 590 | 609 | [SKIDNSADIK] | >TOF1 | 490 | 499 | K11 | S1 |
| 66 | 556.9820083 | 3 | -0.275709343 | [ILKSYK] | >MRC1 | 1030 | 1035 | [FKEGNK] | >MRC1 | 1021 | 1026 | K3 | K2 |
| 66 | 948.8501219 | 3 | -0.29478975 | [FVEKVSPIAVYTSGK] | >MCM5 | 428 | 442 | [NVNGKHSIR] | >MCM2 | 526 | 534 | K4 | S7 |
| 66 | 1008.173914 | 3 | 1.964316253 | [LGDDBLABLKDLK] | >TOF1 | 47 | 59 | [WFKLVDDQQK] | >TOF1 | 61 | 70 | K10 | K3 |
| 65 | 608.338547 | 3 | -0.575326042 | [EKEHEAK] | >MRC1 | 700 | 706 | [LQLKQK] | >MRC1 | 694 | 699 | K2 | K4 |
| 65 | 663.8763511 | 4 | 0.156834147 | [ALYALEKHETIQLR] | >MCM5 | 751 | 764 | [KSPITAR] | >MCM3 | 694 | 700 | K7 | K1 |
| 65 | 525.2983398 | 4 | -0.569176534 | [LSQRPNKSK] | >MRC1 | 455 | 463 | [LSKQNQK] | >MRC1 | 448 | 454 | K7 | K3 |
| 65 | 760.0670955 | 3 | 0.325881978 | [NILKEVSNLR] | >PSF1 | 39 | 48 | {MYGDLGNK] | >PSF1 | 0 | 8 | K4 | {0 |
| 65 | 1011.843117 | 3 | -2.130317612 | [KDANLDYYNFNPMGIK] | >CTF4 | 670 | 685 | [MPVFVKSK] | >CTF4 | 778 | 785 | K1 | K6 |
| 65 | 1008.171035 | 3 | -0.893734092 | [LGDDBLABLKDLK] | >TOF1 | 47 | 59 | [WFKLVDDQQK] | >TOF1 | 61 | 70 | K10 | K3 |
| 65 | 715.760178 | 3 | 0.846505755 | [STVEGIKLR] | >MCM6 | 558 | 566 | [TFKPILTK] | >MCM6 | 760 | 767 | K7 | T1 |
| 65 | 1201.650944 | 3 | 1.481787804 | [SVYTSGKASSAAGLTAAVVR] | >MCM6 | 595 | 614 | [SQFLKYVVGFAPR] | >MCM6 | 582 | 594 | K7 | S1 |
| 65 | 617.0221729 | 3 | -1.193444014 | [SYKTVGSSK] | >MRC1 | 1033 | 1041 | [LIAPKR] | >MRC1 | 1053 | 1058 | K3 | K5 |
| 64 | 965.8194895 | 3 | 0.97962526 | [KYVSQFSDYFLAR] | >TOF1 | 736 | 748 | [ELEDDSIKK] | >TOF1 | 684 | 692 | K1 | K8 |
| 64 | 630.040156 | 3 | 0.230469063 | [KPQKPIPTK] | >MRC1 | 315 | 323 | [KFFSK] | >MRC1 | 324 | 328 | K4 | S4 |
| 64 | 674.0499996 | 3 | -0.863595559 | [ASITYMGKTR] | >MRC1 | 1042 | 1051 | [LIAPKR] | >MRC1 | 1053 | 1058 | K8 | K5 |
| 64 | 1017.552731 | 4 | 0.959667699 | [KLSDEPSDIIPLFETAITQVAKR] | >MCM5 | 78 | 100 | [SFKNFILEFR] | >MCM5 | 30 | 39 | K22 | K3 |
| 64 | 819.4169818 | 3 | 1.084407025 | [ESLYQETNKSK] | >MCM7 | 725 | 735 | [KMLQETGK] | >MCM7 | 750 | 757 | S10 | K1 |
| 64 | 657.3338209 | 3 | 0.184468231 | [SHESYKDTK] | >PSF3 | 180 | 188 | [EIYKK] | >PSF3 | 175 | 179 | T8 | K4 |
| 64 | 777.1050333 | 3 | 0.813046647 | [VPLLSSYANNLKR] | >MRC1 | 383 | 395 | [INEKR] | >MRC1 | 378 | 382 | K12 | K4 |
| 64 | 682.7439276 | 3 | 1.393649686 | [MLSLLFKK] | >CDC45 | 44 | 51 | [KVELLNR] | >CDC45 | 488 | 494 | K7 | K1 |
| 64 | 1047.065546 | 4 | 0.813076361 | [LYEILTNSIAPSIFGNEDIKK] | >MCM5 | 366 | 386 | [FVEKVSPIAVYTSGK] | >MCM5 | 428 | 442 | S12 | K4 |
| 64 | 1081.902647 | 3 | 0.760611348 | [THSFPISLANTGKFR] | >CTF4 | 461 | 475 | [THSFPISLANTGK] | >CTF4 | 461 | 473 | K13 | S7 |
| 64 | 1395.724245 | 3 | 3.451506339 | [VAWHPKGLHFALPBADDTVK] | >CTF4 | 235 | 254 | [GYSLQKTLSTNLSSTK] | >CTF4 | 260 | 275 | K6 | Y2 |
| 64 | 1157.57367 | 3 | 0.444399438 | [KDANLDYYNFNPMGIK] | >CTF4 | 670 | 685 | [THSFPISLANTGK] | >CTF4 | 461 | 473 | K1 | T1 |
| 63 | 607.6588936 | 3 | 0.435604886 | [LVDDQQKR] | >TOF1 | 64 | 71 | [SFKSR] | >MCM6 | 91 | 95 | K7 | K3 |
| 63 | 614.3426036 | 3 | 0.302159693 | [TVNKVDAK] | >MCM3 | 720 | 727 | [SPQKSPK] | >MCM3 | 761 | 767 | K4 | K4 |
| 63 | 590.3459182 | 3 | -2.688585427 | [ATILNLKAR] | >MRC1 | 439 | 447 | [SKDPK] | >MRC1 | 462 | 466 | K7 | S1 |
| 63 | 948.8482668 | 3 | -2.251236388 | [FVEKVSPIAVYTSGK] | >MCM5 | 428 | 442 | [NVNGKHSIR] | >MCM2 | 526 | 534 | K4 | K5 |
| 63 | 895.7445624 | 4 | 1.041987544 | [IDMNLVQTGKSVIQR] | >MCM4 | 845 | 859 | [LQQEDKVIVLGEGVR] | >MCM4 | 910 | 924 | S11 | K6 |
| 63 | 777.1050595 | 3 | 0.846859344 | [VPLLSSYANNLKR] | >MRC1 | 383 | 395 | [INEKR] | >MRC1 | 378 | 382 | K12 | K4 |
| 63 | 967.5244853 | 3 | -1.366298749 | [FNPLLQAGAKLAK] | >MCM3 | 625 | 637 | [NKGNYNGTEIPK] | >MCM3 | 638 | 649 | K10 | K2 |
| 63 | 977.8726113 | 3 | 2.073421748 | {MYGDLGNKLVLEAK] | >PSF1 | 0 | 14 | [NILKEVSNLR] | >PSF1 | 39 | 48 | K8 | K4 |
| 63 | 600.9746309 | 3 | 0.393984297 | [NDDNTKK] | >MCM3 | 688 | 694 | [SPQKSPK] | >MCM3 | 761 | 767 | T5 | K4 |
| 63 | 526.9606648 | 3 | -2.293910118 | [LIAPKR] | >MRC1 | 1053 | 1058 | [TEGSHR] | >MRC1 | 1060 | 1065 | K5 | S4 |
| 63 | 810.3869157 | 3 | -2.14151376 | [EMDSKFSFGQATPR] | >MCM7 | 674 | 687 | [QDSKR] | >MCM7 | 669 | 673 | T12 | K4 |
| 63 | 948.8491449 | 3 | -1.325131456 | [FVEKVSPIAVYTSGK] | >MCM5 | 428 | 442 | [NVNGKHSIR] | >MCM2 | 526 | 534 | K4 | K5 |
| 63 | 895.7454242 | 4 | 2.004952643 | [IDMNLVQTGKSVIQR] | >MCM4 | 845 | 859 | [LQQEDKVIVLGEGVR] | >MCM4 | 910 | 924 | S11 | K6 |
| 62 | 764.175048 | 4 | -2.317873449 | [SKDPKVDHNVLLNTLR] | >MRC1 | 462 | 477 | [KQILDHQK] | >MRC1 | 482 | 489 | K5 | K1 |
| 62 | 943.8461483 | 3 | -0.657407419 | [ALYALEKHETIQLR] | >MCM5 | 751 | 764 | [FEQELKR] | >MCM5 | 714 | 720 | K7 | K6 |
| 62 | 832.7719244 | 3 | 1.272466629 | [VVVDSFVDAQKVSVR] | >MCM2 | 840 | 854 | [YDDLK] | >MCM5 | 530 | 534 | K11 | Y1 |
| 62 | 708.1350447 | 4 | -2.616250925 | [ALYALEKHETIQLR] | >MCM5 | 751 | 764 | [FEQELKR] | >MCM5 | 714 | 720 | T10 | K6 |
| 62 | 1048.260131 | 3 | -2.020901437 | [LSKDESVLPISQLSK] | >MRC1 | 424 | 438 | [TFKPILTKEAR] | >MCM6 | 760 | 770 | S2 | K8 |
| 62 | 1017.554225 | 4 | 2.428963899 | [KLSDEPSDIIPLFETAITQVAKR] | >MCM5 | 78 | 100 | [SFKNFILEFR] | >MCM5 | 30 | 39 | K22 | S1 |
| 62 | 817.1151589 | 3 | 0.508563374 | [KFNISEGDITK] | >TOF1 | 622 | 632 | [TLVIEGKSR] | >TOF1 | 610 | 618 | K1 | K7 |
| 62 | 650.6521117 | 3 | 0.374118862 | [SKEDESPTTK] | >MCM7 | 734 | 743 | [QDSKR] | >MCM7 | 669 | 673 | T8 | K4 |
| 62 | 704.051185 | 3 | 1.262120646 | [QNQKLSQRPNK] | >MRC1 | 451 | 461 | [SKDPK] | >MRC1 | 462 | 466 | K4 | S1 |
| 62 | 618.3638318 | 3 | 1.458854696 | [SYKTVGSSK] | >MRC1 | 1033 | 1041 | [TVKILK] | >MRC1 | 1027 | 1032 | K3 | K3 |
| 62 | 843.4542827 | 3 | 2.211981636 | [ITQGSSSTTHR] | >SLD5 | 24 | 34 | [ALTAAVKISER] | >CTF4 | 894 | 904 | S5 | K7 |
| 62 | 924.7474163 | 4 | 1.133013107 | [TSIHEAMEQQSISISKAGIVTTLQAR] | >MCM2 | 618 | 643 | [KDPITK] | >MCM2 | 582 | 587 | K16 | K1 |
| 62 | 913.2129248 | 4 | 0.91076026 | [SBIELTHSVLKVLEQYSDDK] | >TOF1 | 590 | 609 | [SKIDNSADIK] | >TOF1 | 490 | 499 | K11 | S1 |
| 61 | 704.0504807 | 3 | 0.260888288 | [QNQKLSQRPNK] | >MRC1 | 451 | 461 | [SKDPK] | >MRC1 | 462 | 466 | K4 | S1 |
| 61 | 473.5333275 | 4 | -0.462965832 | [KPQKPIPTK] | >MRC1 | 315 | 323 | [INEKR] | >MRC1 | 378 | 382 | K1 | K4 |
| 61 | 801.7719661 | 3 | -1.002758145 | [LLQKEHQMR] | >TOF1 | 338 | 346 | [NVIKHTSAR] | >TOF1 | 348 | 356 | K4 | K4 |

| | | | | | | | | | | | | | |
|---|---|---|---|---|---|---|---|---|---|---|---|---|---|
| 61 | 1102.26339 | 3 | 0.274820474 | [KQLLINELESTER] | >MCM5 | 631 | 643 | [NTLSYENIVKTVR] | >MCM7 | 758 | 770 | K1 | K10 |
| 61 | 777.1050479 | 3 | 0.831854853 | [VPLLSSYANNLKR] | >MRC1 | 383 | 395 | [INEKR] | >MRC1 | 378 | 382 | K12 | K4 |
| 61 | 924.7470202 | 4 | 0.70433273 | [TSIHEAMEQQSISISKAGIVTTLQAR] | >MCM2 | 618 | 643 | [KDPITK] | >MCM2 | 582 | 587 | K16 | K1 |
| 61 | 1201.649543 | 3 | 0.314840621 | [SVYTSGKASSAAGLTAAVVR] | >MCM6 | 595 | 614 | [SQFLKYVVGFAPR] | >MCM6 | 582 | 594 | K7 | S1 |
| 61 | 997.5680719 | 3 | 0.402114607 | [IDNSADIKQVSEALK] | >TOF1 | 492 | 506 | [KPQKPIPTKK] | >MRC1 | 315 | 324 | K8 | T8 |
| 61 | 1092.519914 | 3 | -1.201915948 | [SDAAYFKDLDNNASDK] | >TOF1 | 984 | 999 | [WFKLVDDQQK] | >TOF1 | 61 | 70 | K7 | K3 |
| 61 | 1018.883953 | 3 | -0.596044577 | [EKEIVENLLEQEILR] | >MRC1 | 505 | 519 | [GLKLEDMAK] | >MRC1 | 496 | 504 | K2 | K3 |
| 61 | 1157.116304 | 4 | 1.651744354 | [MSGGKETTDIHVWPLALAYDTLNBILVK] | >CTF4 | 731 | 758 | [LLEENKAILNK] | >CTF4 | 786 | 796 | S2 | K6 |
| 61 | 785.0808794 | 3 | -0.77318897 | [SKIDNSADIK] | >TOF1 | 490 | 499 | [KVFSFFHR] | >TOF1 | 692 | 699 | S1 | K1 |
| 61 | 1092.526664 | 3 | 4.980550778 | [SDAAYFKDLDNNASDK] | >TOF1 | 984 | 999 | [WFKLVDDQQK] | >TOF1 | 61 | 70 | K7 | K3 |
| 61 | 1458.729619 | 3 | 0.390585715 | [GKHIWPEFPLPLPSEMEIR] | >CTF4 | 759 | 777 | [KDANLDYYNFNPMGIK] | >CTF4 | 670 | 685 | K2 | K1 |
| 60 | 660.3783204 | 3 | -0.740975634 | [LLQKEHQMR] | >TOF1 | 338 | 346 | [KNVIK] | >TOF1 | 347 | 351 | K4 | K1 |
| 60 | 573.000694 | 3 | -1.117084409 | [TFKPILTK] | >MCM6 | 760 | 767 | [SKDPK] | >MRC1 | 462 | 466 | K3 | S1 |
| 60 | 965.1926157 | 3 | -1.262813226 | [FSQLALDKALYALEK] | >MCM5 | 743 | 757 | [KLQEDLSR] | >MCM4 | 860 | 867 | K8 | K1 |
| 60 | 1033.87608 | 3 | 0.020067017 | [FDLVYLVLDKVDEK] | >MCM4 | 702 | 715 | [SAIKDYATDPK] | >MCM4 | 831 | 841 | K10 | K4 |
| 60 | 682.7428453 | 3 | -0.193215374 | [MLSLLFKK] | >CDC45 | 44 | 51 | [KVELLNR] | >CDC45 | 488 | 494 | K7 | K1 |
| 60 | 1092.525068 | 3 | 3.518838826 | [SDAAYFKDLDNNASDK] | >TOF1 | 984 | 999 | [WFKLVDDQQK] | >TOF1 | 61 | 70 | K7 | K3 |
| 60 | 631.1365381 | 4 | 0.588485301 | [TYVDVVHVKK] | >MCM4 | 459 | 468 | [LINLKGLVLR] | >MCM4 | 325 | 334 | K9 | K5 |
| 60 | 680.7359119 | 3 | 0.385056673 | [ATILNLKAR] | >MRC1 | 439 | 447 | [LSKQNQK] | >MRC1 | 448 | 454 | K7 | S2 |

**Table S3. Summary of crosslinks identified in crosslinking mass spectrometry experiments.** The quality of the fragment ion assignment is measured by a scoring function (score) (Iacobucci et al., 2018). Briefly, all peptide pairs matching the identified reporter ions are subjected to scoring. The score is determined in relation to the presence and intensity of the DSBU reporter ions, the number and length of peptide backbone ion series and the number of identified ions related to the spectrum size and the number of possible fragment ions created from a theoretical peptide pair. To correct for random overlaps, features are also calculated for spectra with slightly shifted mass values (Iacobucci et al., 2018). Only peptides used in analysis with a score of ≥60 are shown.

**Vectors generated in this study**

| Name | Construction details | Application |
|---|---|---|
| vJY23 | pRS304-Psf1-Gal1-10-Sld5 | Psf1 / Sld5 |
| vJY24 | pRS306-Psf2-Gal1-10-His-Psf3 | Psf2 / His-Psf3 |
| vJY25 | pRS303-Fob1-TEV-2xFLAG- Gal1-10-Gal4 | Fob1 |
| vJY30 | A 243 bp fragment containing the yeast RFB from chromosome XII was amplified with vJY204 – TTCACTGTTCCTGCAGCACTTGTCTCTTACATCTTTCTTGG / vJY205-TTTTTTTTTTGGATCCGTTGCAAAGATGGGTTGAAAG and cloned into ZN5(Taylor and Yeeles, 2018) using BamHI and PstI. RFB sequence:<br>GTTGCAAAGATGGGTTGAAAGAGAAGGGCTTTCACAAAGCTTCCCGAGCGTG<br>AAAGGATTTGCCCGGACAGTTTGCTTCATGGAGCAGTTTTTTCCGCACCATCA<br>GAGCGGCAAACATGAGTGCTTGTATAAGTTTAGAGAATTGAGAAAAGCTCATT<br>TCCTATAGTTAACAGGACATGCCTTTGATATGAAAAAAAATACTACGAACTAC<br>GATTTTACCAAGAAAGATGTAAGAGACAAGTG | RFB template |
| vJY71 | pRS303-Cdc45$^{iF2}$-Gal1-10-Ctf4 | Cdc45$^{iF2}$ / Ctf4 |
| vJY72 | pRS based vector. Tof1-Gal1-10-Csm3 cloned Not1 / Apa1. Auxotrophic marker from pRS vector replaced with Nat-NT2 resistance marker amplified from pBP83(Yeeles et al., 2015) with oligos (JY247 - TCTAGTCAATGCGGCCGCCGTACGCTGCAGGTCGAC and JY248 - ATCGATGAATTCGAGCTCG) and cloned Zra1/Not1. Lys2 3321-3799 amplified with primers (JY253 – CCGATTGACAGGGCCCTTCCTCCAC TTCTACTCTTGACAC and JY254 – CCGATTGACAAAGCGGAAGAGCCGAC ATCGTAACCATAATCGTG) and cloned Sap1/Apa1. | Csm3 / Tof1 |
| vJY74 | pRS based vector. Mrc1-Gal1-10-Gal4 cloned Not1 / Apa1. Auxotrophic marker from pRS vector replaced with Nat-NT2 resistance marker amplified from pBP83(Yeeles et al., 2015) with oligos (JY247 and JY248) and cloned Zra1 / Not1. Lys2 3321-3799 amplified with primers (JY253 and JY254) and cloned Sap1/Apa1. | Mrc1 |
| vJY113 | pRS306 Tof1-gal1-10-CBP-Csm3$^{R49A/K53A}$<br>PCR mutagenesis with JY344 – AGTTGCTTTGACCGCTGAAAAGTTGTTG / 345 - TGTGGAGCTCTCTTTCTAGCGGTG | Csm3$^{R49A/K53A}$/Tof1 |
| vJY114 | pRS306 Tof1 $^{K400A/ R401A/ K404A}$-gal1-10-CBP-Csm3<br>PCR mutagenesis with JY347 - ATTGCTAAGCACCAATCCGTTGCTG / JY348 - AATAGCAGCGTTCCACTTCTTAGAATCATCC from pRS306/Tof1- Gal-CBP-Csm3 | Csm3/ Tof1$^{K400A/R401A/K404A}$ |
| vJY115 | pRS306 Tof1$^{K400A/R401A/K404A}$-gal1-10-CBP-Csm3$^{R49A/K53A}$<br>Subclone Csm3$^{R49A/K53A}$ into vJY114 with AscI/XhoI | Csm3$^{R49A/K53A}$/Tof1 $^{K400A/R401A/K404A}$ |
| vJY116 | pRS306 Tof1-gal1-10-CBP-Csm3 $^{K47A/ R48A/ R49A/ Q51A/ K53A}$<br>PCR mutagenesis with JY377 - GCTCCAGCTGTTGCTTTGACCG / JY378 - AGCAGCTCTAGCGGTGATAGCAGTTG from pRS306/Tof1- Gal-CBP-Csm3 | Csm3$^{K47A/R48A/R49A/Q51A/K53A}$/ Tof1 |
| vJY117 | pRS306 Tof1 $^{K400A/ R401A/ K404A}$-gal1-10-CBP-Csm3 $^{K47A/ R48A/ R49A/ Q51A/ K53A}$<br>PCR mutagenesis with JY377/JY378 from vJY114 | Csm3$^{K47A/R48A/R49A/Q51A/K53A}$/ Tof1$^{K400A/R401A/K404A}$ |
| vVA30 | The Tof1 open reading frame was amplified from W303 genomic DNA with primers vVA84 and vVA85 and was cloned into pFA6-Ura with AscI / SalI. | Parent vector for Tof1 mutagenesis |
| vVA31 | The Tof1-3A open reading frame was amplified with primers vVA84 and vVA85 and was cloned into pFA6-Ura with AscI / SalI. | Construction of Tof1-3A strains |
| vVA32 | The Csm3 open reading frame was amplified from W303 genomic DNA with primers vVA81 and vVA82 and was cloned into pFA6-Ura with AscI / SalI. | Parent vector for Tof1 mutagenesis |
| vJY136 | PCR mutagenesis form vVA32 with JY422 – GATCCTCAAGTAGATTTAACA GCCGAAAAAC and JY423 - GTCATCATCAGCTGTAATTGCGGTTGGATC | Construction of Csm3-5D strains |
| vJY137 | PCR mutagenesis form vVA32 with JY397 – CCTGCTGTAGCATTAACAGCCGA AAAACTACTCAG and 398 - CGCTGCAGCTCTAGCTGTAATTGCGGTTGG | Construction of Csm3-5A strains |

**Yeast strains used in this study**

| Strain | Genotype | Reference |
| --- | --- | --- |
| yJY36 | *MAT**α** ade2-1 ura3-1 his3-11,15 trp1-1 leu2-3,112 can1-100*<br>*bar1::Hyg*<br>*pep4::KanMX*<br>*ura3::URA3pRS306-Psf2/His-Psf3*<br>*trp1::TRP1pRS304Psf1/Sld5*<br>*his::HISpRS303Cdc45iFlag2/Gal4* | This study |
| yAM22 | *MAT**a** ade2-1 ura3-1 his3-11,15 trp1-1 leu2-3,112 can1-100*<br>*bar1::Hyg*<br>*pep4::KanMX*<br>*trp1::TRP1pRS304Mcm4, Mcm5*<br>*ura3::URA3pRS306/Mcm2, CBP-TEV Mcm3*<br>*leu2::LEU2pRS305/Mcm6, Mcm7* | (Zhou et al., 2017) |
| yJY37 | *MAT**a** / MAT**α** ade2-1 ura3-1 his3-11,15 trp1-1 leu2-3,112 can1-100*<br>*bar1::Hyg*<br>*pep4::KanMX*<br>*trp1::TRP1pRS304Mcm4, Mcm5*<br>*trp1::TRP1pRS304Psf1/Sld5 (vJY23)*<br>*ura3::URA3pRS306/Mcm2, CBP-TEV Mcm3*<br>*ura3::URA3pRS306-Psf2/His-Psf3 (vJY24)*<br>*leu2::LEU2pRS305/Mcm6, Mcm7*<br>*his::HISpRS303Cdc45iFlag2/Gal4* | This study |
| yJY39 | *MATa ade2-1 ura3-1 his3-11,15 trp1-1 leu2-3,112 can1-100*<br>*bar1::Hyg*<br>*pep4::KanMX*<br>*his3::HIS3pRS303-Fob1-TEV-2xFLAG- Gal1-10-Gal4* | This study |
| yJY72 | *MAT**α** ade2-1 ura3-1 his3-11,15 trp1-1 leu2-3,112 can1-100*<br>*bar1::Hyg*<br>*pep4::KanMX*<br>*ura3::URA3pRS306-Psf2/His-Psf3 (vJY23)*<br>*trp1::TRP1pRS304Psf1/Sld5 (vJY24)*<br>*his::HISpRS303Cdc45iFlag2/untagged Ctf4 (vJY71)*<br>*lys2::NAT-untagged Mrc1 (vJY74)* | This study |
| yJY69 | *MAT**a** ade2-1 ura3-1 his3-11,15 trp1-1 leu2-3,112 can1-100*<br>*bar1::Hyg*<br>*pep4::KanMX*<br>*trp1::TRP1pRS304Mcm4, Mcm5*<br>*ura3::URA3pRS306/Mcm2, CBP-TEV Mcm3*<br>*leu2::LEU2pRS305/Mcm6, Mcm7*<br>*lys2::NAT- untagged Csm3/Tof1 (vJY72)* | This study |
| yJY74 | *MAT**a** /MAT**α** ade2-1 ura3-1 his3-11,15 trp1-1 leu2-3,112 can1-100*<br>*bar1::Hyg*<br>*pep4::KanMX*<br>*trp1::TRP1pRS304Mcm4, Mcm5*<br>*trp1::TRP1pRS304Psf1/Sld5 (vJY23)*<br>*ura3::URA3pRS306/Mcm2, CBP-TEV Mcm3*<br>*ura3::URA3pRS306-Psf2/His-Psf3 (vJY24)*<br>*leu2::LEU2pRS305/Mcm6, Mcm7*<br>*his::HISpRS303Cdc45iFlag2/untagged Ctf4 (vJY71)*<br>*lys2::NAT-untagged Csm3/Tof1 (vJY72)*<br>*lys2::NAT-untagged Mrc1 (vJY74)* | This study |
| yJY110 | *MATalpha ade2-1 ura3-1 his3-11,15 trp1-1 leu2-3,112 can1-100*<br>*bar1::Hyg*<br>*pep4::KanMX*<br>*URA::Ura pRS306 Tof1-gal1-10-CBP-Csm3*$^{R49A/K53A}$ *(vJY113)* | This study |
| yJY112 | *MATalpha ade2-1 ura3-1 his3-11,15 trp1-1 leu2-3,112 can1-100*<br>*bar1::Hyg*<br>*pep4::KanMX*<br>*URA::Ura pRS306 Tof1*$^{K400A/R401A/K404A}$*-gal1-10-CBP-Csm3*$^{R49A/K53A}$ *(vJY115)* | This study |
| yJY115 | *MAT**a** ade2-1 ura3-1 his3-11,15 trp1-1 leu2-3,112 can1-100*<br>*bar1::Hyg*<br>*pep4::KanMX*<br>*URA::Ura pRS306 Tof1* $^{K400A/ R401A/ K404A}$*-gal1-10-CBP-Csm3 (vJY114)* | This study |

| | | |
|---|---|---|
| yJY120 | *MAT**a** ade2-1 ura3-1 his3-11,15 trp1-1 leu2-3,112 can1-100*<br>*bar1::Hyg*<br>*pep4::KanMX*<br>*URA::Ura pRS306 Tof1-gal1-10-CBP-Csm3* $^{K47A/\ R48A/\ R49A/\ Q51A/\ K53A}$ *(vJY116)* | This study |
| yJY121 | *MAT**a** ade2-1 ura3-1 his3-11,15 trp1-1 leu2-3,112 can1-100*<br>*bar1::Hyg*<br>*pep4::KanMX*<br>*URA::Ura pRS306 Tof1* $^{K400A\ /R401A/\ K404A}$*-gal1-10-CBP-Csm3* $^{K47A/\ R48A\ /R49A/\ Q51A/\ K53A}$<br>*(vJY117)* | This study |
| W303-1a | *MAT**a** ade2-1 ura3-1 his3-11,15 trp1-1 leu2-3,112 can1-100* | Lab strain constructed by R. Rothstein |
| yJY145 | *MAT**a** ade2-1 ura3-1 his3-11,15 trp1-1 leu2-3,112 can1-100*<br>*Csm3 (Ura3)* | This study |
| yVA57 | *MAT**a** ade2-1 ura3-1 his3-11,15 trp1-1 leu2-3,112 can1-100*<br>*Tof1-3A* | This study |
| yVA67 | *MAT**a** ade2-1 ura3-1 his3-11,15 trp1-1 leu2-3,112 can1-100*<br>*Csm3-5D (Ura3)* | This study |
| yVA68 | *MAT**a** ade2-1 ura3-1 his3-11,15 trp1-1 leu2-3,112 can1-100*<br>*Csm3-5A (Ura3)* | This study |
| yVA70 | *MAT**a** ade2-1 ura3-1 his3-11,15 trp1-1 leu2-3,112 can1-100*<br>*Tof1-3A*<br>*Csm3-5D (Ura3)* | This study |
| yVA73 | *MAT**a** ade2-1 ura3-1 his3-11,15 trp1-1 leu2-3,112 can1-100*<br>*Tof1-3A*<br>*Csm3-5A (Ura3)* | This study |
| yBH77 | *MAT**a** ade2-1 ura3-1 his3-11,15 trp1-1 leu2-3,112 can1-100*<br>*tof1Δ::hphNT* | (Hodgson et al., 2007) |

**Table S4. Details of vectors and yeast strains.**

**Movie S1. Comparison of MCM subunits in conformations 1 and 2.** Movie shows morphing between models adjusted to the cryo-EM density maps of the complex in conformation 1 and conformation 2. The large global repositioning of the MCM C-tier (lower) relative to the N-tier (upper, binding dsDNA) is clear. The map for conformation 2 was that with 5 AMP-PNP molecules bound. To generate the movie, models for each conformation were aligned on MCM N-tier regions using PyMOL, before morphing between these models using Chimera.

**Movie S2. Overview of MCM N-tier interactions with DNA at the fork junction with the residues involved in the direct interaction highlighted.** Details of the Mcm7 NTH strand-separation pin are shown at the end. Movie produced using PyMOL.

**Movie S3. Overview of the structure of Csm3/Tof1 demonstrating the positions of the Tof1 Ω-loop, MCM-plugin and Csm3-binding element and showing the binding of Csm3 to Tof1.** Movie produced using PyMOL.

**Movie S4. Overview of the interface formed between Csm3/Tof1 and MCM.** Movie produced using PyMOL.

**Movie S5. Overview of the interactions between Csm3/Tof1 and the parental dsDNA duplex.** Movie produced using PyMOL.
